# Beyond benchmarking: an expert-guided consensus approach to spatially aware clustering

**DOI:** 10.1101/2025.06.23.660861

**Authors:** Jieran Sun, Kirti Biharie, Peiying Cai, Niklas Müller-Bötticher, Paul Kiessling, Meghan A. Turner, Søren H. Dam, Florian Heyl, Sarusan Kathirchelvan, Martin Emons, Samuel Gunz, Sven Twardziok, Amin El-Heliebi, Martin Zacharias, SpaceHack 2.0 participants, Roland Eils, Marcel Reinders, Raphael Gottardo, Christoph Kuppe, Brian Long, Ahmed Mahfouz, Mark D. Robinson, Naveed Ishaque

## Abstract

Spatial omics technologies have revolutionized the study of tissue architecture and cellular heterogeneity by integrating molecular profiles with spatial localization. In spatially resolved transcriptomics, delineating higher-order anatomical structures is critical for understanding how cellular organization affects tissue and organ function. Since 2020, more than 50 spatially aware clustering (SAC) methods have been developed for this purpose. However, the reliability of current benchmarks is undermined by their narrow focus on Visium and brain tissue datasets, as well as incorrect interpretation of manual annotation as ground truth. Here, we present SACCELERATOR, a community-driven, extensible framework that standardizes data formatting, method integration, and metric evaluation, and is designed to rapidly incorporate new methods and datasets. SACCELERATOR currently includes 22 SAC methods applied to 15 datasets spanning 9 technologies and diverse tissue types. Our analysis revealed substantial limitations in the generalizability and reproducibility of SAC methods across tissues and platforms. We also demonstrate that anatomical labels commonly used as ground truths are often biased, potentially error-prone, and, in some cases, unsuitable for benchmarking efforts.

Rather than scoring and comparing methods, we propose a consensus-guided workflow that aggregates clustering results to generate consensus representations. Descriptive spatial metrics highlight areas of high entropy where method disagreement is highest, enabling targeted feedback for tissue experts. Applied to brain and cancer datasets, this approach uncovered biologically meaningful patterns overlooked by individual methods and manual annotations. Our results underscore the need for iterative, expert-in-the-loop analysis and reveal that traditional evaluation metrics do not always capture the subjective qualities of results. By improving tissue annotation and addressing key benchmarking limitations, SACCELERATOR provides a robust foundation for advancing spatial omics research.

## Introduction

Spatially resolved transcriptomics (SRT) technologies map gene expression while preserving the spatial architecture of tissues, offering insights into how cells are organized into higher-order anatomical structures, such as tissues and organs [1, 2]. Delineating higher-order anatomical structures is pivotal for advancing our understanding of biological systems. These structures reveal how cells communicate within and between tissues, offering new perspectives on the mechanisms underlying both healthy and diseased states. Such insights are critical for identifying the origins of tumor-suppressing or -promoting microenvironments, discovering biomarkers, and informing the development of targeted therapies [3]. In the brain, the intricate cellular relationships between anatomical structures, cell types, and functional circuits remain poorly understood. Leveraging SRT data to map these relationships lays the groundwork for developing comprehensive and multimodal brain atlases, bridging molecular and functional insights at unprecedented resolution [4].

Since 2020, more than 50 spatially aware clustering (SAC) methods have been developed to delineate the structures in SRT data into spatial regions. However, terminology describing these spatial regions remains inconsistent across studies. In some studies, “spatial domains” or “spatial clusters” are loosely defined as tissue regions exhibiting spatially coherent gene expression patterns [5]. Others characterize domains as unique local features with distinct cell-type compositions, transcriptomic heterogeneity, and cell-cell interactions [6]. Yet others use the terms “spatially homogeneous regions” [7], “cellular neighborhoods” [3], “schematics” [8] or “niches” [9] to describe regions of homogeneous cell-type composition and/or gene expression. However, the term “niche” has also been used to describe a local structure in tissues that can reside within a tissue region [10]. Additionally, “milieu” refers to the immediate environment surrounding a cell, encompassing its direct neighbors, extracellular fluids, and the extracellular matrix [11]. This lack of standardized terminology reflects the diversity of biological questions and spatial scales addressed by SRT, ranging from individual cells [12] to organs [13], which further complicates method comparison and downstream analysis. It is important to distinguish between (i) the well-established anatomical structures (e.g., “Functional Tissue Units” [14] or parcellations [15]), (ii) the computationally identified features, and (iii) the modeling approaches used. SAC methods typically integrate gene expression and spatial information through feature engineering, dimensionality reduction, and clustering. However, the resulting clusters may not always align with biological structures of interest, highlighting the need for clarity in definitions and expectations.

Robust benchmarking is required to guide method selection and parameterization. Previous studies have compared SAC methods across various settings, including SRT technologies, tissues and simulated datasets, pre-processing, and random seed initializations [5, 16–18]. However, these studies are not easily extensible to new methods or datasets, and are limited in scope, focusing primarily on brain datasets generated by early technologies such as Visium.

The most common approach to benchmark SAC methods is to compare their output with ground truth (GT) annotations. Traditionally, these GT annotations are manually annotated based on apparent structures and regions using histological stains such as hematoxylin and eosin stain (H&E), often augmented with tissue-specific labels to visualize cytoarchitecture, pathology, or other features of interest [19]. Because GT annotations are commonly determined manually by experts, they are prone to human error and knowledge biases. The lack of well-defined ontologies describing anatomical structures in tissues hinders accurate and standardized definition of GTs, although, efforts are underway to define cell-type ontologies using single-cell and SRT technologies [20–23].

The challenges of benchmarking SAC methods became apparent during the BioHackathon Germany SpaceHack 2.0 project [24]. Our initial goal was to implement an extensible, community-driven benchmark to quantitatively evaluate the performance of SAC methods across different SRT technologies and tissues, but we encountered three main obstacles. First, most SAC methods were not sufficiently demonstrated on modern SRT technologies. Second, the conceptual complexity in defining GTs rendered standard performance metrics difficult to interpret. Third, in most real-world scenarios, SAC methods are utilized in an interactive and iterative way, giving more importance to being able to investigate the output of different methods and parameters.

To address these challenges, we propose a consensus-guided, expert-in-the-loop framework to systematically evaluate commonalities and differences between methods and parameter choices. This framework enhances tissue annotations, facilitates method recommendations, and reflects the iterative, collaborative nature of spatial omics research.

In this study, we present evidence of inconsistencies in previously reported method performances and demonstrate the value of leveraging consensus among SAC results. We also highlight the need for improved computational efficiency in SAC methods to accommodate modern large-scale SRT datasets. Finally, we present two case studies to illustrate the utility of method consensus and entropy in expert-guided analysis. Our extensible framework, SACCELERATOR, is available on GitHub (see Data and code availability) with clear contribution guidelines and templates to enable the research community to reuse and extend the existing implemented datasets, methods, and metrics investigated in this study.

## Results

### Reported performance of spatially aware clustering methods is inconsistent across studies

We observed inconsistencies in reported method performances across existing SAC benchmarks. To evaluate this systematically, we conducted a meta-analysis of 13 studies [9, 12, 16, 25–34], using the widely-used Visium dataset of the human brain dorsolateral prefrontal cortex (LIBD DLPFC) region [35].

We focused on BayesSpace, a foundational SAC method, as a benchmark control. We expected relatively consistent BayesSpace results across studies; however, collected adjusted Rand indices (ARI) revealed substantial variability (Figure 1A). For instance, slice 151673 showed strong and consistent performance (and is often showcased in the literature [12, 25–29]), whereas the results of slice 151669 fluctuated dramatically, highlighting the lack of robustness of reported method performances across previous studies.

**Figure 1:**
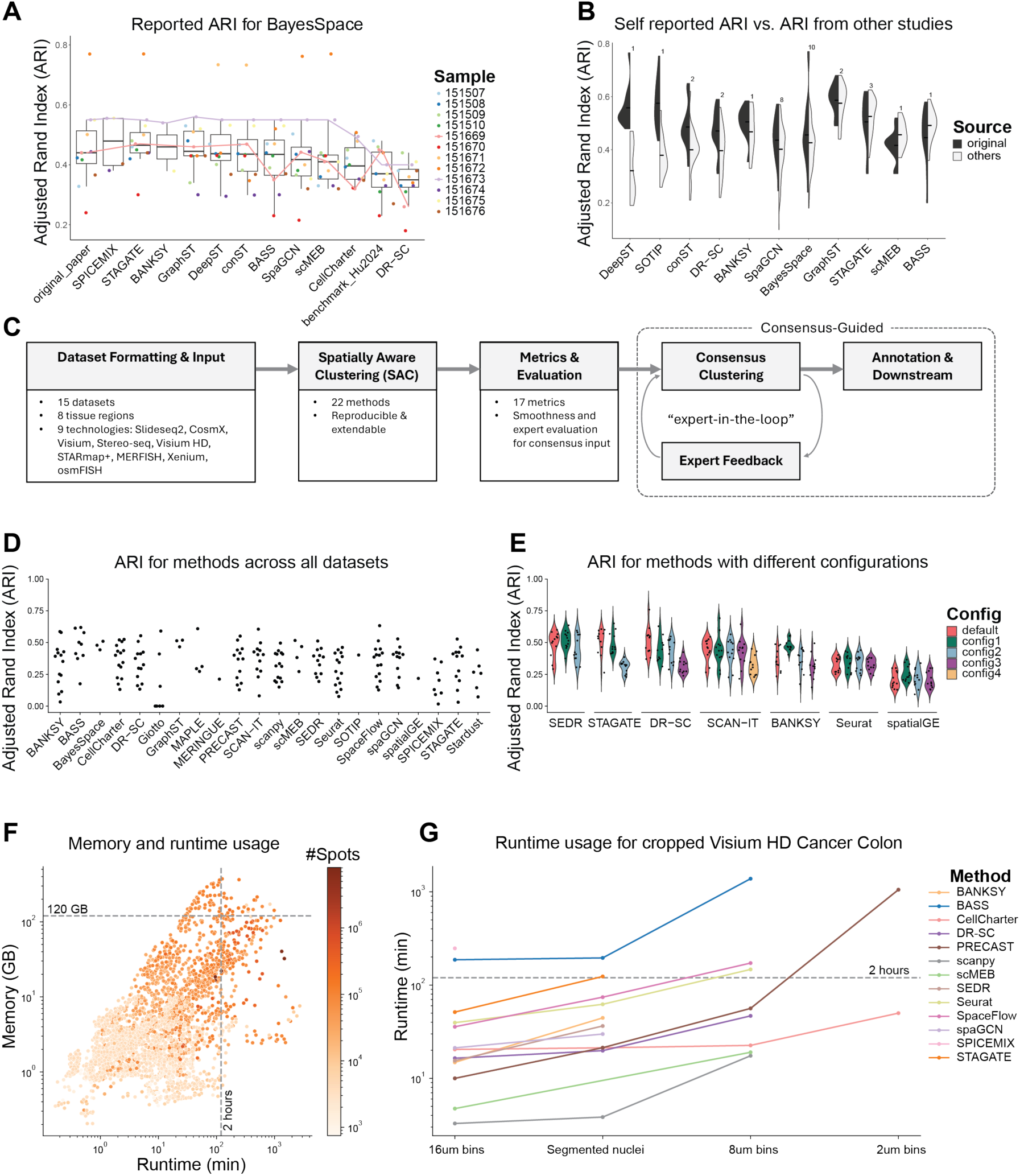
Addressing inconsistent reporting of method performance. A. Reported Adjusted Rand Index (ARI) of BayesSpace on the Visium dataset of the human dorsolateral prefrontal cortex (LIBD DLPFC), in the original BayesSpace study and other studies. The reported ARI for each sample in the dataset is depicted in a different color, with those for slices 151669 and 151673 connected with a line. Studies after BayesSpace are ordered by decreasing median ARI. B. The original self-reported ARI (dark grey) vs the ARI reported in other studies (light grey) for method performance on the LIBD DLPFC dataset. Methods are arranged by the difference of mean of self-reported ARI vs those in other studies. C. Schematic of the SACCELERATOR framework. D. ARI for methods across different datasets. E. ARI for selected methods with multiple configurations applied to the LIBD DLPFC dataset. F. Memory usage vs CPU runtime across SRT technologies, colored by the number of spots/cells in those datasets. G. CPU runtime of methods on 25% of the Visium HD colorectal cancer dataset across various bin sizes.

As expected, we observed that performances reported in method papers are frequently higher than those reported by subsequent competitors (Figure 1B). This discrepancy could arise from the original studies optimizing their methods to demonstrate superior performance, or suboptimal parameter choices in subsequent comparisons. While neutral benchmarking studies have aimed to systematically address such biases, we still encountered inconsistencies. For example, BayesSpace was ranked as 4^th^ out of 14 methods in Yuan et al. [5] but 13^th^ out of 16 methods in Hu et al. [16] based on the same LIBD DLPFC dataset. Taken together, this meta-analysis questions the utility of conventional SAC benchmarks.

### An extensible framework for running and evaluating SAC methods

Given the limitations of existing benchmarking efforts, we investigated the utility of SAC methods across various real-world scenarios. To achieve this, we developed SACCELERATOR, a modular and reproducible workflow built on Snakemake [36], specifically designed for straightforward extension with new datasets, methods, and evaluation metrics (Figure 1C). Within this framework, each dataset, method, and metric is encapsulated as an independent module, with standardized command-line interfaces and outputs. This modular structure enables seamless integration of additional methods or datasets, and method-specific configuration files to facilitate systematic exploration of parameter spaces.

Recognizing that SRT technologies produce data in diverse formats, we defined a set of necessary and optional metadata for each dataset and systematically stored it with the SRT data in generic file formats (TSV, JSON). This design ensures that downstream analyses are agnostic to the underlying technology and that data can be easily accessed across different programming languages.

The SACCELERATOR framework enables performance evaluation via both spatially agnostic clustering metrics, such as ARI or normalized mutual information (NMI), as well as spatially aware metrics, such as spatial chaos score (CHAOS)[6], percentage of abnormal spots (PAS) score [6], or spot entropy [37]. These spatially aware metrics consider the spatial location of the predicted labels and/or evaluate the spatial continuity of the clusters [6, 37, 38]. Moreover, the clustering results of different parameter combinations or multiple methods can be aggregated and leveraged for analyses using consensus clustering approaches (more on this below).

Our current implementation includes data from over 170 tissue samples from 15 datasets, generated using the sequencing-based SRT technologies Visium [39], Visium HD [40], Slide-seqV2 [41], and Stereo-seq as well as the imaging-based technologies Xenium, CosMx, STARmap PLUS [42], and MERFISH [43] (See Methods, Table 1 for dataset summaries). The framework applied the 22 implemented methods (Supplementary Table S1) to these datasets, each with one to five parameter configurations (61 total combinations; Supplementary Table S2). The method outputs were evaluated using multiple metrics that used the GT labels and spatial information to varying extents (Supplementary Table S3). In general, we observed substantial variation in SAC method performance as measured by ARI (Figure 1D). In the workflow, we have implemented both accuracy-based metrics, such as ARI and NMI, and spatial continuity-based metrics, such as smoothness entropy (SE) and CHAOS. Across different datasets, the two groups of metrics show general concordance among themselves in terms of correlation and rankings (Supplementary Figure S1 to S2).

**Table 1:**
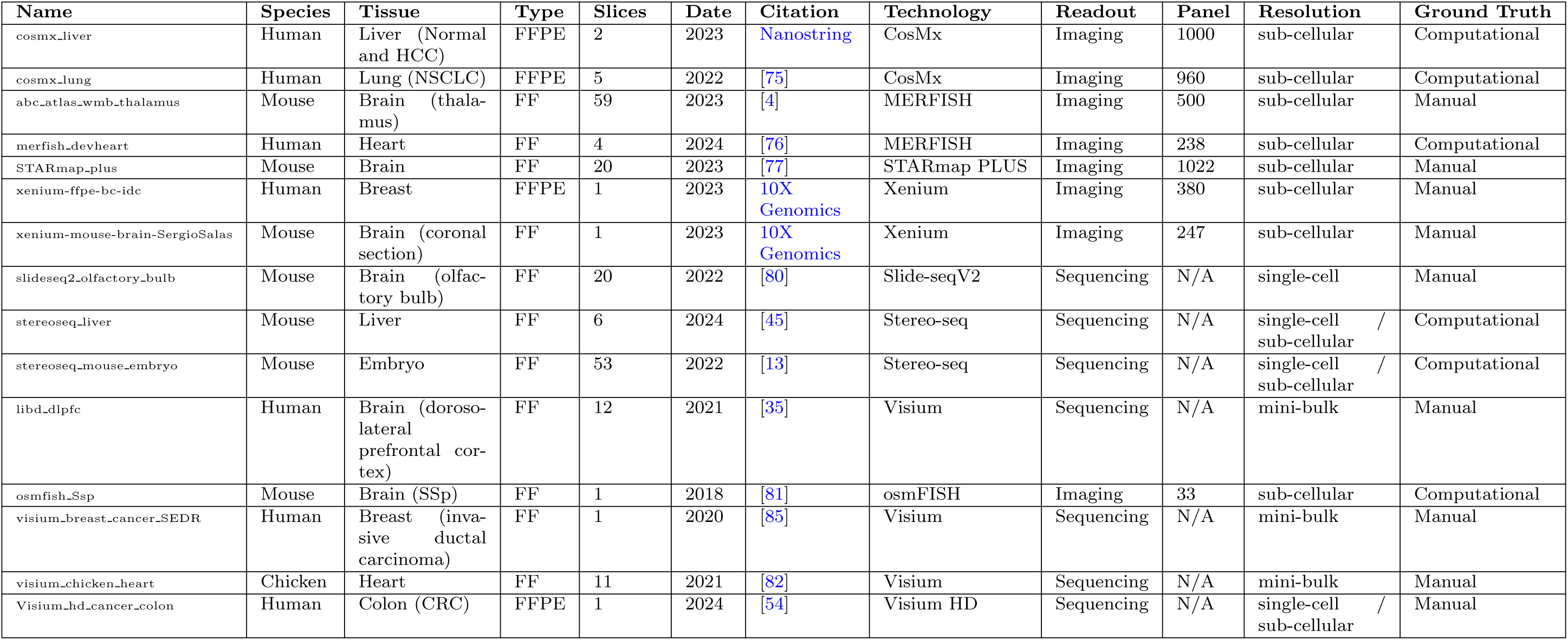
An overview of the datasets used in this study. FF = fresh frozen. FFPE = formalin-fixed paraffin-embedded.

Method-specific parameter choices, such as the number of principal components or the degree of smoothing, can contribute to performance variability. To assess the impact of parameterization, we evaluated each method using both default settings and parameters recommended in tutorials or dataset-specific analyses after uniform application of quality control and filtering steps (see Methods). Our results indicate that differences between parameter configurations can exceed those between different methods (Figure 1E, Supplementary Figure S3). Some methods exhibited stable performance across configurations (e.g., SEDR), while others (e.g., STAGATE) achieved results ranging from top- to bottom-tier performance, likely reflecting sensitivity to technology-specific parameters (Figure 1E). This pronounced sensitivity underscores the challenge for users in selecting optimal settings and highlights the need for clear guidance and robust defaults.

As a community-driven resource, SACCELERATOR is designed for ongoing expansion, and we encourage users and developers to contribute additional datasets and methods (see Data and code availability for a link to the framework and guidelines).

### Few current methods scale to modern datasets

Newer SRT technologies have improved spatial resolution and/or gene plexity, leading to a substantial increase in data volume. The smallest dataset we evaluated contained fewer than 3,500 spots (Visium LIBD DLPFC), while the largest dataset contained more than 460,000 cells (CosMx liver cancer). To assess whether SAC methods scale to large datasets, we applied them to large Visium HD, Stereo-seq, Xenium, and CosMx datasets. We measured the CPU runtime and memory usage of the methods with respect to the number of spatial observations (i.e., cells/bins/spots) and features (genes). We found that both runtime and memory usage increased exponentially with the number of spatial observations (Figure 1F), whereas the number of features had little impact, likely due to the widespread use of feature selection and/or dimensionality reduction steps in most SAC methods (Supplementary Figure S4A). Similarly, the specific SRT technology had less impact on runtime and memory usage than the number of spatial observations (Supplementary Figure S5). Most methods scaled well until 200,000 spatial observations (Figure 1F), but only 5 of 22 methods were able to process the largest dataset (CosMx liver cancer) on a compute node with 500 GB RAM and 10 cores within 2 days.

To further explore method scalability, we examined resource usage for the Visium HD colorectal cancer dataset across multiple bin-sizes (2, 8, 16 µm, resulting in 8, 0.5, and 0.125 million bins, respectively; Supplementary Figure S4B). While 11 out of the 22completed for the 16 µm bins, only 5 could process the 8 µm bins, and just CellCharter managed to process the 2 µm bins within the memory and runtime constraints; all other methods exceeded the 500 GB memory limit. Since only one method completed for all bin sizes, we repeated the experiment using a cropped 25% subset of the dataset (Figure 1G). This increased the number of methods able to process the 16 µm bins to 13 (of 22), and 8 successfully finishing for the 8 µm bins. For the smallest bin size, only PRECAST could now successfully run in addition to CellCharter. Of these two, CellCharter demonstrated both lower memory usage and faster runtime. When comparing the resulting clusters to each other and to the ground truth, the two methods showed low similarity (ARI = 0.10), with CellCharter being more similar to the ground truth than PRECAST (ARIs = 0.31 vs 0.07; Supplementary Figure S6).

Taken together, these results demonstrate that most SAC methods do not currently scale to state-of-theart SRT technologies and may not be usable for large-scale analyses, such as those involving large cohorts or extensive tissue sections.

### Over-reliance on ground truth annotations complicates method evaluations

One of the principal limitations affecting the validity of benchmarking studies for SAC methods is the reliance on GT labels. GT labels are often derived from multi-modal information, which may not be fully reflected in the transcriptome alone. For example, in the Visium LIBD DLPFC dataset [35], GT annotations were generated by cross-referencing histological H&E staining images (Figure 2A) with marker gene expression. Except for white matter and, to a lesser extent, layer 6, the brain isocortex layers in this dataset were grouped closely in the first two principal components based solely on transcriptional profiles (Figure 2B). Similar intermixed patterns can also be observed in other datasets (Supplementary Figure S7).

**Figure 2:**
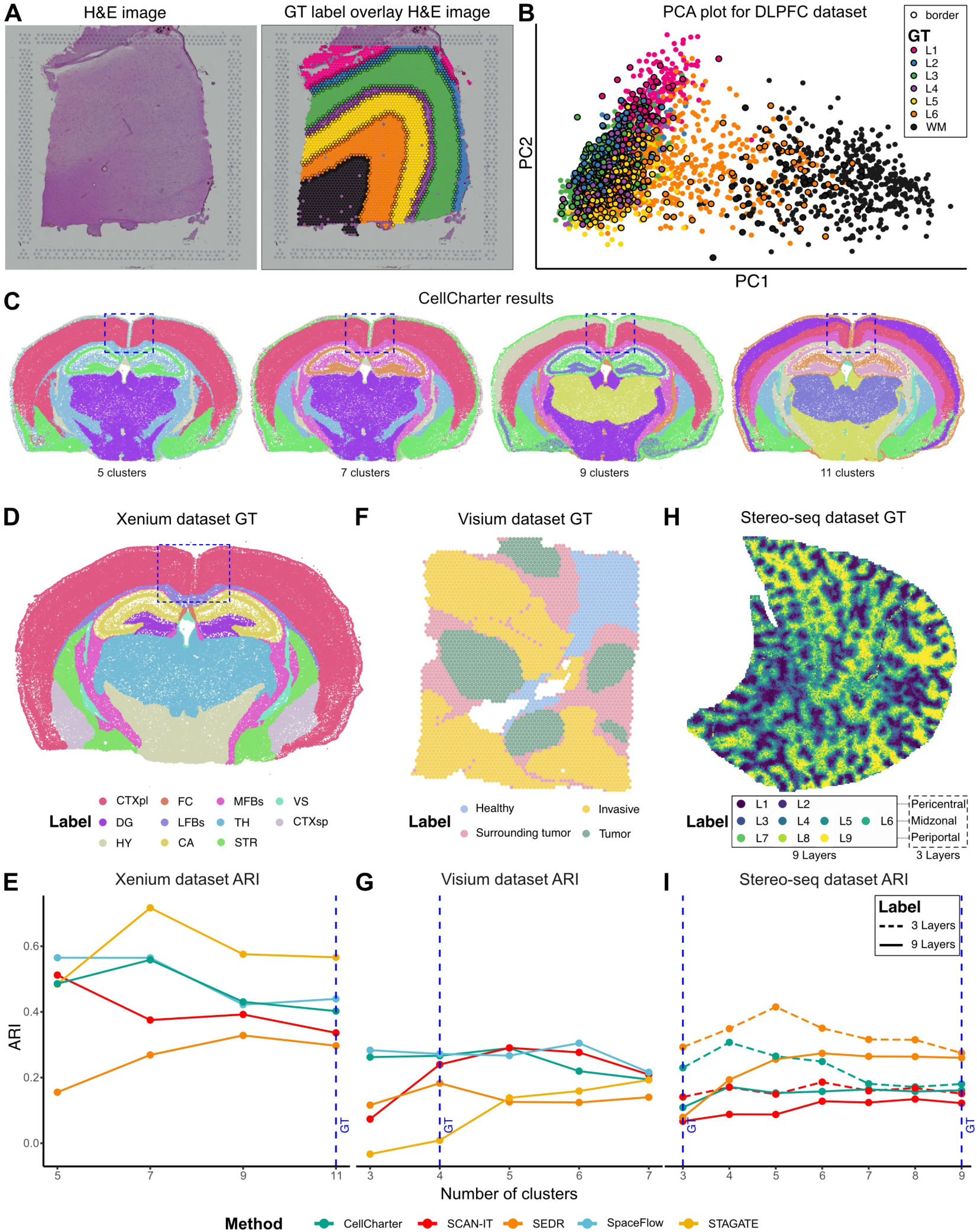
Challenges in utilizing ground truths effectively for benchmarking. A. H&E images of dorsolateral prefrontal cortex from the Visium LIBD DLPFC Br8100 151673 slice (left), and the ground truth (GT) annotation of the same sample overlaying on the H&E image (right). B. LIBD DLPFC Br8100 151673 slice, projected on the 1^st^ and 2^nd^ principal component (PC), colored by GT labels. Spots that are spatially located at the border between GT clusters are marked with black outlines. C. The results of CellCharter on the same sample while setting the number of clusters to 5, 7, 9, and 11. D, F, H. Spatial plots of the GT labels of Xenium mouse brain dataset (D), Visium Breast Cancer dataset (F), and Stereo-seq mouse liver dataset (H). E, G, I. ARI against the GT labels of several methods when set to identify varying numbers of clusters for the corresponding Xenium (E), Visium (G), and Stereo-seq (I) datasets. The cluster number of the GT labels is marked by dashed blue lines. Two sets of GT labels are shown in the Stereo-seq dataset (I), marked by different line types. GT: Ground truth, ARI: Adjusted Rand Index.

Gene expression differences across the isocortex layers were subtle and could be further obscured by technical limitations, such as coarse resolution (multiple cells per spot), transcript-capture biases (e.g., lateral diffusion), and other technical limitations of sequencing-based SRT technologies [44]. Beyond these technical challenges, there may be conceptual misalignments between the identified spatial clusters and the biological reality of tissue organization. Many biological phenomena, including transcriptomic variations, gradientdriven expression patterns (see thalamus case study below), and transitional cell stages, are continuous and diffusive processes, whereas spatial cluster detection inherently discretizes this landscape.

The assignment of GT labels is also inherently subjective, typically reflecting the annotator’s biological interests and the scale of their investigation. For instance, in the Xenium mouse brain datasets (Datasets, Figure 2C), the entire isocortex is labeled as a single region in the GT (Figure 2D; highlighted in box). However, when methods were configured to identify the same number of clusters as the GT labels, some methods detected more refined, layered structures within the isocortex (Figure 2C; 11 clusters, Supplementary Figure S8). Although these layered structures in the isocortex are biologically meaningful, they diverge from the GT label delineations and result in lower ARI scores. Conversely, configuring methods to find fewer clusters (Figure 2C; 7 and 9 clusters) produces a less granular isocortex structure that better matches the GT labels and yields higher ARI scores (Figure 2E; green). However, this reduction in granularity can obscure other important anatomical features, such as the distinction between thalamus and hypothalamus, which is only captured at higher cluster numbers. As a result, the subjectivity of GT labels often penalizes alternative clustering strategies and induces granularity bias where methods often achieve peak ARI against the GT at cluster counts different from that of the GT labels (Figure 2E, G, I). This phenomenon is pervasive across datasets and methods.

The challenges of using GT labels to evaluate SAC methods stem from both subjective granularity and divergent annotation objectives. For example, the GT annotation for the Visium breast cancer dataset [45] (see Datasets, Figure 2F) consists of four manually defined regions: tumor, tumor-surrounding, invasive, and healthy tissue. While these labels provide a biologically motivated scaffold, their coarse granularity introduces evaluation biases, where methods such as STAGATE and SCAN-IT achieve their highest ARIs only when configured to identify more clusters than those defined in the GT label (Figure 2G). More fundamentally, there exists an intrinsic mismatch: GT labels are annotated to capture broad pathological zones based on visual histological features, whereas most SAC methods are optimized to identify transcriptionally coherent spatial patterns. This conceptual disconnect leads to uniformly low ARI scores across all methods (mean ARI = 0.20), even when the resulting clusters visually correspond to meaningful tissue structures (Supplementary Figure S9).

In the Stereo-seq mouse liver dataset [45] (Figure 2H, see Datasets), the GT labels were annotated to reflect hepatocyte zonation patterns between central and portal veins in liver lobules [46]. While zonation patterns represent physiologically relevant processes, most SAC methods achieved a low ARI score for this dataset (Figure 2I, Supplementary Figure S10). This low ARI could potentially be attributed to several factors, including the use of supervised gene selection for GT annotation or the granularity of the annotated zones.

To assess the impact of expert gene selection, we identified 925 zonation-associated gene sets from three studies [47–49] that were not used during the original annotation of the Stereo-seq mouse liver dataset. This had mixed effects on the results, with SpaceFlow achieving high ARI and SEDR and CellCharter reporting lower ARI compared to their performance based on all genes (Supplementary Figure S11).

We further investigated the effect of annotation granularity. The original GT divided the zonation pattern into nine equal parts, but these could also be grouped into three biologically established layers: pericentral, midzonal, and periportal. When using this three-layer level of granularity, ARI scores improved markedly compared to the original nine-layer GT (Figure 2H, I), highlighting the importance of matching the granularity of GT labels to the biological structures of interest. This also suggests that some SAC methods may benefit from leveraging expertly defined gene sets rather than relying solely on unsupervised gene selection.

Since manually annotating GT labels is a challenging and time-consuming process, some datasets have used computationally generated GTs (Table 1). For example, in the Stereo-seq mouse embryo dataset, GT labels were derived using Spatially Constrained Clustering, a method building on the Leiden algorithm [13]. This methodological overlap likely explains why Leiden clustering implementations in Seurat and Scanpy achieved the highest mean ARI scores (0.428 and 0.421, respectively; Supplementary Figure S12). The use of computationally determined GTs introduces circularity, which is also evident in SAC benchmarking studies of the STARmap mouse cortex dataset. BASS, the method used to generate GT labels, consistently achieved top-tier performance in neutral benchmarks [5, 16].

Collectively, these challenges, such as discretization of continuous processes, granularity bias, circularity, and technical artifacts, are pervasive across current GT labels used in SAC benchmarking. While careful curation may mitigate some issues, the inherent limitations of GT labels still undermine their reliability, and consequently, the validity of current benchmarking practices.

### Methods are often more similar to each other than to the ground truth

Given the challenges in effectively utilizing GT references, we explored the similarity of clustering results produced by different SAC methods. We compared “base clusterings” (i.e., the set of clustering results from multiple configurations of multiple methods) to each other and also computed SE (see Methods) as an intrinsic, GT-independent performance measure, since some level of spatial continuity of clusters is expected. For the Visium LIBD DLPFC dataset, base clusterings revealed a wide range of smoothness levels (Figure 3A, B; Supplementary Figure S13). SAC methods often exhibit higher concordance with each other than to the GT (Figure 3B). Notably, for slice 151673 of the LIBD DLPFC dataset, we identified a block of base clusterings with method-vs-method ARIs averaging around 0.56 (Figure 3B, blue box), and another block of methods exceeding 0.62 (Figure 3B, pink box with dashed lines). This pattern of method concordance was consistent across several algorithms and all slices of the LIBD DLPFC dataset (Supplementary Figure S14).

**Figure 3:**
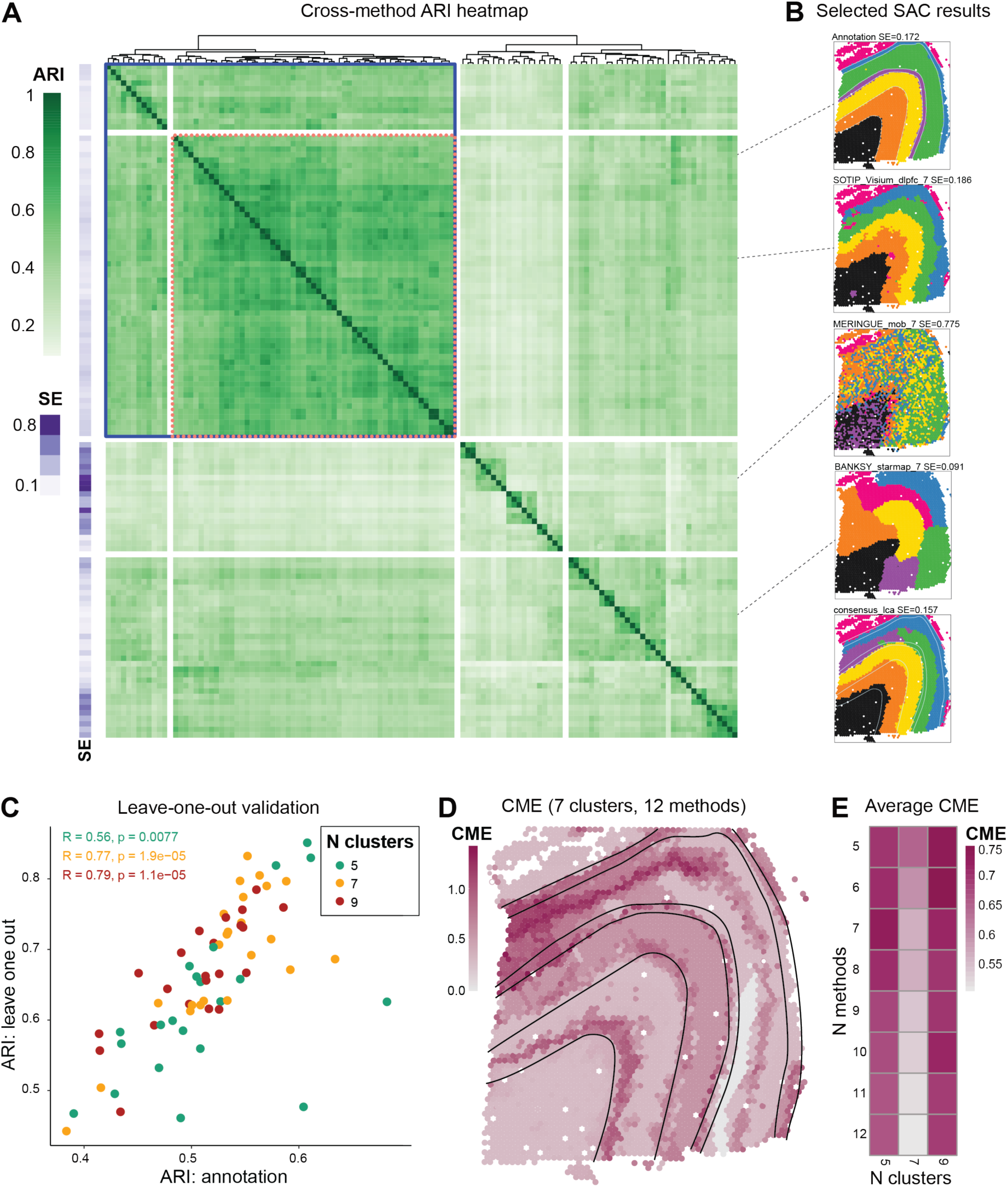
Consensus and entropy of spatially aware clustering. A. Method-vs-method ARIs for 119 base clusterings (including the manual annotation). The row annotations show the smoothness entropy (SE; lower SE means ‘smoother’ domains or, specifically, fewer total boundaries between domains). B. Selected base clusterings. Consensus_lca denotes a consensus clustering built from 12 methods at 7 clusters (D and E, see Methods). C. ARI of SAC methods calculated against a leave-one-out consensus versus the ARI against the original ground truth annotations. D. Cross-method entropy (CME; spot-wise similarity of cluster membership across methods) for 7 clusters and the 12 smoothest non-duplicated SAC methods. E. CME (averaged over all spots) for combinations of the number-of-smoothest-methods and number-of-clusters.

To determine whether combining outputs of methods with both high concordance (high pairwise ARI) and spatial continuity (low SE) could yield biologically relevant groupings, we selected a subset of SAC methods (see Methods for criteria) and formed a consensus clustering using latent class analysis (LCA) [50]. LCA was chosen for its scalability to large datasets, although K-modes and weight-based consensus approaches [51–53] were also implemented in the pipeline and can be applied based on dataset characteristics. We then applied a leave-one-out strategy among the pre-selected continuous and concordant methods (Figure 3B, blue box). For each method, we built consensus clusterings using all remaining methods, which then served as a pseudoGT to compare the method to. Comparing this leave-one-out consensus gives a readout of whether a method returns a significantly different clustering from other methods. While this is not meant to serve as a *bona fide* performance metric, clusterings that correspond well to pseudo-GTs also exhibit correspondence to the manual annotation (Figure 3C).

A by-product of consensus clustering is that we can spatially visualize where commonalities and uncertainties in base clusterings lie, for example, using a cross-method entropy (CME; lower values highlight concordance across methods; see Methods). To illustrate, we selected the 12 lowest-SE concordant-block base clusterings with seven clusters (Supplementary Figure S15) and generated a consensus clustering for slice 151673 of the LIBD DLPFC dataset (Figure 3B consensus lca). While the human brain structure of the LIBD DLPFC dataset is known to exhibit a layered structure, we included base clusterings without perfect layering (Supplementary Figure S15). Despite the inclusion of these imperfect clusterings, the consensus exhibited a smooth (SE = 0.17) and a mostly layered cluster organization (Figure 3B). To understand areas of disagreement, we visualized the CME across the set of base clusterings (Figure 3D). Compared to the GT, many heterogeneous regions align with the boundary regions marked by both consensus and anatomical labels. Additionally, while none of the methods were able to identify cortical layer 4 (L4) in this sample (Figure 3B Annotation, purple cluster), overlapping the L4 spots with the CME plot shows that the L4 layer aligns with a high entropy region, indicating the difficulty of correctly capturing this boundary based on transcriptional data at Visium resolution.

Furthermore, we can sweep across the number of clusters and the number of base clusterings (again prioritizing the most spatially continuous) to inform analysts about common structures found across granularities. Figure 3E highlights that 7 clusters induce the highest (average) correspondence between methods, and low CME remains stable even when selecting up to 12 base clusterings for the consensus. This approach can serve as a practical guide for an initial clustering of spatial data in the absence of a GT.

### Consensus with expert-in-the-loop

In practical research settings, SAC methods are rarely applied in a purely automated manner; instead, researchers frequently work in multidisciplinary teams and interact with multiple methods and parameter settings, iteratively evaluating their outputs to select those that best address specific biological questions. The consensus clustering and parameter sweeps integrated into our framework are designed to mimic this common research workflow, enabling biologists to systematically evaluate and compare a broad range of SAC method results within a unified environment. To demonstrate the utility of this approach in a real-world context, we collaborated with domain experts to evaluate SAC method performance on two modern SRT datasets: a MERFISH dataset of the [4] and a Visium HD dataset of human colorectal cancer [54] (see Datasets).

### Case study 1: Consensus SAC identifies anatomical boundaries in mouse thalamus

In the adult mouse brain, researchers have parcellated the thalamus region into approximately 40 anatomical labels, known as thalamic nuclei (Figure 4A), based on cytoarchitecture (cell shape, size, and density) and histochemical stains. However, this level of granularity is insufficient to capture the finer topographical patterns of neuronal connectivity and function that are known to exist within the thalamus [55].

**Figure 4:**
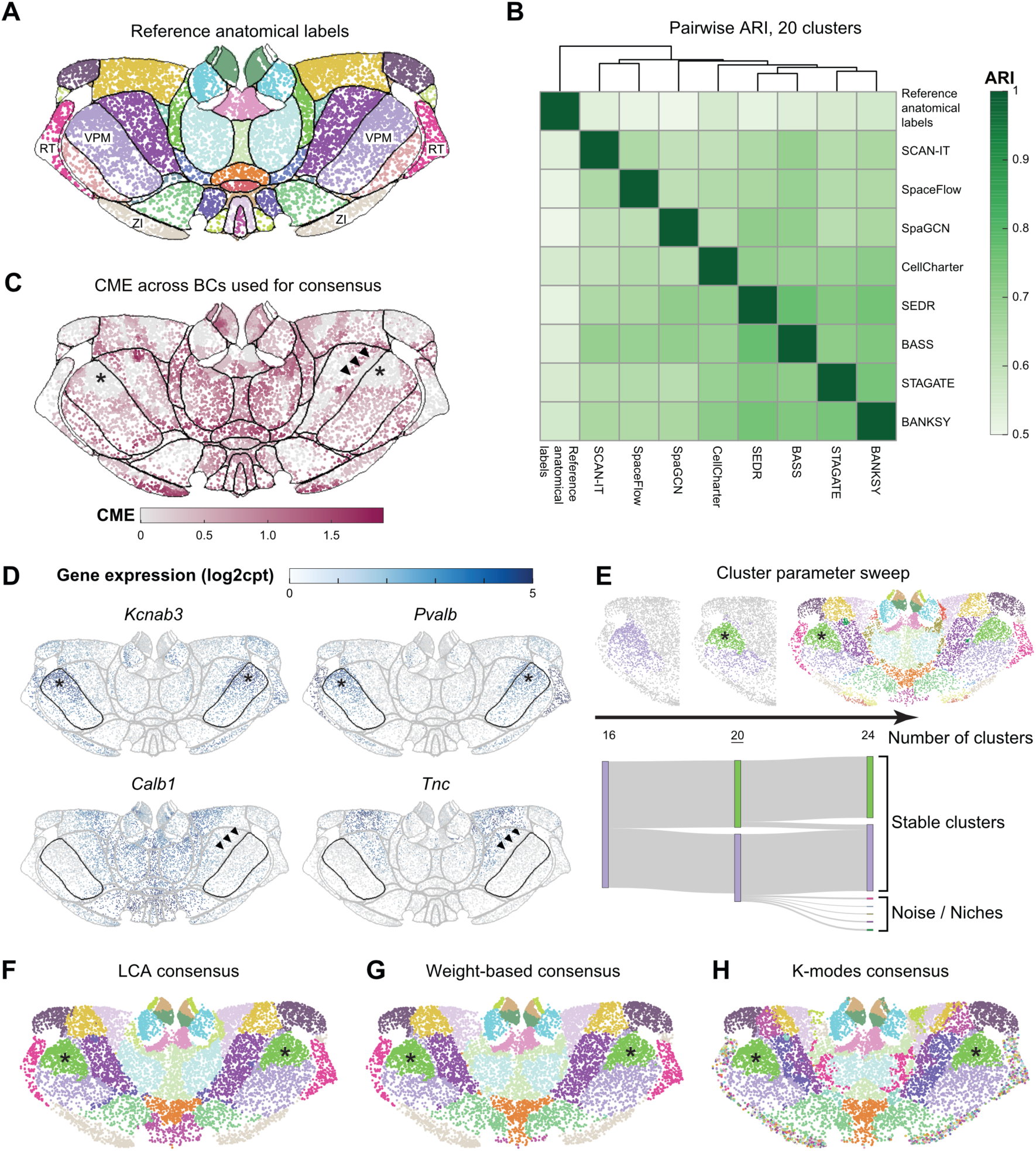
MERFISH dataset of the adult mouse brain thalamus region. A. Reference anatomical labels with borders indicated in black. VPM: ventral posteromedial nucleus; RT: reticular thalamic nucleus; ZI: zona incerta. B. Pairwise method-vs-method ARI comparison heatmap of the eight base SAC methods and the reference anatomical labels. C. Cross-method entropy (CME) of the eight base SAC results selected to build the consensus clusters. Reference anatomical label borders are indicated in black. Asterisks mark a low-entropy subdivision of VPM. Arrowheads mark a discrepancy between the reference anatomical borders and the SAC consensus in the right hemisphere. D. Gene expression of four genes (*Kcnab3*, *Pvalb*, *Calb1*, *Tnc*) that have spatially variable expression within or at the boundaries of the VPM subregion of the thalamus. The borders of all reference anatomical labels are outlined in grey; VPM is highlighted in black. E. Weight-based consensus of the datasets for 16, 20, and 24 clusters, with left VPM structure highlighted for the 16 and 20 cluster cases (top). 20 clusters (underlined) was used to generate the consensus clusterings in (F-H). Sankey diagram of the VPM region with the increase of cluster number (bottom). F. Consensus clustering computed with Latent Class Analysis (LCA) and clustering parameter set to 20 clusters. G. Consensus clustering computed using a weight-based framework and clustering parameter set to 20 clusters. H. Consensus clustering computed using K-modes and clustering parameter set to 20 clusters. Data from Yao et al. [4]; downloaded from the Allen Brain Cell Atlas.

SAC methods have the potential to identify novel, functionally relevant parcellations based on the information-rich and reproducible data of single-cell SRT. However, the results of eight SAC methods on the mouse thalamus MERFISH dataset showed a high level of discordance (Supplementary Figure S16). Without further refinement, this lack of agreement would only introduce confusion, rather than improve our understanding of molecular thalamic organization. Therefore, we leveraged our consensus clustering and SAC entropy analyses to interpret and apply the outputs of SAC methods on the same SRT dataset.

We compared the SAC results with each other and to the reference anatomical labels from the Allen Mouse Brain Common Coordinate Framework (CCFv3) [56] using pairwise ARIs (Figure 4B). Consistent with our findings in the Visium LIBD DLPFC dataset, we found that SAC methods tended to agree more with each other than with the anatomical reference labels (Figure 4A, B). This reiterates the limitations and risks of using anatomical annotations as GT for SAC evaluation.

Expert reviewers compared the resulting clusterings to both the reference anatomical labels and their domain knowledge of thalamic biology. The CME plot (Figure 4C) revealed regions of the thalamus that exhibited good agreement between SAC methods, as well as regions where the agreement was poor. High entropy (low cross-method agreement) frequently occurred near the boundaries between different anatomical labels. Two key insights emerged from this analysis. First, boundaries between thalamic nuclei often correspond to transitions in gene expression profile, and the entropy plot visually highlights how small differences in method choice or parameter setting can lead to variation in SAC calls at these transitions. Second, the entropy-indicated border helped us identify regions where the reference anatomical labels could be refined. While the MERFISH data was spatially registered to a whole brain reference coordinate space (CCFv3) to obtain the reference anatomical labels, this process is imperfect and results in misalignments, which are most notable at the borders between anatomical regions. The ventral posteromedial nucleus (VPM) is one example where the expression profile of key marker genes in the VPM (Figure 4D) aligns better with the entropy-indicated border than the boundaries from anatomical reference labels (Figure 4C-D; denoted by arrowheads).

Next, we generated consensus clusterings for the thalamus (Figure 4E-H). By sweeping the number of clusters from 16 to 24 (Figure 4E), we found cluster size and continuity to stabilize around 20 clusters, while higher cluster numbers resulted in more small and noisy clusters. We then generated consensus clusterings using three aggregation methods (LCA, weight-based, and K-modes), each set to 20 clusters (Figure 4F-H). Expert evaluation of these consensus clusterings revealed distinct behaviors among the aggregation approaches. Kmodes consensus subdivided many anatomical labels into multiple clusters and introduced a high number of salt-and-pepper (non-contiguous) clusters spread across two anatomical regions: the zona incerta (ZI) and the reticular thalamic nucleus (RT). This is consistent with additional transcriptional diversity and spatial structure observed within these areas [57, 58]. This stands in contrast to the weight-based and LCA consensus clusterings, which found one cluster each for ZI and RT.

Interestingly, we observed that some anatomical regions were consistently subdivided by all three consensus methods. For example, VPM was split into two clusters (Figure 4F-H; denoted by asterisks), reflecting the multi-gene expression gradients in the SRT data (Figure 4D; denoted by asterisks). The genes shown were chosen based on differential expression between the cells in the two VPM clusters (Methods) and may correspond to the spatial organization of neurons in VPM based on their axonal projections [59]. Given the high interest in identifying novel molecular parcellations of the thalamus, this robust consensus approach, combined with reassurance that we can find a stable number of clusters, is a promising target for subsequent experimental investigation with additional molecular and multimodal measurements.

### Case study 2: Detecting neoplastic progression stages in colorectal cancer

To evaluate our approach in a setting lacking established anatomical reference labels, we analyzed a previously unannotated SRT dataset of a pre-treatment human colon adenocarcinoma specimen sample profiled using Visium HD [54]. To incorporate domain expertise, we engaged both a histologist (A.E-H.) and a pathologist (M.Z.) specializing in colorectal cancer, who provided iterative feedback throughout the analysis process. This use case reveals another aspect of cluster annotation: a pathologist may not be equally interested in all regions, instead focusing their attention on pathophysiologically distinct areas.

Initially, the histologist annotated the tissue sections based on a high-resolution H&E image. The neoplasm (regarded as tumor tissue in this sample) boundary was then refined using *EPCAM* expression (Supplementary Figure S17), consistent with established FISH workflows (Figure 5A). These annotations served as a baseline for evaluating the performance of the SAC methods applied to the dataset. We observed a clear correspondence between expert annotation and the SAC results (Supplementary Figure S18), with a consistent difference being the split of the area annotated as neoplasm into two distinct clusters (Figure 5B). Only a subset of algorithms were able to process the dataset at full resolution (2 µm bins) due to memory and runtime limitations (Figure 1G), which limited our analysis to the 16 µm bin resolution. Before analyzing ARI scores, the two experts independently ranked each algorithm based on its ability to detect relevant tissue features, including: (i) distinguishing cancer from non-cancer tissue, (ii) identifying neoplastic differentiation, (iii) differentiating between all colon tissue compartments, (iv) identifying lymphovascular and perineural structures, (v) detecting small groups of neoplastic cells, (vi) identifying immune cells, and (vii) phenotyping of connective tissue (Figure 5C). Interestingly, this expert ranking differed substantially from the ARI scores calculated against the GT labels defined by the experts themselves (Figure 5D). DR-SC and STAGATE received the highest expert score while performing average in the ARI scoring, whereas the best-performing methods in the ARI metric, Seurat and CellCharter, did not reach the top rank as assigned by the experts. SEDR (“dlpfc” configuration) received the same rank as SCAN-IT, although the latter achieved a much higher ARI (Figure 5D). This lack of concordance between expert versus metric evaluation was also evident when evaluating the NMI, Smoothness Entropy, and CHAOS scores (Supplementary Figure S19). Our finding is consistent with previous results and highlights the limitations of global metrics, such as ARI, in evaluating performance for small, biologically relevant structures [60].

**Figure 5:**
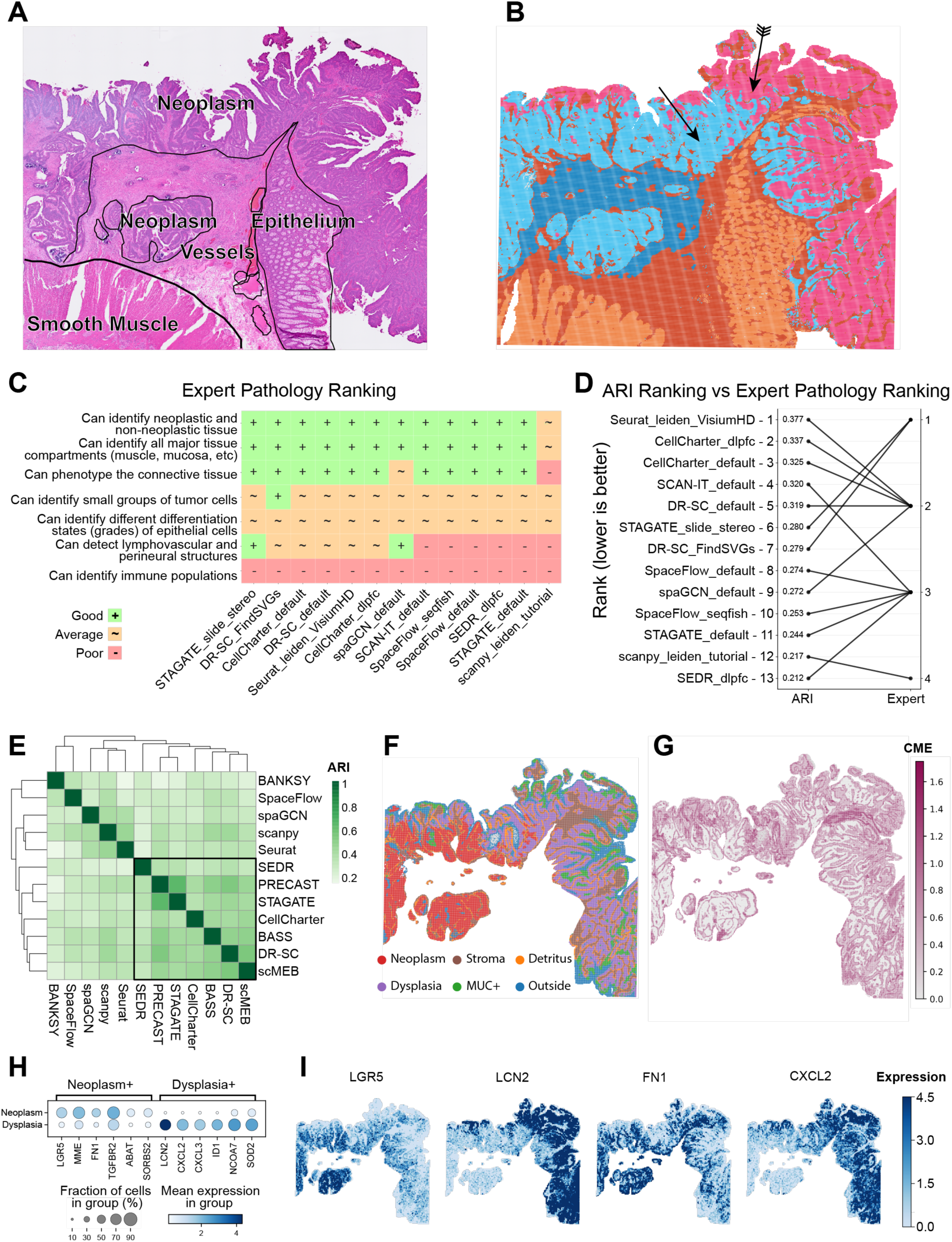
Colorectal cancer Visium HD Consensus with expert in the loop. A. Initial annotation of tissue regions based on the H&E image by an expert histologist. B. SEDR clustering (6 Clusters) identifies two distinct clusters in the dysplastic epithelium (feathered and smooth arrow). C. Expert ranking of SAC method performance for spatial feature identification relevant for pathological assessment (9 Clusters, full result can be found in Supplementary Figure S18). D. Comparison of the ARI calculated based on annotations (A) vs the expert ranking (C) for each SAC method. E. Method-vs-method ARI comparison heatmap for the neoplasm area. The boxed methods were identified as a concordant cluster based on Euclidean distance. F. Consensus clustering computed using K-modes for the 7 methods highlighted in panel E. G. Cross-method entropy map of the consensus clustering. H. Top differentially expressed genes between neoplasm and dysplasia. Color-coded by log1p normalized gene expression and size-coded by percentage of bins expressing the gene in the clusters. I. Spatial expression patterns of selected genes. Expression in log1p normalized counts.

Further investigation of nuclear morphology in the two distinct neoplasm clusters (Figure 5B, arrows) indicated a possible transition between tissue areas presenting neoplastic and dysplastic changes, characterized by increased nuclear-to-cytoplasmic ratio and nuclear crowding [61]. Annotation of these areas is usually carried out based on morphology [62], while more fine-grained molecular assessment of neoplastic cells has only recently emerged [63]. Thus, this sub-clustering of the neoplasm represents different scales of biologically relevant features that can be missed when rigidly evaluating the performance of SAC methods against manually annotated GTs.

To assess whether SAC methods could assist with this task, we subset the data to contain only bins in the neoplasm (based on the initial expert annotation) and re-applied available methods using default configurations. In the absence of a reliable way to delineate the neoplasm and dysplasia in the GT, we formed a consensus from methods showing a high concordance in terms of pairwise ARIs, reasoning that this could pinpoint pathological structures of interest (Supplementary Figure S20). Agreement between methods was generally high, as indicated by low entropy, with the exception of the boundary between tumor stroma and healthy epithelium (Figure 5G), which was the main area of disagreement between SAC methods. This might be caused by transitional zones, similar to what was observed in case study 1. The six distinct regions identified included (i) separate tumor and (ii) dysplastic zones, (iii) mucin signaling hotspots (glycoprotein-rich areas linked to specific colorectal cancer phenotypes [64]), (iv) tumor stroma, (v) cellular detritus split off from the tumor, and (vi) an external region (likely diffusion artifacts) outside the tissue (Figure 5F).

We found several well-studied differentially expressed genes between neoplastic and dysplastic areas (Figure 5H-I). The neoplasm showed high expression levels of *LGR5* (stem cell marker gene, investigated as a therapeutic target in CRC) [65], *FN1* (extracellular matrix remodeling) [66], and *TGFBR2* (conditional tumor suppressor/activator as part of TGF signaling) [67]. Conversely, the suspected dysplasia was characterized by the expression of *CXCL2/3* (inflammatory markers) [68], *LCN2* (a secretory protein potentially acting as a tumor suppressor in colorectal cancer) [69], and *SOD2* [70] and *NCOA7* [71] (stress response genes). Collectively, the gene expression patterns support the transition from a premalignant, inflammatory, and stressed dysplasia to a proliferative neoplasm with pro-invasive traits.

Our findings illustrate the value of an expert-in-the-loop approach where data-driven clusters combined with domain knowledge guide method selection and analysis direction, leading to biologically meaningful and novel insights. Notably, the experts did not initially set out to investigate the dysplasia to neoplasia transition – an observation that underscores the iterative and exploratory nature of investigating tissue architecture. Their understanding evolved as they interacted with the results, highlighting the importance of flexibility in clustering approaches. Taken together, this experience suggests that defining a single GT for spatial regions is often impractical, especially in complex tissues where biological heterogeneity and functional gradients are expected.

These case studies challenge the traditional paradigm of benchmarking SAC methods solely against fixed reference labels and emphasize the necessity of context-specific evaluation. They demonstrate that SAC methods can capture a spectrum of biologically relevant patterns, revealing insights that may be overlooked by rigid, single-label GTs. By embracing this multiplicity of interpretations, we can better leverage the full potential of SRT to uncover novel tissue architectures and disease mechanisms.

### Practical guidelines for an expert-in-the-loop workflow

Bioinformatics analyses are inherently iterative and question-driven, and spatially aware clustering analyses are no exception. Their strong dependence on biological context is what challenges the idea of a one-size-fits-all benchmarking framework and motivates the development of the expert-in-the-loop workflow. While our workflow has demonstrated its utility across several case studies, its broader application inevitably depends on case-specific conditions and expert interpretation. To summarize this complexity, we compiled a concise guideline outlining key considerations and checkpoints for the workflow, including approaches for expert involvement, parameter selection, and ensuring reproducibility (Supplementary File 1). Rather than prescribing a standardized best practice, this guideline serves as a reference manual to promote thoughtful and transparent use of the workflow across diverse analysis settings.

Whether studies are hypothesis-driven or exploratory, expert knowledge offers critical guidance that helps prevent misinterpretations and analytical pitfalls. The scope and nature of involvement of tissue experts inevitably vary across projects. In the case studies, we demonstrate how experts can guide clustering granularity (Figure 4E), assess base clustering performance (Figure 5C), and interpret consensus-level biological features (Figure 4F-H). Beyond these specific instances, the guideline outlines additional scenarios for expert participation and the reasoning behind them. We also illustrate how they translate into actionable steps by mapping analysis decisions to guideline checkpoints, exemplified by a detailed breakdown of analysis logic for Case study 1 (Supplementary Figure S21), which offers practical insights into the workflow’s implementation.

Base clustering (BC) and the selection of consensus algorithms are pivotal steps in the workflow. Strategies for BC selection can generally be classified into three categories: concordance-based, ranking-based, and manual selection. Manual selection relies on expert feedback or contextual biological criteria, while the other two approaches can be flexibly tailored to the research question at hand. In our case studies, base clusterings were chosen based on concordance blocks identified within the cross-method ARI heatmap (Figure 3B), as well as rankings of application-specific metrics that evaluate the spatial smoothness of clusters, such as SE (Supplementary Figure S22). Given the diversity of metrics included in the workflow, we encourage users to adapt these selection strategies to whichever metrics most effectively capture the biological phenomena of interest. Since consensus clustering aims to synthesize outputs from methods based on different modeling frameworks, a valuable sanity check during BC selection is to ensure a mix of clustering methods.

The choice of consensus algorithm should be tailored to the specific research question, taking into account tissue type, spatial technology, and the underlying biological hypothesis. In this study, our recommendations are primarily guided by scalability considerations (Supplementary Figure S23). We assessed runtime and memory usage for the three implemented consensus algorithms (K-modes, LCA, and weighted consensus) across a range of dataset sizes (i.e. number of cells or bins) and the number of input base clusterings. Overall, resource requirements increase with dataset size for all three methods. K-modes scaled best to large datasets, followed by LCA and weighted consensus. When varying the number of base clusterings, no consistent trend in runtime was observed; however, LCA showed a tendency toward higher memory consumption with increasing numbers of base clusterings. Across both variables, K-modes was the most resource-efficient, Weighted consensus was the most resource-intensive, and LCA showed intermediate performance.

A critical, yet often underexplored, aspect of spatial clustering is the investigation of the granularity of tissue features, analogous to the granularity considerations commonly addressed in cell typing for singlecell transcriptomics. We further examined the clustering granularity in the Xenium mouse brain dataset (Supplementary Figure S24). Although a broad range of cluster numbers corresponded well with established anatomical segmentations, clusterings with very low numbers were insufficient to resolve larger, coarse anatomical structures and very high cluster numbers tended to capture cellular heterogeneity rather than spatially contiguous anatomical domains. This finding highlights the importance of selecting an appropriate clustering resolution that balances anatomical interpretability with cellular detail when analyzing spatial transcriptomic data.

Although only two case studies were presented, the selected datasets represent two major classes of spatial transcriptomic technologies: imaging-based (MERFISH) and sequencing-based (Visium HD). They also cover tissue types that particularly benefit from spatial context, including brain and solid tumors. We expect that the practical guideline, comprehensive documentation, and aforementioned considerations will encourage the community to further evaluate and adapt the workflow across a wide range of settings. As a first step to facilitate those adaptations, we have included unsupervised (i.e., no expert involvement) consensus results from a selection of representative datasets covering different recent technologies (Supplementary Figure S25 to S29, Data Availability) as an initial resource to spur further investigation.

## Discussion

Existing benchmarks of SAC methods have generally focused on older SRT technologies and brain tissue datasets, often depending on questionable GT annotations or specific annotation scales, limiting the generalizability and real-world applicability of their results. To address these limitations, we developed SACCELERATOR, a consensus clustering framework designed to better reflect realistic research workflows, where researchers iteratively analyze data and work in multidisciplinary teams to interpret and refine findings. By integrating consensus of base clusterings with spatial metrics that highlight areas of high entropy (method disagreement), our approach enables targeted feedback for tissue experts. When applied to brain and cancer datasets with expert-in-the-loop evaluation, SACCELERATOR revealed biologically meaningful patterns that were previously overlooked, demonstrating that traditional evaluation metrics do not always reflect the subjective quality of results.

Our meta-analysis of reported performances across studies revealed considerable inconsistencies, highlighting the unreliability of current benchmarking practices. Method performances varied widely between studies, even when using the same datasets and algorithms. This variability can be attributed to several factors, including sensitivity to parameter choices and preprocessing steps, unreliable ground truths, and technology-specific performance. In particular, the impact of parameterization and preprocessing further complicates benchmarking efforts. To better reflect real-world usability, we evaluated methods using default settings and developer-recommended technology-specific configurations rather than greedily sweeping through multiple configurations.

To date, performance evaluation of SAC methods has largely relied on metrics that assess alignment with GT labels. Our results illustrate the limitations of solely relying on such metrics, given the inherent challenges in defining GT for complex tissues. Instead, we showed that a consensus clustering approach can be used to identify reproducible clusters, i.e., those consistently identified across methods and configurations, as well as pinpoint interesting regions of method disagreement. In our case studies, high entropy between methods highlighted biologically meaningful gradients, such as the transition from dysplasia to neoplasia in colorectal cancer. Such nuanced patterns cannot be captured by a single, discrete annotation. The consensus approach also showed robust subdivisions within anatomical regions, such as the VPM nucleus in the thalamus, highlighting its potential to uncover molecular markers for functionally relevant anatomical regions, such as the organization of thalamocortical cells in VPM [59]. Using our framework, we investigated a wide range of granularity levels in the Xenium mouse brain dataset. Our results indicate that for a range of clusterings, there is good correspondence with the expected anatomical divisions, while low numbers were insufficient to resolve larger structures and high numbers capturing cellular heterogeneity instead of anatomical domains.

Consensus among computational methods has been used in biomedical research as it conceptually overcomes limitations with the lack of ground truth knowledge [72]. In the context of SAC, the concept of consensus clustering is not new. For example, STCC offers a range of consensus strategies for SAC base clusters [73], but in our tests, STCC did not scale beyond Visium-sized datasets. Similarly, EnSDD, which also implements ensemble strategies based on binary similarity matrices [53], is unlikely to scale to datasets from newer technologies (the methods that overlap with our study did not scale to 2-µm-resolution Visium HD datasets). Conceptually, these frameworks utilize the consensus approach to better align with the GT. By contrast, SACCELERATOR employs consensus clustering as a guide for users and, explicitly, not as a means to maximize GT alignment. Beyond their biological relevance, consensus clusterings provide valuable feedback for method developers. If their method’s output diverges from the consensus, this can pinpoint unique strengths or limitations. One further aspect to note is that our consensus clustering is not using spatial information; therefore, methods from geography that impose spatial constraints may be valuable for future development [74].

As SRT technologies continue to improve in resolution and throughput, scalability becomes an increasingly critical concern. We found that many current SAC methods struggle with large datasets, particularly those from high-resolution platforms like Visium HD or imaging-based technologies capable of profiling large tissue areas. Only a small subset of methods could process the largest datasets within reasonable time and memory constraints, underscoring the urgent need for more computationally efficient SAC methods. In addition to the scalability of the individual methods with the number of cells and genes, the scalability of our proposed consensus-based clustering is also affected by the number of included base clusterings. Our results indicated K-modes as the most resource-efficient approach followed by LCA and finally the weighted-consensus. In addition to the number of base clusterings included, users should also consider the diversity of the selected methods in terms of their underlying algorithms. Since we envision the use of consensus clustering within an exploratory analysis without clear prior knowledge on the expected tissue clusters, aiming for a sufficiently diverse set of base clustering is desirable.

In our work, we rely on several metrics to compare clustering results between different methods. And while we rely on similar metrics to those reported in other studies (e.g. ARI), these are not intended as a critical or exclusive validation metric. Their use is explicitly described as a familiar reference for readers and to compare clustering assignments between computational approaches, not as the primary measure of performance or biological relevance. Our analysis of metric-metric correlation across various datasets revealed that intermetric relationships are generally similar, with general-purpose metrics (ARI, NMI) clustering separately from application-specific spatial metrics (CHAOS, PAS), and internal validation metrics (Davies—Bouldin index, Calinski–Harabasz index) also forming distinct clusters based on their focus on cluster compactness and separation. Overall, ARI and SE were found to be generally representative of the other metrics in most scenarios.

Although our study focuses on the use case of spatially aware clustering, the SACELLERATOR framework is generic and provides a flexible and robust foundation for advancing meaningful evaluation of computational tools in other areas of genomics research. By bridging the gap between computational analysis and biological interpretation, and by reflecting the collaborative and iterative nature of real-world scientific inquiry, our approach supports more accurate, reproducible, and biologically meaningful discoveries.

## Methods

### Datasets

We evaluated SAC methods across 15 published datasets representing both newer and older SRT technologies. Datasets were categorized based on their spatial resolution: mini-bulk, single-cell, and sub-cellular (Table 1). Only datasets with available GT annotations were considered for inclusion. To ensure biological relevance and technical diversity, we prioritized non-brain tissues datasets with sub-cellular spatial resolution.

The spatially resolved gene expression data, GT annotations, and further metadata were downloaded from original sources and converted to a standardized TSV/JSON format using our Snakemake pipeline. Since the datasets used in this study originate from publicly available sources that have already undergone careful curation by their respective authors, we applied minimal filtering. Specifically, we removed cells/bins with no gene expression, removed genes that were not expressed in any cell/bins, and we excluded cells in which mitochondrial transcripts accounted for more than 30% of total transcripts. For datasets generated using mini-bulk technologies, we filtered out bins containing fewer than 10 cells when this information was available. Furthermore, if the metadata contained bin-wise information of whether the bins overlapped with the tissue region, we excluded those outside the tissue area. No additional preprocessing was applied. Quality-filtered raw data were directly input into each method, following the method implementation or vignette without modification. Table 1 provides an overview of the analyzed datasets.

### CosMx human liver dataset (cosmx_liver)

The CosMx human liver dataset was obtained from the NanoString website (https://nanostring.com/products/cosmx-spatial-molecular-imager/ffpe-dataset/human-liver-rna-ffpe-dataset). The dataset consists of 2 Formalin-Fixed Paraffin-Embedded (FFPE) samples from 2 patients, one being normal liver and the other from a hepatocellular carcinoma patient with grade G3 cancer. Data was generated on the CosMx platform using the Human Universal Cell Characterization Panel 1000 plex. The ground truth annotations were computationally identified using Mclust clustering on the frequency of each cell type among its 200 nearest neighbors.

### CosMx human non-small-cell lung cancer dataset (cosmx_lung)

The CosMx human non-small-cell lung cancer dataset was obtained from the NanoString website (https://staging.nanostring.com/products/cosmx-spatial-molecular-imager/ffpe-dataset/nsclc-ffpe-dataset). The dataset consists of 8 FFPE samples from 5 patients presenting with non-small-cell lung cancer grade G1-G3. Data was generated on a CosMx prototype instrument using a 960 gene panel [75]. The ground truth annotations were computationally identified using Mclust clustering on the frequency of each cell type among its 200 nearest neighbors.

### MERFISH mouse brain thalamus (abc_atlas_wmb_thalamus)

The MERFISH mouse brain thalamus dataset [4] was obtained from the Brain Knowledge Platform (https://portal.brain-map.org/atlases-and-data/bkp/abc-atlas; https://alleninstitute.github.io/abcatlasaccess/descriptions/MERFISH-C57BL6J-638850.html). The dataset consists of 59 fresh frozen (FF) serial full coronal sections at 200 µm intervals spanning one entire mouse brain. Data was generated on a Vizgen MERSCOPE instrument using a custom gene panel of 500 genes. The ground truth annotations were identified by aligning the MERFISH data to the CCFv3 coordinate space and labeling cells with the corresponding CCFv3 anatomical parcellation term [56]. Only the thalamus (TH; CCFv3 structure ID 549) and hypothalamic zona incerta (ZI; CCFv3 structure ID 797) were analyzed in this study. Spatially variable genes in the thalamus were identified by differential gene expression analysis on neighboring consensus clusters.

### MERFISH human developmental heart dataset (merfish_devheart)

The MERFISH human developmental heart dataset [76] was obtained from Dryad (https://datadryad.org/stash/dataset/doi:10.5061/dryad.w0vt4b8vp). The dataset consists of 4 FF samples from 2 donors at 13 and 15 post-conception weeks (PCW). Data was generated using MERFISH with a custom 238-gene panel. The ground truth annotations (referred to as cellular communities in the original study) were computationally identified using *k* -means clustering of relative cell-type composition within 150 µm of each cell.

### STARmap PLUS mouse brain dataset (STARmap_plus)

The STARmap PLUS mouse brain dataset [77] was obtained from Zenodo (https://zenodo.org/records/8327576). The dataset consists of 20 FF samples from 3 mice. Data was generated using STARmap PLUS using a custom 1,022 gene panel. The ground truth annotations were manually identified by aligning the data to the CCFv3.

### Xenium human breast cancer dataset (xenium-ffpe-bc-idc)

The Xenium breast cancer dataset was obtained from the 10x website (https://www.10xgenomics.com/datasets/xenium-ffpe-human-breast-with-custom-add-on-panel-1-standard). The dataset consists of 1 FFPE sample from a patient with infiltrating ductal carcinoma breast cancer. Data was generated on a Xenium Analyzer using the Xenium human breast gene expression panel v1 (280 genes) with 100 additional custom genes. The ground truth annotation was manually identified using the matched histopathology image, annotating for eight region types: ductal carcinoma in-situ, invasive tumor, normal ducts, immune cells, cysts, blood vessels, adipose tissue, and stroma [78].

### Xenium mouse brain dataset (xenium-mouse-brain-SergioSalas)

The Xenium mouse brain dataset was obtained from the 10x Genomics website (https://www.10xgenomics.com/datasets/fresh-frozen-mouse-brain-replicates-1-standard). The dataset consists of 1 FF sample of a full coronal section. Data was generated on a Xenium Analyzer using the v1 mouse brain gene expression panel (247 genes). The ground truth annotation was manually identified using the mouse coronal P56 sample from Allen Brain Atlas [56] to specify anatomical regions [79].

### Slide-seqV2 mouse brain olfactory bulb dataset (slideseq2_olfactory_bulb)

The Slide-seqV2 mouse brain olfactory bulb dataset [80] was obtained from the STOmicsDB website (https://db.cngb.org/stomics/datasets/STDS0000172). The dataset consists of 20 samples of a mouse olfactory bulb evenly spaced along the anterior-posterior axis. Data was generated using Slide-seqV2 and sequenced using paired-end reads on an Illumina NovaSeq 6000 instrument, targeting 200 million reads per sample. The ground truth annotations were manually identified based on the expression of marker genes.

### Stereo-seq mouse liver dataset (stereoseq liver)

The Stereo-seq mouse liver dataset [45] was obtained from the STOmicsDB website (https://db.cngb.org/stomics/datasets/STDS0000059). The dataset consists of 6 FF samples. Data was generated on Stereo-seq chips and sequenced using paired-end reads on a DIPSEQ T1 instrument. The ground truth annotations were computationally identified where zonation layers were annotated based on the differences between the scores of pericentral and periportal hepatocyte landmark genes.

### Stereo-seq mouse embryo dataset (stereoseq_mouse_embryo)

The Stereo-seq mouse embryo dataset [13] was obtained from the STOmicsDB website (https://db.cngb.org/stomics/datasets/STDS0000058). The dataset consists of 53 FF samples from mouse embryos spanning E9.5–E16.5 with one-day intervals. Data was generated on Stereo-seq chips and sequenced using paired-end reads on a MGI DNBSEQ-Tx sequencer. The ground truth annotations were computationally identified using Spatially Constrained Clustering (SCC), which is built on top of the Leiden clustering algorithm [13].

### Visium human brain LIBD DLPFC dataset 1 (libd_dlpfc)

The Visium human brain LIBD DLPFC dataset 1 [35] was obtained from the spatialLIBD Bioconductor package (https://research.libd.org/spatialLIBD). The dataset consists of 12 FF samples from 3 donors. The data was generated on Visium chips and sequenced using paired-end reads on an Illumina NovaSeq 6000 instrument. The ground truth annotations were manually identified based on cytoarchitecture and selected gene markers.

### osmFISH mouse brain somatosensory cortex dataset (osmfish_Ssp)

The osmFISH mouse brain somatosensory cortex dataset [81] was obtained from the Linnarsson Lab website (https://linnarssonlab.org/osmFISH). The dataset consists of a single FF sample from the mouse brain somatosensory cortex. Data was generated using osmFISH using a custom 33-gene panel. The ground truth annotation was computationally identified using an iterative graph-based algorithm.

### Visium human breast cancer (visium breast cancer SEDR)

The Visium human breast cancer dataset, originally from 10x Genomics (https://www.10xgenomics.com/resources/datasets/human-breast-cancer-block-a-section-1-1-standard-1-1-0), was obtained from GitHub (https://github.com/JinmiaoChenLab/SEDR analyses). The dataset consists of a single FF sample of invasive ductal carcinoma breast tissue. The data was generated on a Visium chip and sequenced using paired-end reads on an Illumina NovaSeq 6000 instrument. The ground truth annotation was manually identified based on the H&E image.

### Visium chicken heart (visium_chicken_heart)

The Visium chicken heart dataset [82] was obtained from GEO (https://www.ncbi.nlm.nih.gov/geo/query/acc.cgi?acc=GSE149457). The dataset consists of 11 FF samples from four hearts at different stages of ventricular development. The data was generated on a Visium chip and sequenced using paired-end reads on an Illumina NextSeq 500/550 instrument. The ground truth annotations were computationally identified using Louvain clustering as implemented in Seurat v3.

### Visium HD human colorectal cancer dataset (visium_hd_cancer_colon)

The Visium HD human colorectal cancer dataset [54] was obtained from the 10x Genomics website (https://www.10xgenomics.com/datasets/visium-hd-cytassist-gene-expression-libraries-of-human-crc). The dataset consists of 1 FFPE sample of colorectal cancer obtained from the sigmoid colon of a patient. Data was generated on a Visium HD chip and sequenced using paired-end reads on an Illumina NovaSeq 6000 instrument.

### Ground truth annotation of the Visium HD human colorectal cancer dataset

At the time of analysis, no ground truth annotation was available for the Visium HD human colorectal cancer dataset. After data download, annotation of the Visium HD colorectal cancer histology sections was performed on H&E-stained whole-slide images by a trained histologist. Regions of smooth muscle, vessels, normal epithelium, and neoplastic epithelium were manually annotated, followed by review by a pathologist. Smooth muscle was identified by its characteristic pink eosinophilic staining in organized bundles, with elongated, centrally located nuclei. Blood vessel regions were annotated based on the presence of luminal structures, filled with erythrocytes and lined by flat cells (endothelium). Normal epithelium was identified by recognizing well-organized crypt architecture, composed of columnar epithelial cells with basally oriented nuclei, low nuclear-to-cytoplasmic ratios, and regular cell polarity. Neoplastic epithelium was annotated by identifying regions with loss of normal crypt architecture, increased nuclear pleomorphism, hyperchromasia, irregular nuclear contours, aberrant nuclear-to-cytoplasmic ratios, and loss of polarity. Stroma or connective tissue was identified by its bright eosinophilic staining, low cellular density, presence of spindle-shaped fibroblast-like cells, and visible connective tissue fibers (e.g., collagen). Rare interspersed immune cells, such as neutrophils, lymphocytes and others, could occasionally be observed. Stroma was typically located adjacent to epithelial or neoplastic areas. Dense accumulation of lymphoid cells, including lymphocytes and histiocytes would represent lymphoid structures. However, such structures were not observed in the present CRC case. Dissociated cell material indicative of detritus or necrosis was identified by the presence of cellular debris, often accompanied by infiltration of neutrophilic granulocytes. Annotations were further refined by expression of the characteristic marker genes *EPCAM* (epithelial neoplasm), *VWF* (vessels), and *MYL9* (smooth muscle). The annotations were attached to individual Visium HD bins by annotation in SpatialData [83] with the help of the Napari interface [84].

### Spatially aware clustering methods

We implemented all 22 SAC methods according to their original publications and official documentation. To ensure fidelity to method designs, preprocessing steps (e.g., normalization, feature selection) were replicated exactly as described in method vignettes or GitHub repositories. For parameter configurations, we included a “default” configuration, where we used author-predefined parameter values, with undefined parameters set via prominent tutorials or GitHub READMEs. Additional configurations were limited to those available in the method documentation, where parameters were set as specified in case-specific vignettes. For added provenance tracking, the configuration files include a reference to their sources.

Graph-based SAC methods often use resolution parameters to control cluster granularity that ultimately dictates the number of resultant clusters. To enable fair comparison across metrics requiring fixed cluster counts, we performed binary searches for the resolution values yielding the number of clusters defined in the GT annotation. This approach ensured compatibility with spatially agnostic metrics like adjusted Rand index (ARI) while maintaining method-specific spatial constraints.

Additionally, we included the most prominent single-cell transcriptomics clustering algorithm, Leiden, as implemented in Seurat and Scanpy. See Supplementary Table S1 for a complete overview of the implemented methods.

### Metrics

We implemented 17 metrics in the framework, categorized based on their use of GT labels or spatial information. See Supplementary Table S3 for a complete overview of the implemented metrics. In addition to the conventional clustering metrics, we developed two entropy-based metrics. Both metrics require the calculation of Shannon entropy. In our implementation, if we have a vector of cells from different classes (i.e., clusters), the entropy can subsequently be calculated as:

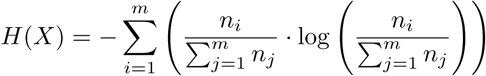

where *m* is the number of classes, *n_i_* is the occurrence of class *i* in vector *X*, and 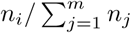 is the empirical probability of class i.

### Smoothness entropy (SE)

SE quantifies the spatial coherence of clustering results by measuring label consistency within local neighborhoods. The SE is calculated by identifying the *k* -nearest neighbors for each spot or cell in the dataset. Subsequently, the entropy is calculated spot-wise across the labels of the spot and its *k* neighbors. Spot-wise entropy is then averaged to provide the SE for the sample on a particular label. For SE shown in Figure 3, *k* is set to 6.

### Cross-method entropy (CME)

CME identifies regions of uncertainty by measuring label discordance across multiple clustering results. To generate the CME for a dataset, a set of labelings with the same number of clusters is aligned by finding the optimal matches across different labelings using the Hungarian method implemented in the R package clue (version 0.3 66) [86]. Once the labelings are aligned, spot-wise entropy is calculated across the labelings from different methods. The labeling alignment requires a reference labeling where other labelings are aligning against. To avoid stochasticity from arbitrary selection of reference labeling, the entropy is calculated once for each instance where a labeling is selected as the reference. The final spot-wise CME is calculated by averaging the entropies from different reference labeling cases.

### Computational framework

SACCELERATOR is orchestrated by a modular Snakemake [87] workflow, which coordinates all computational steps for standardized and reproducible analysis. The framework is organized into distinct modules for datasets, SAC methods, metrics, and consensus clustering. Each dataset module consisted of a script to download the dataset, GT annotations, images, and other metadata and store them in a standardized format. Likewise, each method and metric module consisted of a separate script, ensuring interoperability with the standardized data structure produced by other modules. Consensus clustering within SACCELERATOR is implemented as a three-step process: results aggregation, base clustering selection, and consensus generation. Each of these steps is documented with a minimally executable example, clear instructions to add new instances, and explanatory notes in its respective folder in the GitHub repository and the documentation (https://spatialhackathon.github.io/SACCELERATOR). To guarantee reproducibility and prevent dependency conflicts, each script is linked to a version-pinned conda environment file. A centralized configuration file allows the Snakemake workflow to specify parameters for each module, such as which datasets to download and the number of clusters to identify.

### Consensus clustering algorithms

Three consensus clustering approaches were implemented in the framework and applied in the paper: Kmodes, Latent Class Analysis (LCA), and weight-based consensus. While many consensus algorithms exist, these three were selected as representative examples to demonstrate the flexibility of the consensus workflow. K-modes was selected as the categorical analogue of *k* -means. In the context of consensus clustering, each base clustering contributes to the consensus clustering equally, and their labels serve merely as nominal categories without ordering or magnitude, which aligns with the assumption of the K-modes method. As an algorithmic approach to consensus clustering, K-modes aims to find clustering partitions that minimize the overall hamming distance to cluster-specific mode vector [88]. This approach returns interpretable results in a scalable manner, especially for big datasets (Supplementary Figure S23). In the SACCELERATOR framework, K-modes is implemented via R package diceR [89].

LCA was chosen as a representative model-based consensus clustering approach. LCA assumes that there exists an unobserved categorical latent class (the consensus clustering) that explains the observed base clusterings, and that individual base clustering results are conditionally independent given the latent class. Unlike the distance-based K-modes, LCA is a probabilistic approach that allows for flexible handling of base clustering disagreement by method-specific conditional probabilities [90]. In the SACCELERATOR framework, LCA is implemented via R package diceR [89].

Additionally, we also implemented a weighted-based consensus based on a Bregmannian divergence framework [52, 53] utilizing the weighted Jensen-Shannon Divergence (JSD) as the loss function [91]:

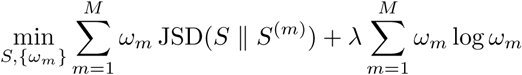

subject to

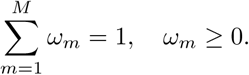

where *M* is the number of base clusterings, *S* is the binary similarity matrix for consensus results, *S*^(*m*)^ is the binary similarity matrix of individual base clusterings, and *ω_m_* are the respective weights of each base clustering. The second term is an entropy regularization of the weights to prevent the inclination of the binary similarity matrix to any single base clusterings. The resulting binary similarity matrix *S* was clustered using the Leiden algorithm to generate the final consensus clusters. Unlike LCA and K-modes, the weighted consensus also returns the weight terms assigned to each base clustering, providing an interpretable understanding of the relative contributions of different methods. However, the incorporation of large binary matrices limits the scalability of this method (Supplementary Figure S23).

Although the three approaches (K-modes, LCA, and weighted consensus) are already integrated into the workflow, we encourage users to explore additional consensus algorithms as needed, such as majority voting or linkage clustering ensemble [92]. The choice of method should be guided by both computational considerations (e.g., scalability, Supplementary Figure S23) and the specific biological questions of interest, ideally in consultation with domain experts (see Supplementary Figure S21 for an overview of the expert-guided workflow instruction).

The cluster scalability results were generated using the benchmark feature of Snakemake. The consensus part of the pipeline was applied to all existing clustering results of datasets with the following names: xenium-mouse-brain-SergioSalas, abc atlas wmb thalamus, libd dlpfc, visium hd cancer colon, cosmx lung, STARmap plus, visium breast cancer SEDR. Results of those consensus results can be found on Zenodo (Data and Code Availability).

### Base clustering selection

Base clustering selection is an essential process for generating consensus clusterings. As implemented in SACCELERATOR, this process is designed to be flexible and context-dependent, reflecting the expert-in-theloop workflow. There is no universally optimal criterion for selecting the “best” base clusterings for consensus clustering; rather, selection should be tailored to the specific technological platform and biological context under study. In the analysis-specific sections below, we describe in detail how base clusterings are selected based on different biological settings.

### Meta-benchmark analysis

We collated ARI values for the Visium LIBD DLPFC dataset from original method papers and benchmarking studies, sourcing data from the main text and supplementary material of published articles, GitHub repositories, and Zenodo repositories. In total, we gathered results for 22 SAC methods that we implemented and three benchmarking studies [5, 16, 17]. Among these, 17 method papers and one benchmarking study [16] reported ARI values for the DLPFC dataset. BayesSpace, one of the earliest SAC methods, was the most frequently benchmarked, with ARI reported in 16 studies. We excluded studies that only reported summary statistics (e.g., mean or median ARI) or results from a single tissue slice. For comparisons of self-reported ARI with those from other studies, we excluded methods that had not been independently applied by at least one subsequent study.

### Computational resource management

All SACCELERATOR computational jobs were orchestrated using Snakemake (v8.14.0) for workflow management and SLURM (v21.08.5) for workload management on the de.NBI cloud infrastructure. Each method was executed using a dedicated allocation of 10 CPU threads and 500 GB of RAM per job. For each sample, the maximum allowed runtime was set to 48 hours. If a job exceeded this limit, it was automatically terminated and recorded as a failure for that sample. Resource utilization, including wall-clock runtime and peak memory usage, was monitored and recorded using Snakemake’s built-in benchmarking functionality. Benchmarking data were parsed and summarized using custom Python scripts.

### Configuration benchmarking

For the Visium DLPFC dataset, we calculated the ARI values for all methods using the same number of clusters and method-specific configurations. Methods without additional configurations were excluded. This resulted in ARI values for 18 methods, each with 2 to 6 configurations. For details on the configuration settings, refer to the section on Spatially Aware Clustering Methods.

### GT granularity analysis

To generate the PCA plots for the LIBD DLPFC dataset shown in Figure 2A and B, we performed quality control on slice 151673 by filtering out spots with less than 600 total mRNA counts or more than 28 percent mitochondrial transcript content. Spots with fewer than 10 cells detected were also excluded. The 10% most highly variable genes were detected for the sample via getTopHVGs() function in R package scran (version 1.34.0) [93]. PCA was run on the highly variable genes via function runPCA() in the R package scater (version 1.34.1) [94]. To identify border spots, we constructed a *k* -nearest neighbor graph based on the spatial coordinates of the spots using function kNN() from R package dbscan (version 1.2.0) [95] with *k* = 4. We filtered out neighbors with a distance larger than 1.5. A spot was deemed a border spot if any of its neighbors, including itself, did not belong to the same cluster as the GT labels.

For the Xenium mouse brain dataset and the Visium breast cancer dataset presented in Figure 2, SAC results were generated using the default configurations of five methods (Supplementary Table S2). To examine the relationship between ARI and cluster granularity, all methods were configured to identify various numbers of clusters for the datasets (Xenium: 5, 7, 9, and 11; Visium: 3–7). The ARI against the GT annotation was then calculated using the implemented workflow.

The granularity scanning for the Xenium Mouse Brain Dataset (Supplementary Figure S24) is conducted using the default configuration of CellCharter in the workflow while setting cluster numbers from 5 to 49. Results of the cluster sweeping have been uploaded to Zenodo (Data and code availability), and additional visualization of the sweeping results can be found in Supplementary File 2.

For the Stereo-seq mouse liver dataset, we subsetted the datasets to the 925 unique zonation-related genes reported in three studies [47–49]. Five methods were then run on both the full and subsetted liver dataset with a memory and runtime cap of 500 GB and 48 hours. All methods except STAGATE managed to generate clustering results. The four remaining methods were configured to generate between 3 and 9 clusters. In addition to the default 9-layer GT, we aggregated the layers into 3 regions according to the original study. Specifically, Layer 1 and 2 were aggregated as the pericentral cluster, Layer 4 to 6 were merged into a midzonal cluster, and the rest into the periportal cluster. ARI values were calculated for the SAC results against both the original nine-layer and aggregated three-region GTs.

### LIBD DLPFC Visium consensus analysis

All base clusterings generated from the workflow for slice 151673 (with 5, 7, or 9 clusters; Supplementary Table S2) were aggregated to compute pairwise ARIs (Figure 3A). To identify a concordant block of SAC methods, we applied cutoffs to the hierarchical clustering tree (euclidean distance, complete linkage) of the pairwise ARIs, which revealed two groupings, one of which represented a highly concordant set of 14 distinct clustering methods (Figure 3A; blue box). From this set of concordant methods, base clusterings were selected as input to LCA consensus clustering [50], separately for each number of clusters (5, 7, 9), according to their SE, with priority given to smoother base clusterings. For example, Figure 3D (“7 clusters, 12 methods”) represents a consensus clustering of the 12 smoothest (i.e., lowest SE) 7-cluster base clusterings. In some cases, we have multiple clusterings from the same algorithm (different parameter configurations) with the same number of clusters; to build the consensus (as described above), however, we enforce algorithm diversity by only keeping one base clustering (the smoothest) per algorithm. For the leave-one-out validation, a consensus clustering (LCA) was generated for each method, separately for each number of clusters. Each consensus comprised the 6 smoothest base clusterings of the starting set of methods (blue box of Figure 3A), excluding the method itself.

### Thalamus consensus analysis

Thalamus sample C57BL6J-638850 6800 was selected to showcase the consensus workflow. We first generated SAC results of the sample from existing method configurations in the workflow (Supplementary Table S2). Then, we filtered out any base clusterings with a prominent class imbalance, where one cluster comprised more than 90% of the cells. Next, the eight methods with the lowest SE were selected as base clusterings for the consensus clustering for all three consensus algorithms. A consensus was calculated separately for base clusterings with 16, 20, and 24 clusters. The cluster numbers were selected based on expert suggestions.

### Visium HD consensus analysis

The dataset was prepared as described in the dataset section above. Multiple configurations were employed for the tested SAC methods (Supplementary Table S2). For the full-section analysis, the pipeline was configured to identify 5, 6, 7, 9, 11, and 14 clusters. For the consensus calling, the sample was subset to only the neoplasm region, and the pipeline was configured to identify six clusters. The base clustering selection was done similarly to the LIBD DLPFC example (see above), via selecting a core set of “concordant” methods according to pairwise ARIs and setting cutoffs on the hierarchical clustering tree (black box in Figure 5E). As with the LIBD DLPFC consensus approach, the consensus clustering was generated using the LCA algorithm [50]. Marker genes for the neoplastic region and dysplasia were calculated with the tie-corrected Wilcoxon rank-sum test implemented in Scanpy [96].

## Data and code availability

The SACCELERATOR workflow and all relevant code to run the analysis are available as open-source software on GitHub (https://github.com/SpatialHackathon/SACCELERATOR). All code for reproducing results presented in this study, including figures and collected ARIs for the meta benchmark are available via https://github.com/SpatialHackathon/SpaceHack2023 study. Links to raw data are provided in the Datasets section. All filtered data, intermediate results, and final results are deposited on Zenodo (https://zenodo.org/records/15487519).

## Author contributions

NI, AM, MDR conceptualized the project. NMB implemented the concept of the benchmarking framework. NMB, FH, KB, JS, MAT, PK performed data curation. JS, KB, PC, NMB, SHD, MDR, PK performed formal analysis. NI and MDR acquired finances. NMB, JS, MDR, SK, SG, ME, PK, SHD, PC developed and designed the methodology. NMB, FH, JS, KB, PC, SHD, PK implemented and tested the softwares. RE, ST, RG provided critical resources. ST established the SpaceHack cloud infrastructures. NI, MDR, AM, CK, RE, RG, BL performed supervision. MAT, AEH, MZ performed validation. JS, KB, PK, MDR, PC, MAT, NMB performed visualization. NI, AM, SHD, JS, MDR, KB, PC, MAT, PK wrote the original draft of the manuscript, and NI, AM, FH, SHD, JS, MDR, MAT, MR, NMB reviewed and edited the manuscript.

## Acknowledgments

We thank the organizers and participants of the de.NBI BioHackathon SpaceHack 2.0 project in Bielefeld, Germany in December 2023 (Supplementary Table S4). The SpaceHack 2.0 project was funded by the German Federal Ministry of Education and Research in the frame of de.NBI & ELIXIR-DE (W-de.NBI-001). We also thank Martin Braun and Valentin Schneider-Lunitz for their support with the cloud.

