## Supplementary Figures for "Beyond benchmarking: an expert-guided consensus approach to spatially aware clustering"

### Supplementary Materials

#### Supplementary Figures

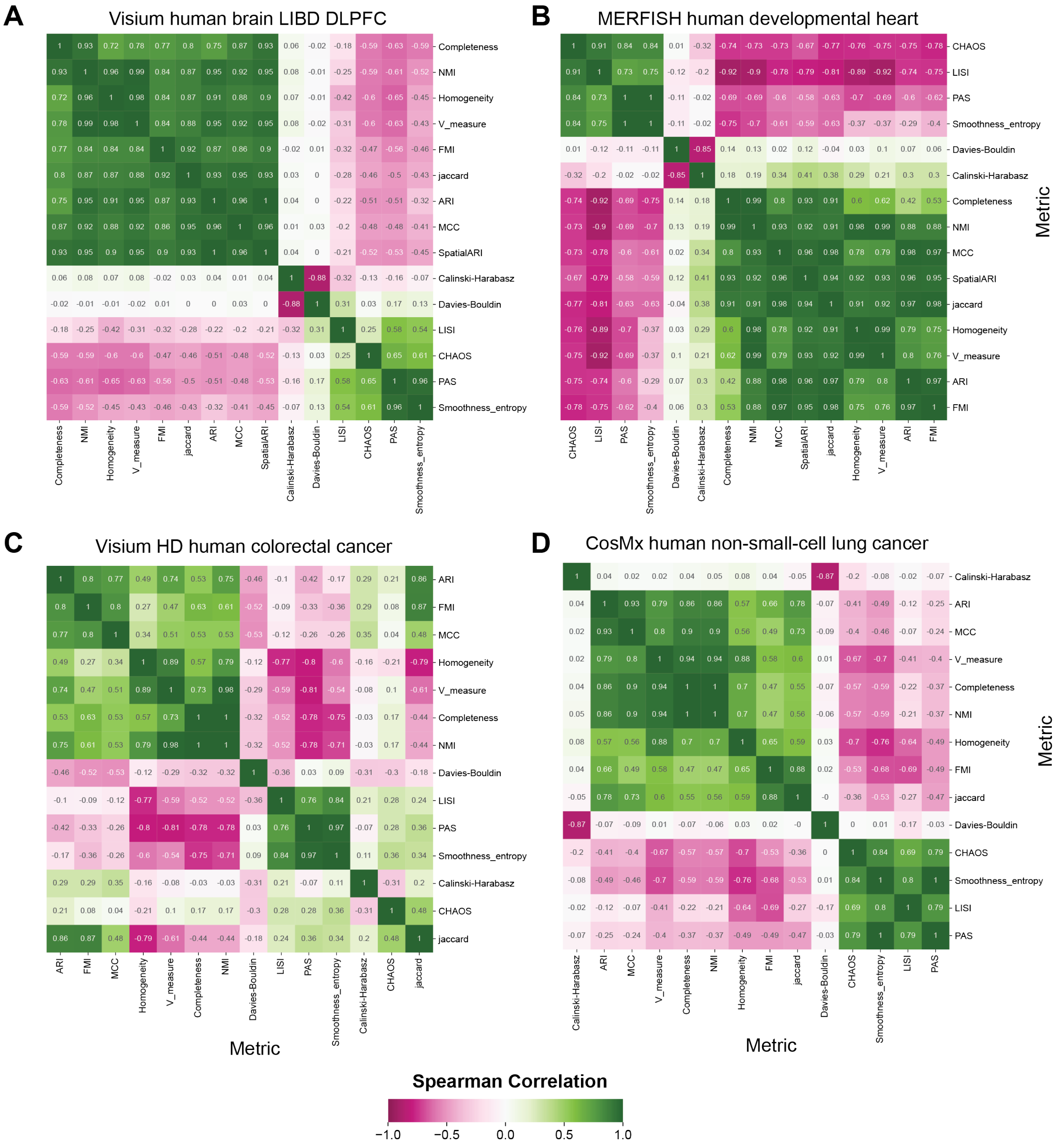


**Figure S1**: Spearman correlation matrix of different metrics across all methods and samples from **A**. LIBD DLPFC, **B**. MERFISH human developmental heart, **C**. Visium HD human colorectal cancer, **D**. CosMx lung.

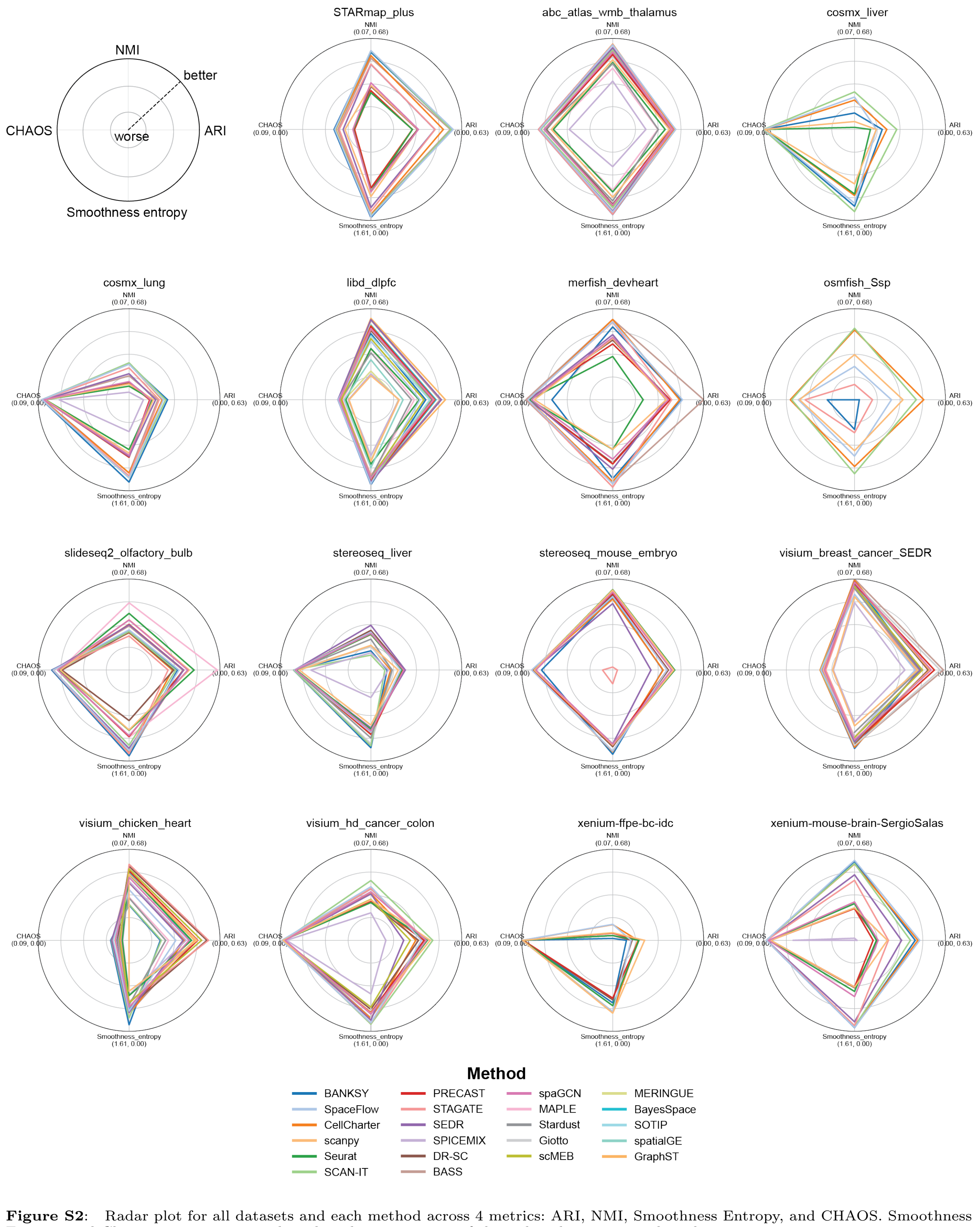


**Figure S2**: Radar plot for all datasets and each method across 4 metrics: ARI, NMI, Smoothness Entropy, and CHAOS. Smoothness Entropy and Chaos scores are inverted so that the outer range of the radar plot corresponds to better scores.

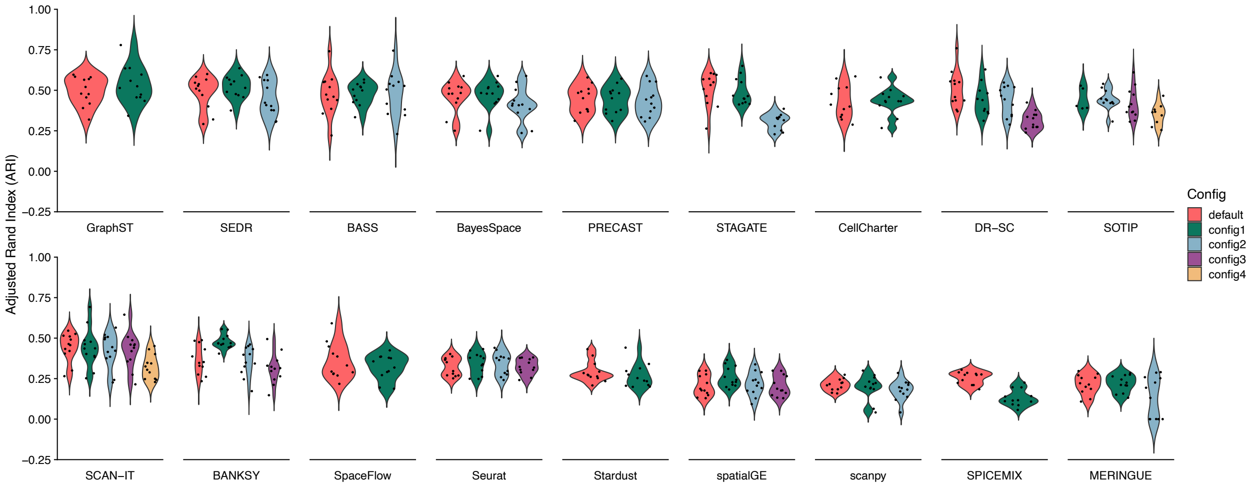


**Figure S3**: ARI for methods with multiple configurations applied to the LIBD DLPFC dataset. Methods are ordered by mean ARI.

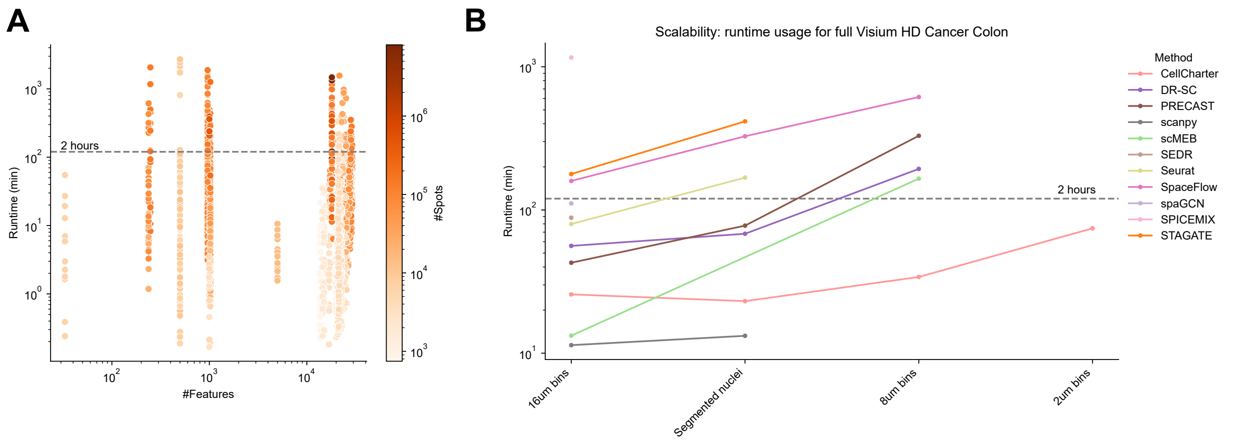


**Figure S4**: **A**. CPU runtime vs the number of features, colored by the number of spots/cells in the dataset. **B**. CPU runtime of methods on the full Visium HD colorectal cancer dataset across various bin sizes.

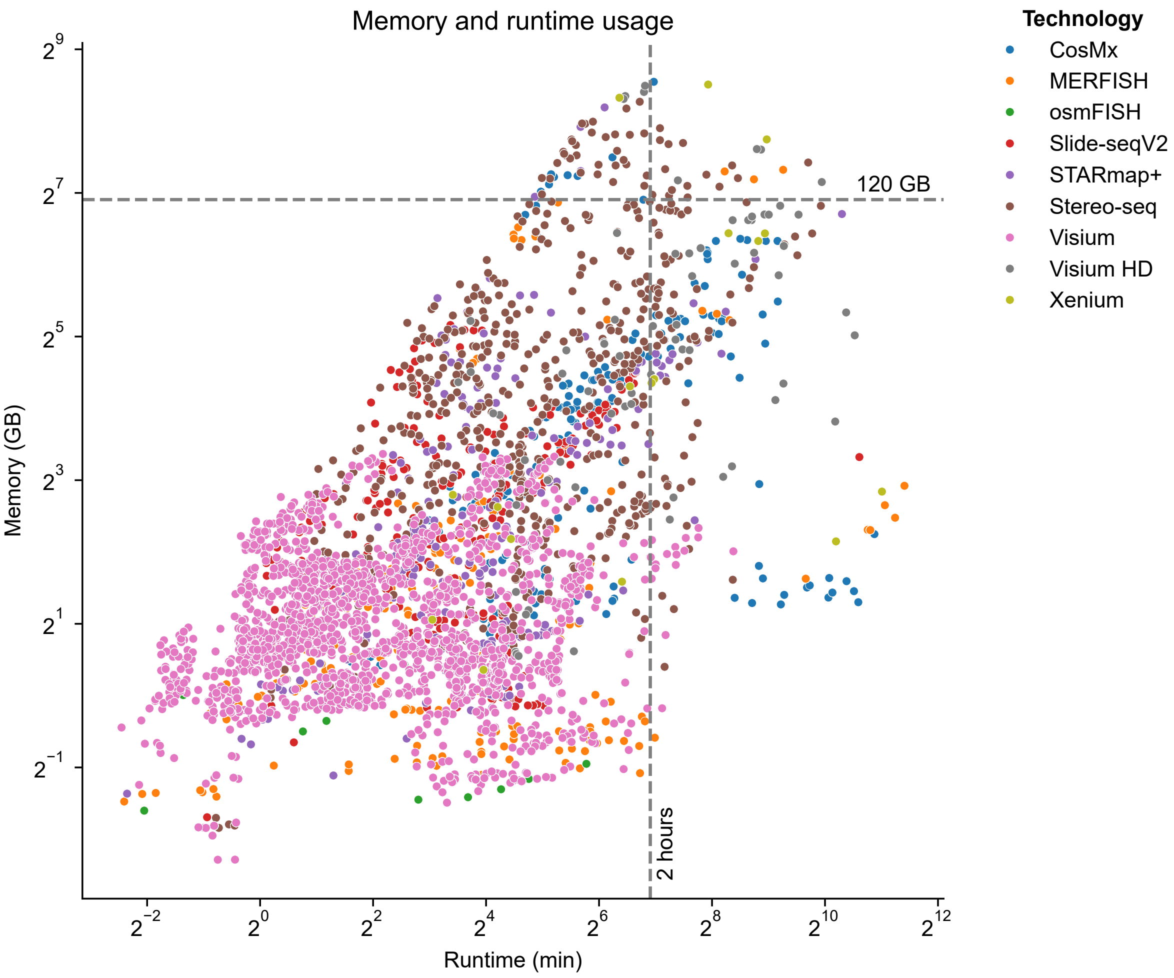


**Figure S5**: Memory usage vs CPU runtime, colored by the technology of the dataset.

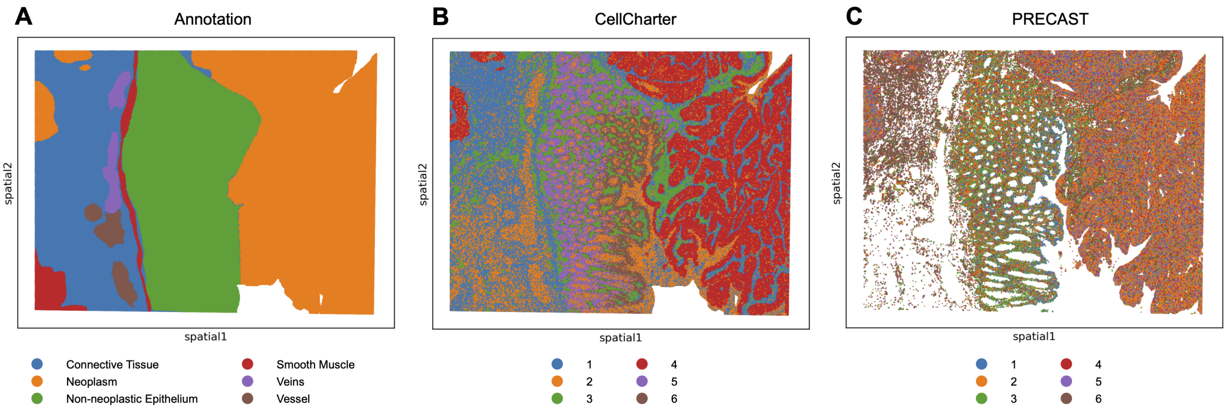


**Figure S6**: Spatial plots of the cropped Visium HD colon cancer dataset with 2 m bins, colored by **A**. ground truth, **B**. CellCharter clusters, and **C**. PRECAST clusters.

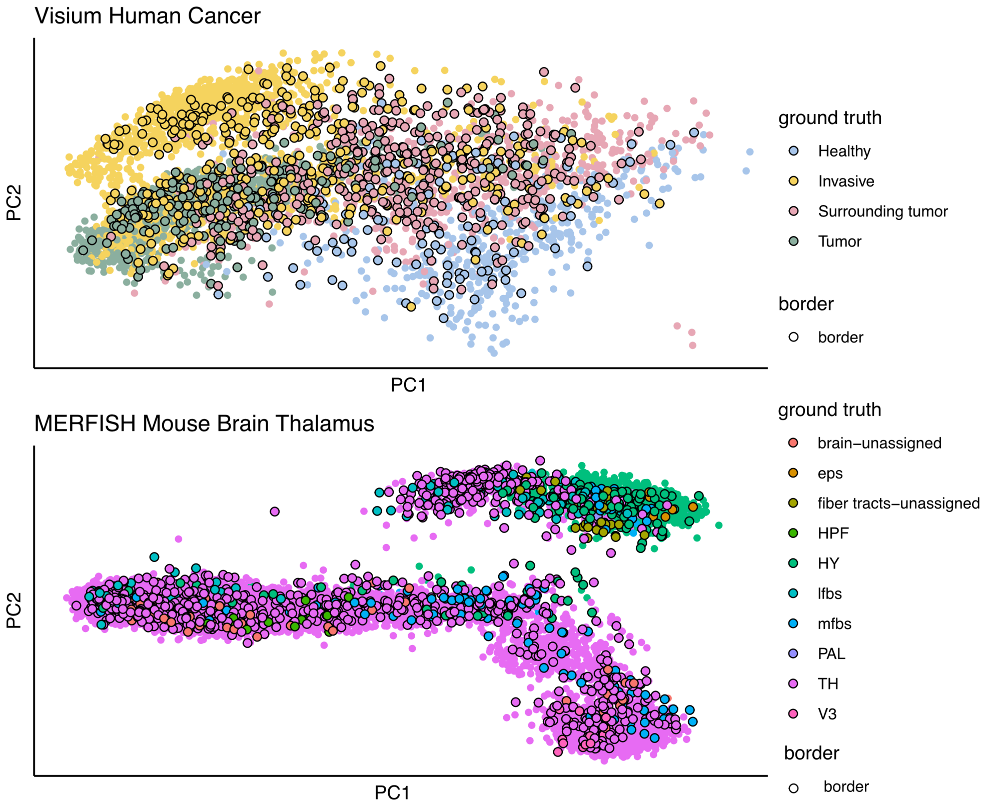


**Figure S7**: Visium human cancer, and MERFISH mouse brain thalamus datasets, projected on the 1st and 2nd principal component (PC), colored by GT labels. Spots that are spatially located at the border between GT clusters are marked with black outlines.

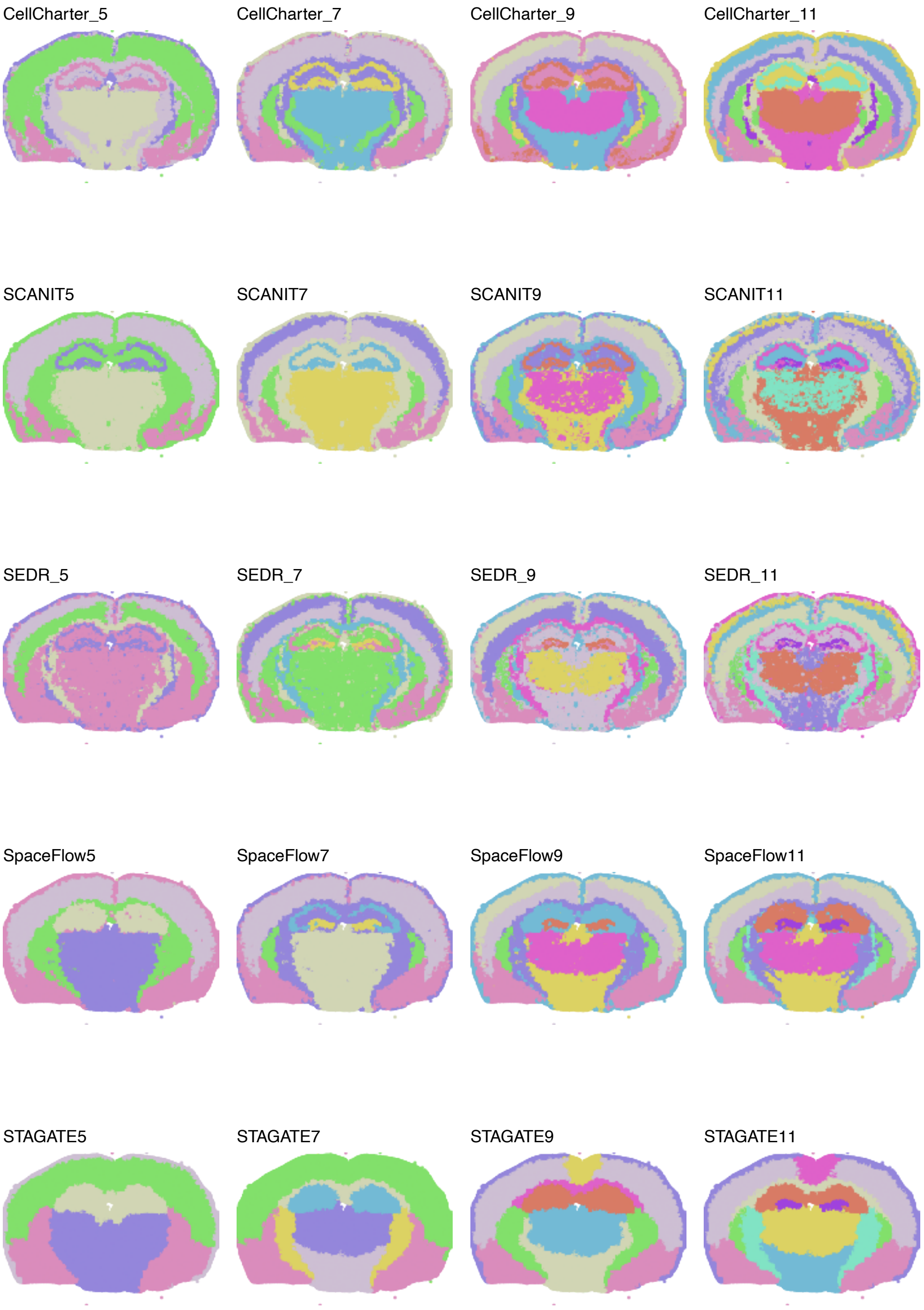


**Figure S8**: Spatial plots of Xenium mouse brain dataset cluster sweep results.

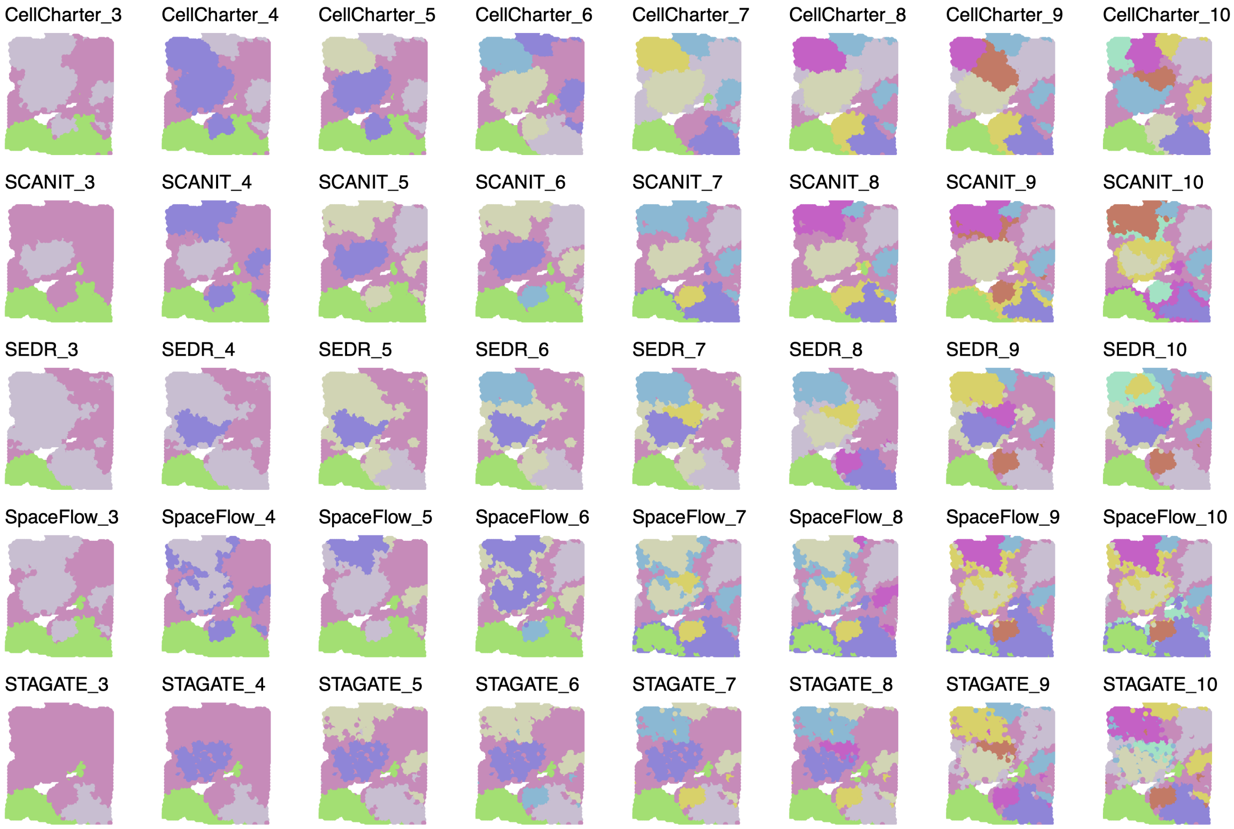


**Figure S9**: Spatial plots of Visium breast cancer dataset cluster sweep results.

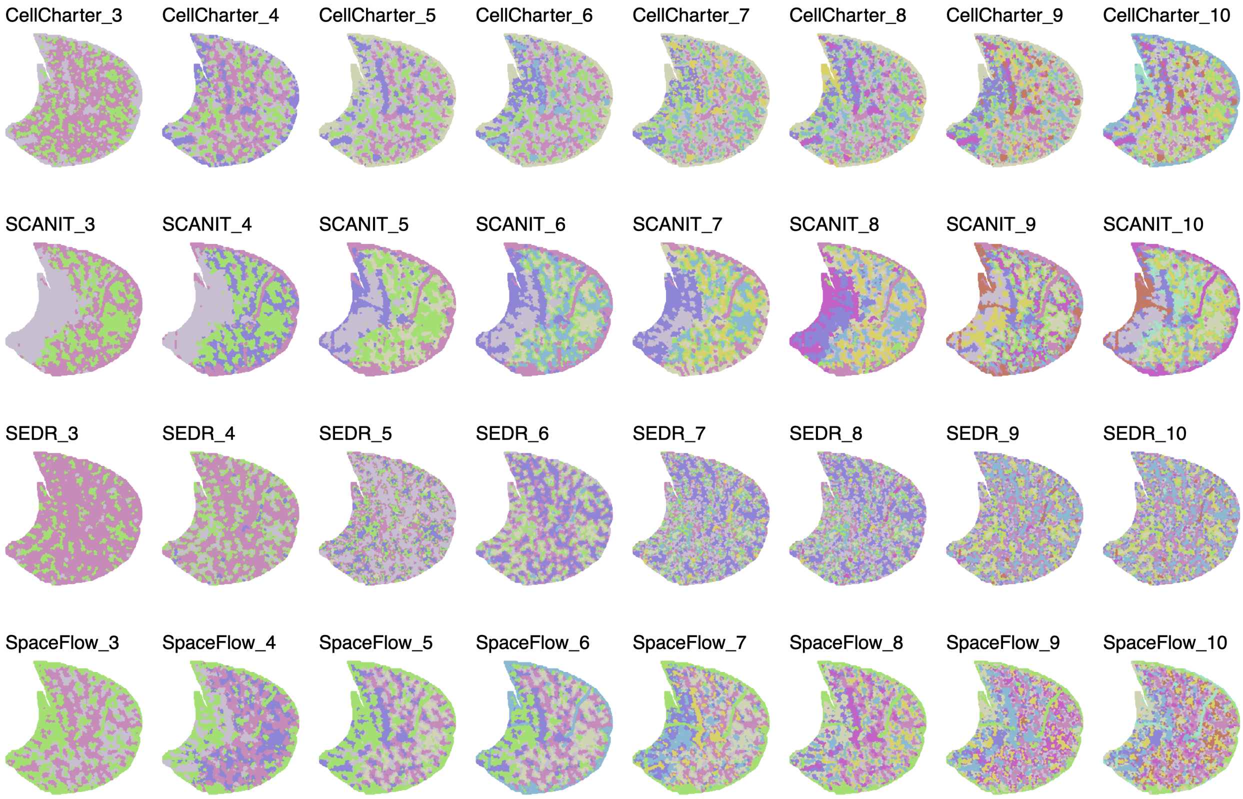


**Figure S10**: Spatial plots of Stereo-seq mouse liver dataset cluster sweep results with all genes.

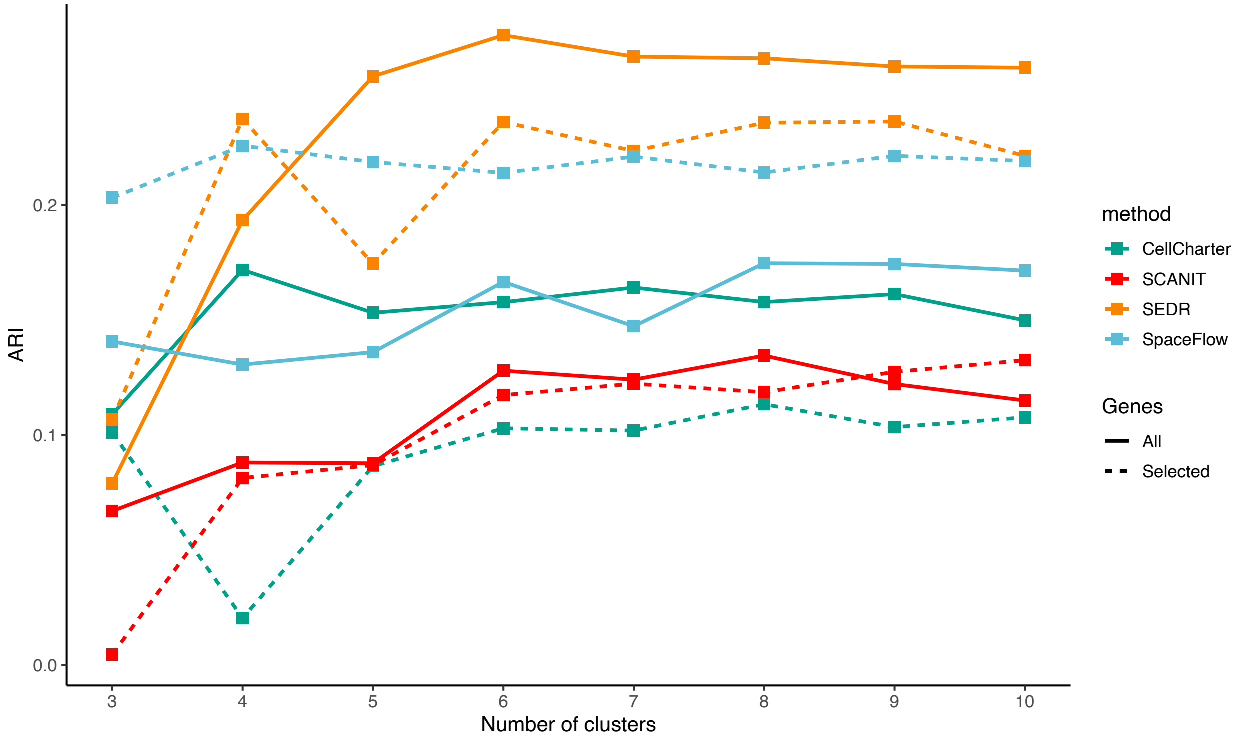


**Figure S11**: ARI results of Stereo-seq mouse liver datasets with all genes or selected marker genes.

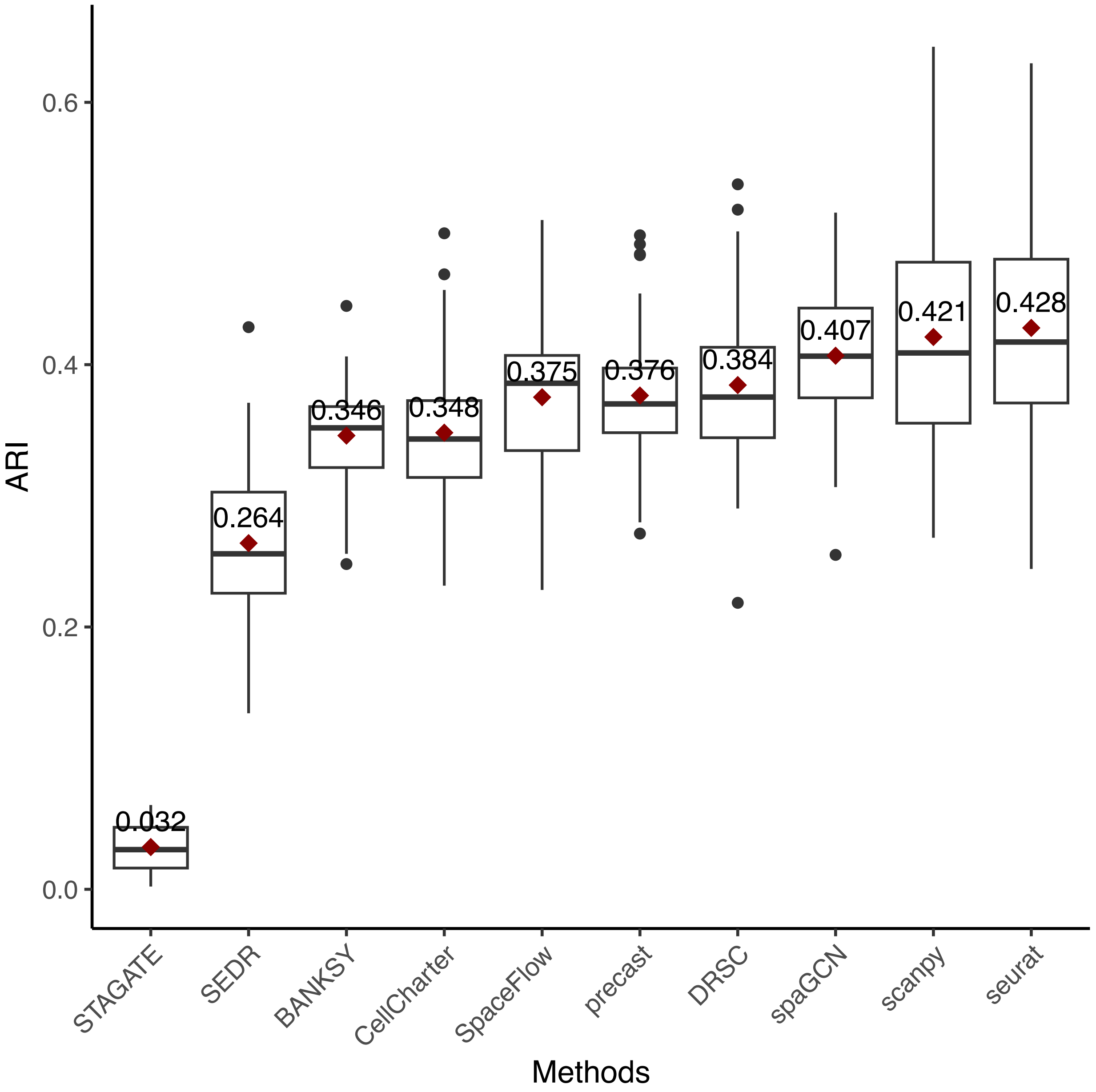


**Figure S12**: ARI results for Stereo-seq mouse embryo dataset, with mean value indicated in red square and corresponding values.

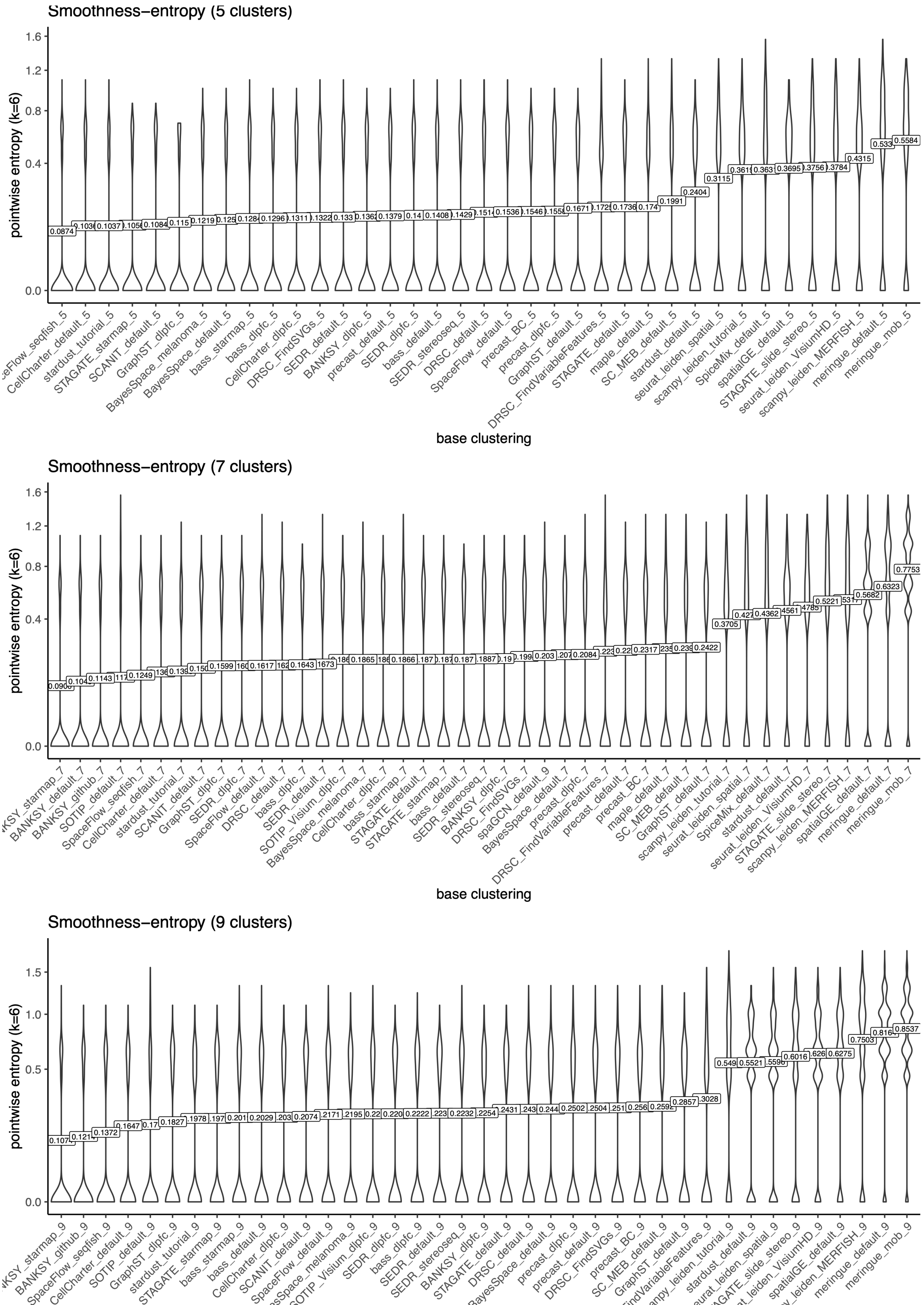


**Figure S13**: Distributions of SE scores for all base clusterings of LIBD DLPFC slice 151673, with mean value indicated in the text boxes.

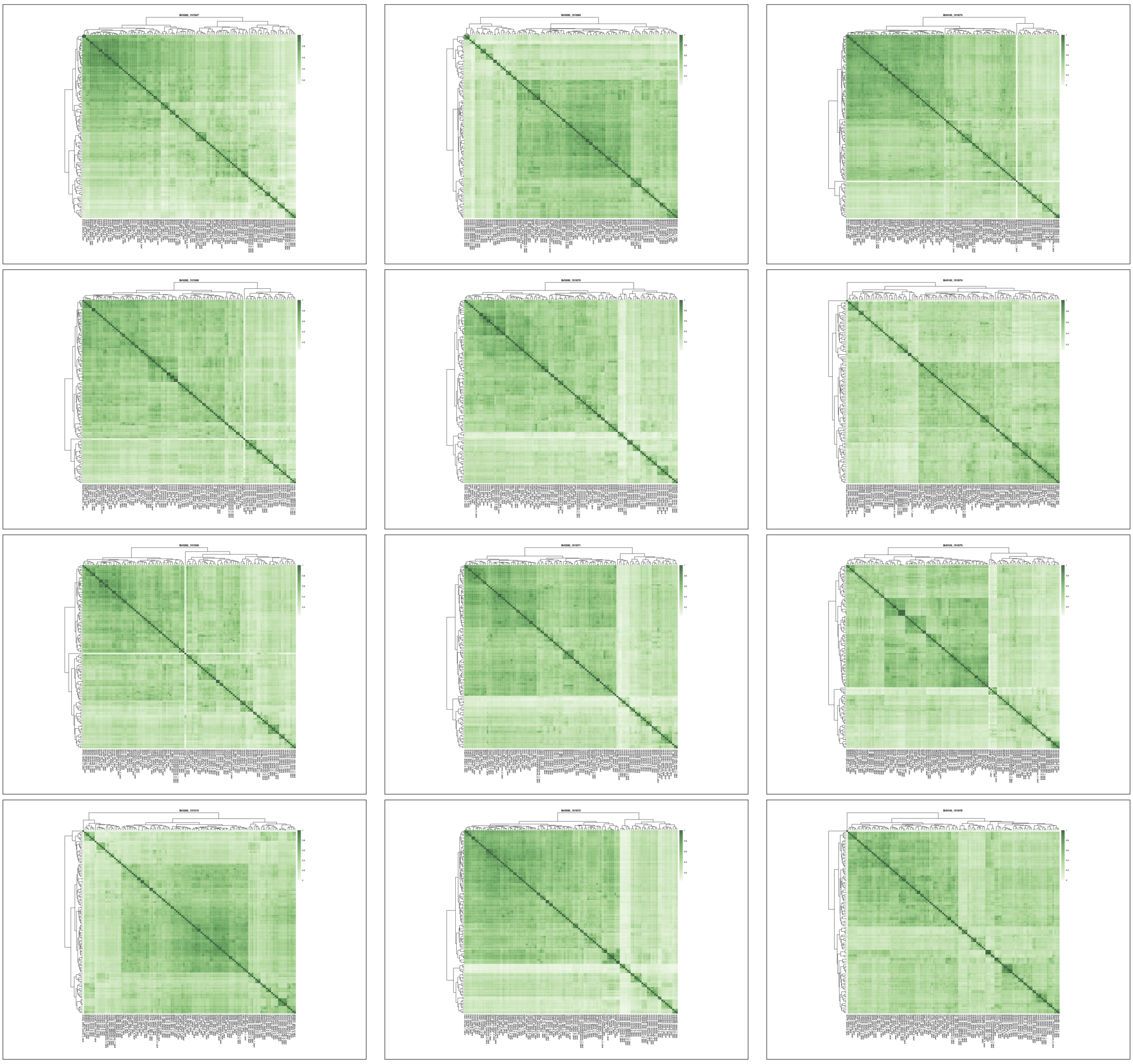

**Figure S14**: Heatmaps of pairwise ARIs between all base clusterings for each LIBD DLPFC slice. In all cases, we observed a block of concordant methods.

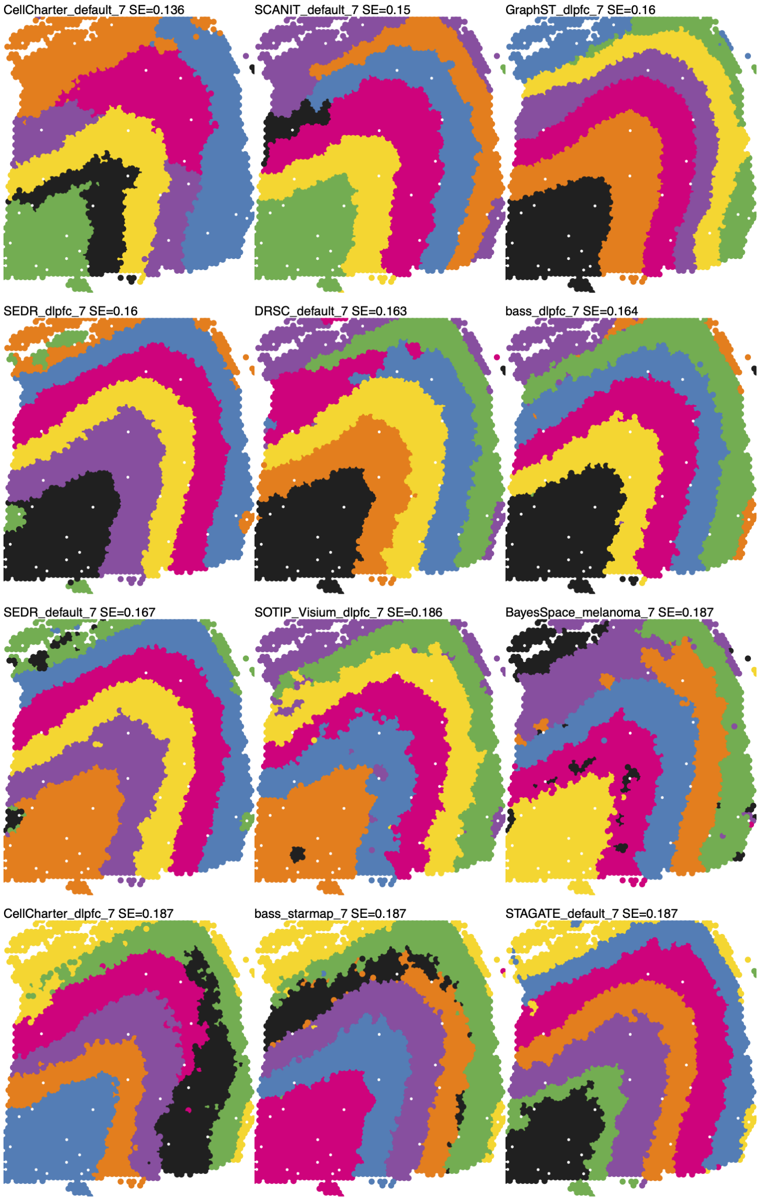


**Figure S15**: 12 selected base clusterings at 7 clusters to form consensus used for Figure 3B ("consensus_lca", and represented in Figure 3D) for LIBD DLPFC slice 151673.

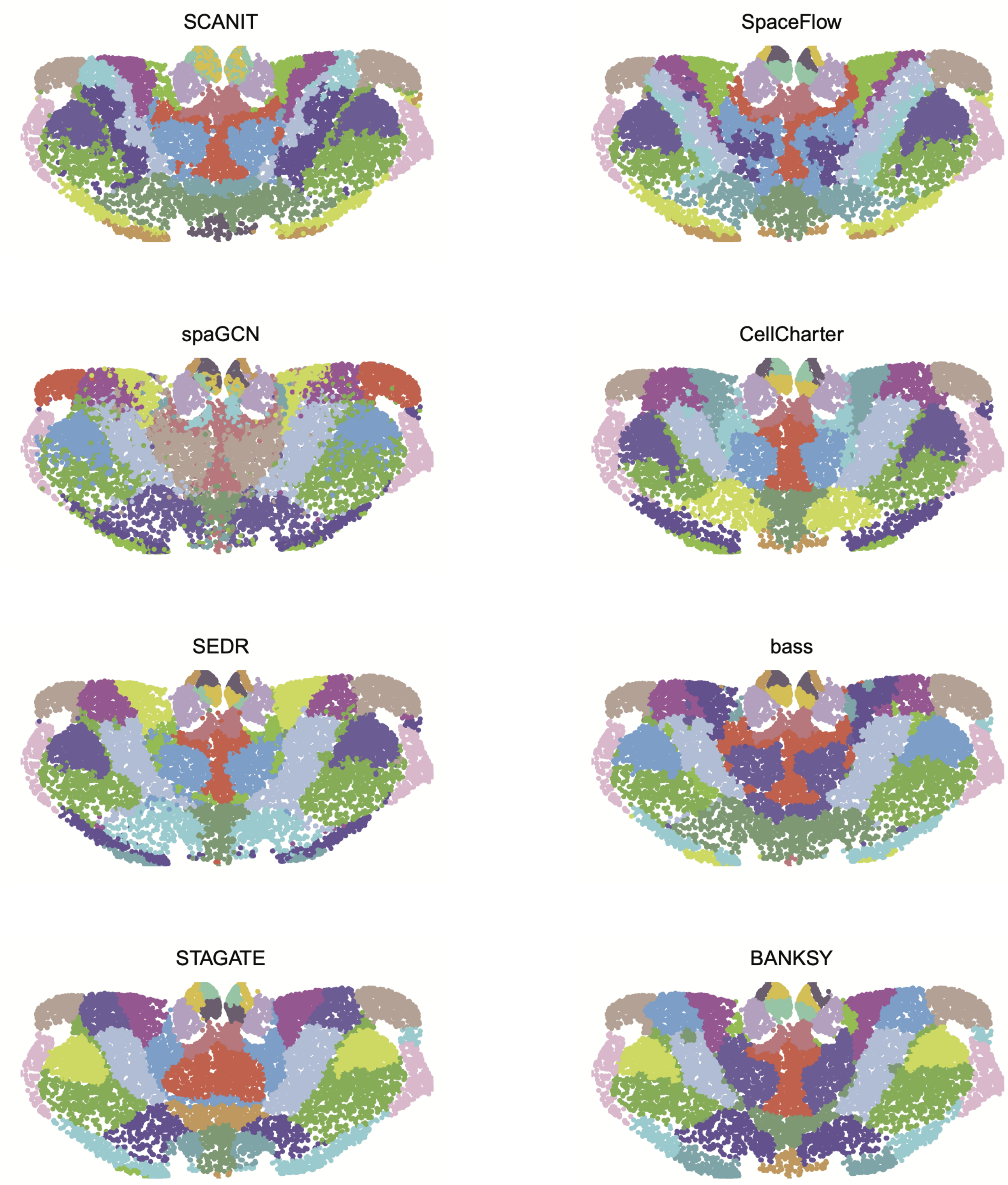


**Figure S16**: Clusters in the mouse thalamus generated from running 8 SAC methods with the cluster parameter set to 20.

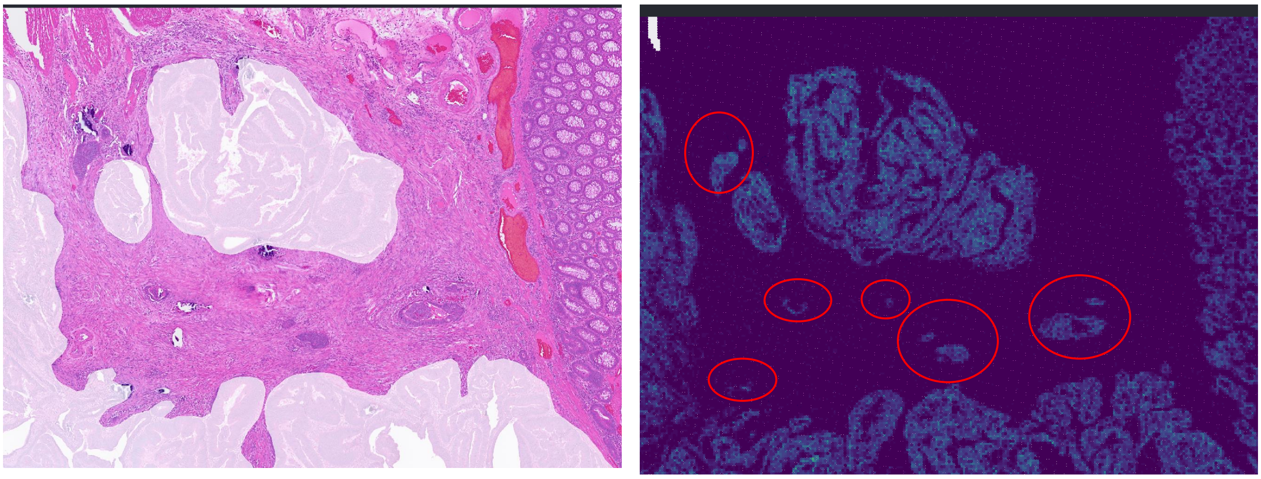

**Additional tumor areas added based on *EPCAM* expression**

**Figure S17**: *EPCAM* expression used for fine annotation of the neoplasm. Circles in red were initially missed based solely on the HE

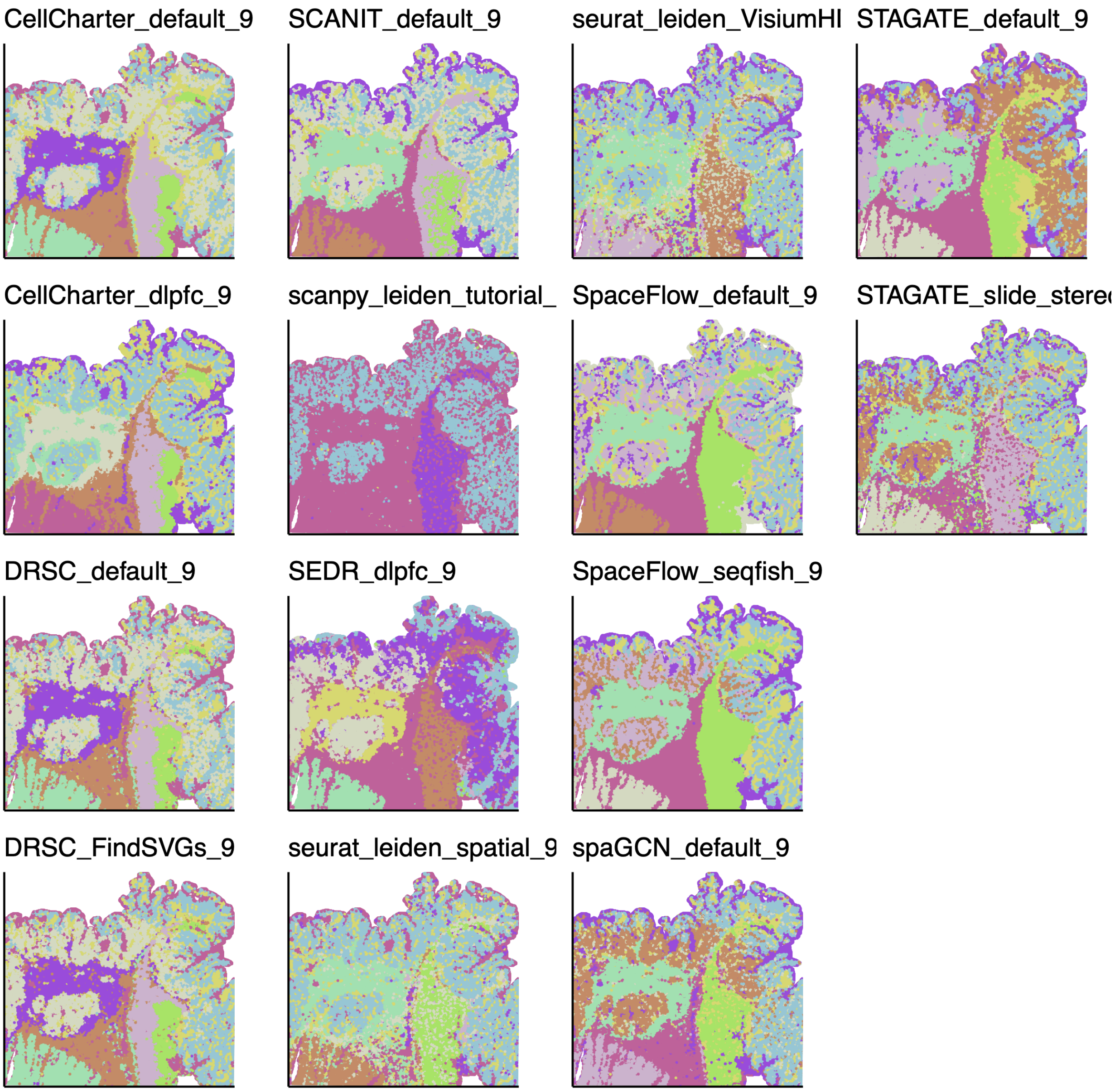


**Figure S18**: Base clusterings of full Visium HD cancer dataset with 9 clusters for pathologist evaluation.

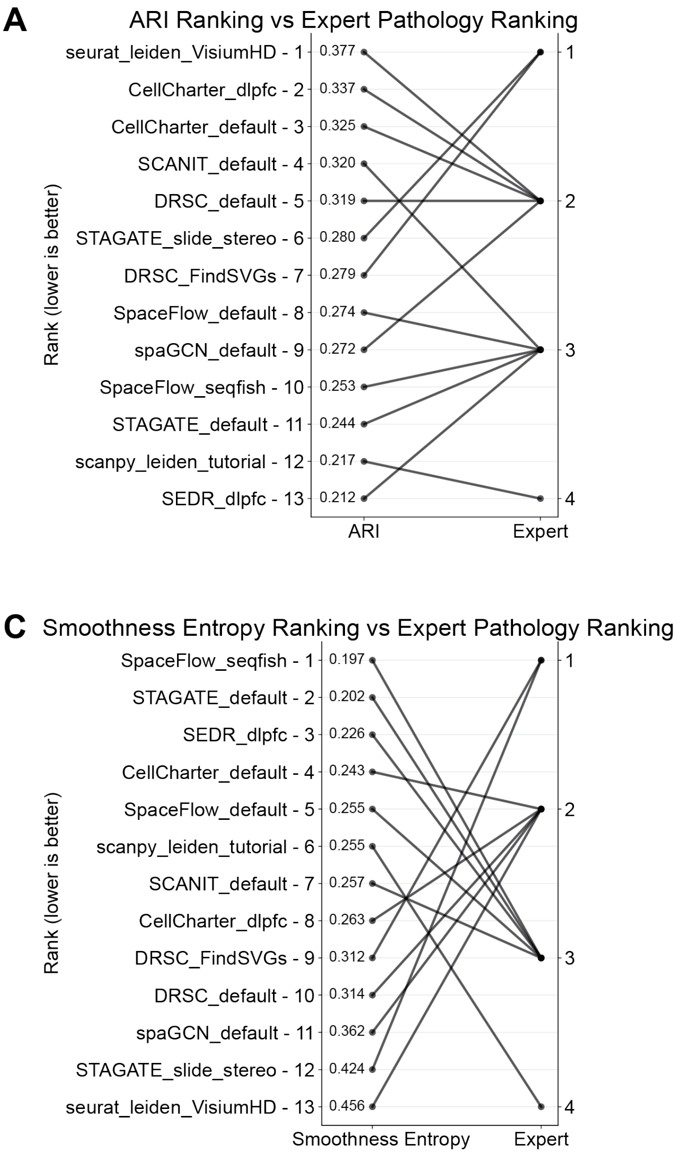


**Figure S19**: Comparison of expert rankings against evaluation metrics for the Visium HD colorectal cancer sample. Rankings according to **A**. ARI, **B**. NMI, **C**. Smoothness entropy, and **D**. CHAOS vs the expert rankings.

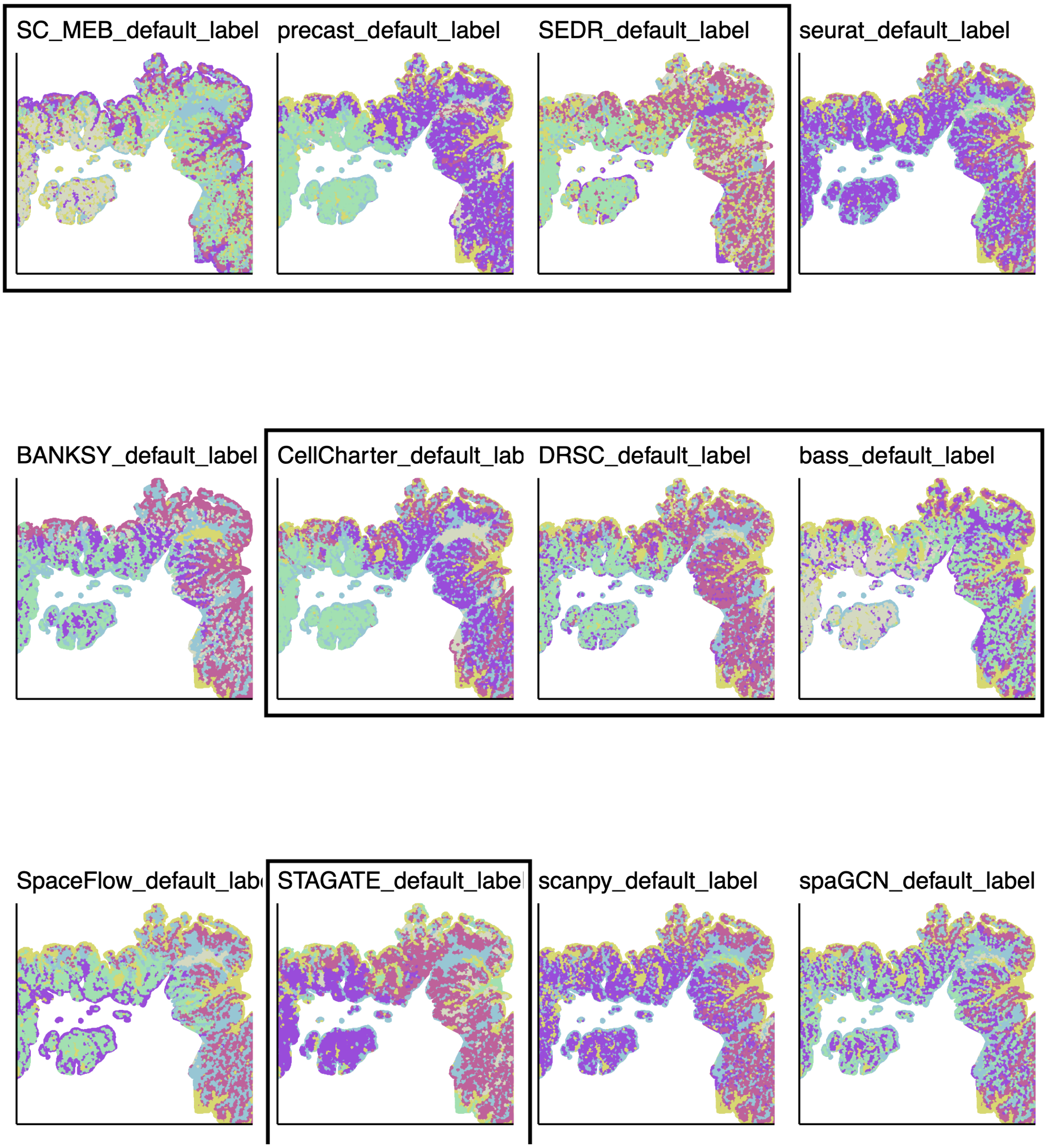


**Figure S20**: Base clusterings generated for subsetted Visium HD dataset, the instances selected for consensus are enclosed with black boxs.

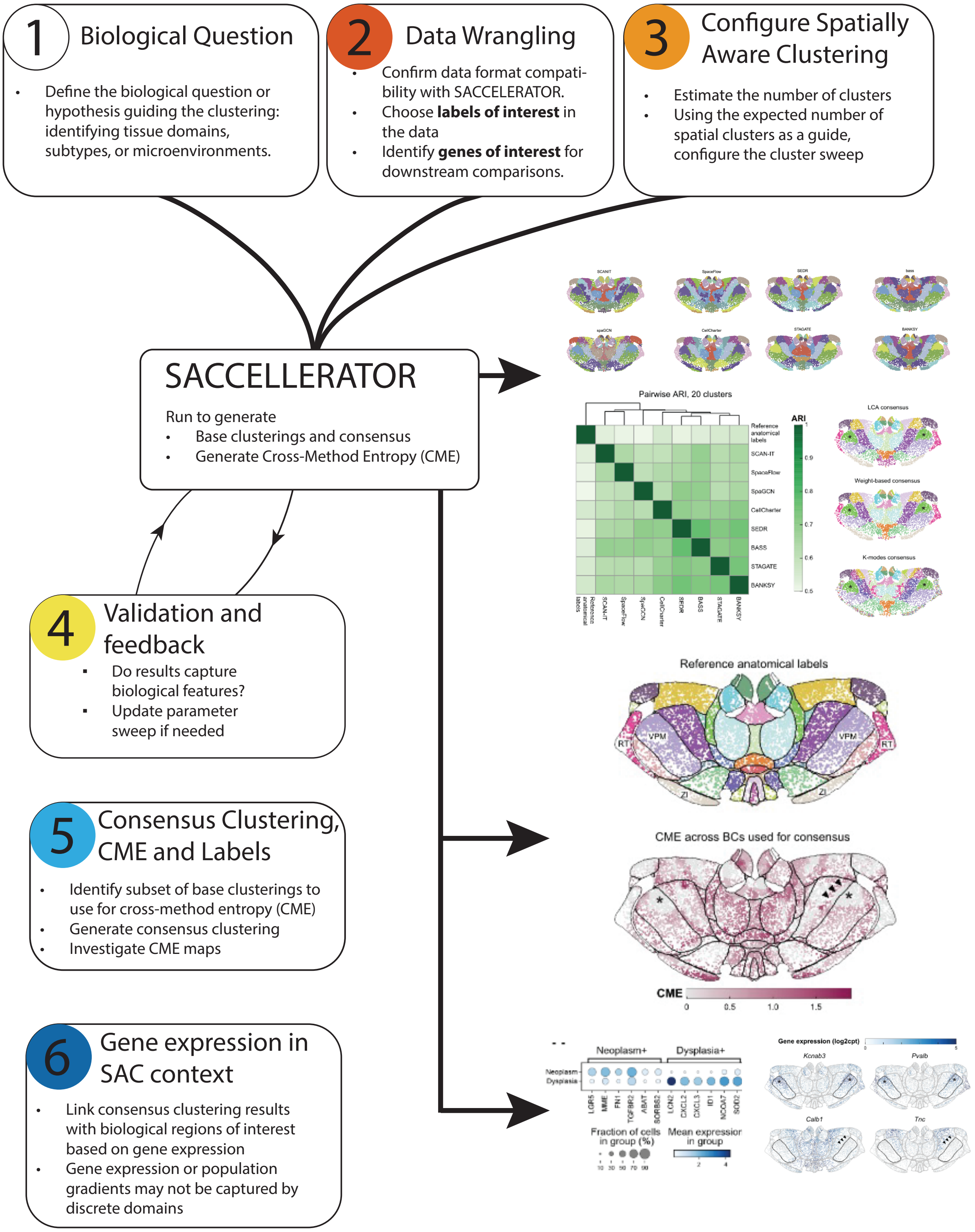


**Figure S21**: Schematic of the SACCELERATOR consensus guided expert in the loop framework as applied to analysis of the MERFISH mouse brain thalamus dataset (Case study 1). To address issues in identifying biologically relevant regions in spatial transcriptomics data, we adopt a flexible expert-in-the-loop consensus-driven approach. This goes beyond traditional ensemble/consensus methods, and allows researchers to interact with intermediate results to determine which tools should be used to generate a consensus (steps 1-3). Inclusion of an expert-in-the-loop is critical to ensure that the computational analysis matches the biological question at hand, and we believe that when the focus of the analysis is to uncover novel biological discoveries, tissue experts are accessible more often than not (steps 4-6). The figure can also be found in SACCELEATOR GitHub landing page (Code availability).

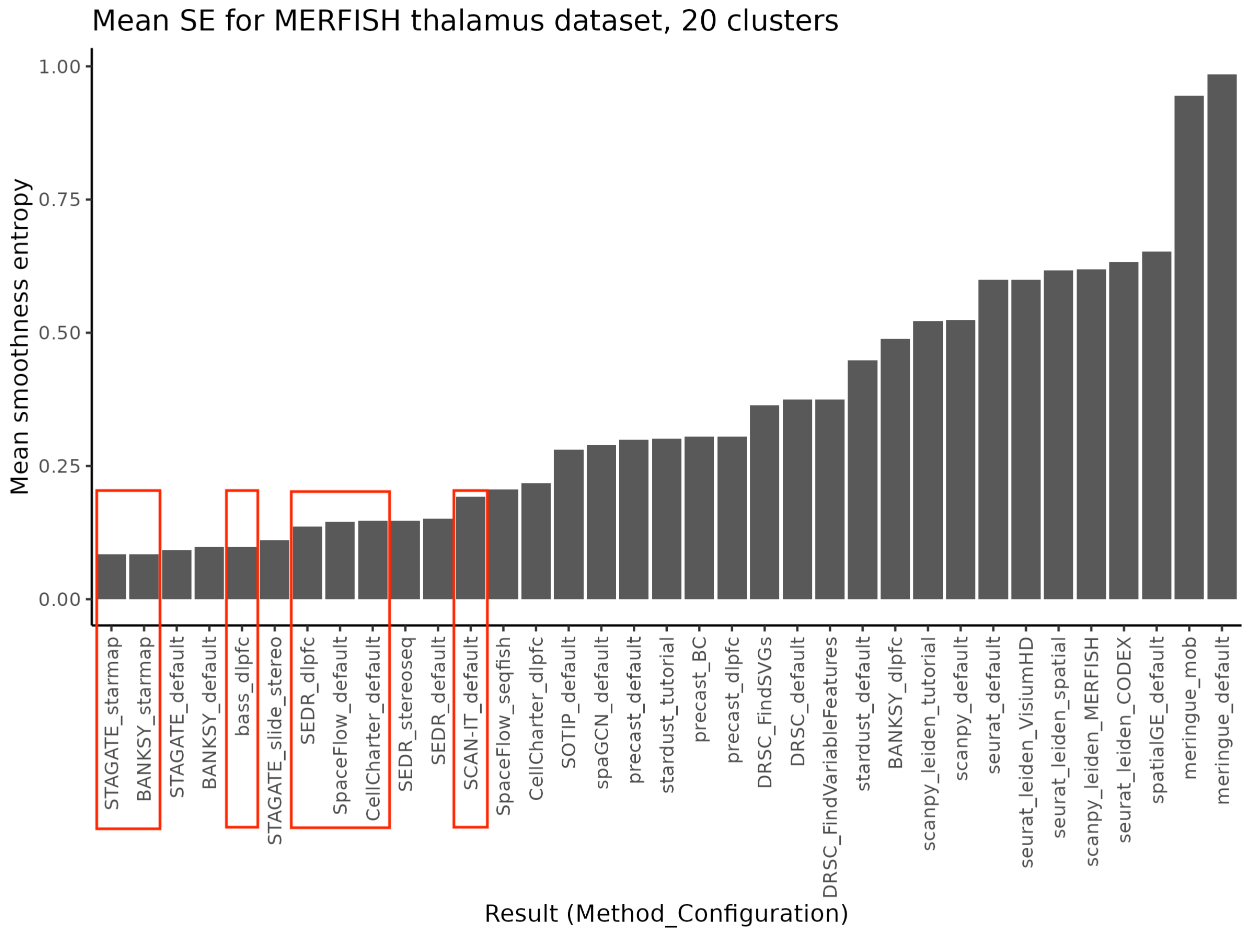


**Figure S22**: Smoothness entropy (SE) of MERFISH thalamus dataset with 20 clusters. Selected instances of base clusterings are marked by red boxes. Only the smoothness configuration of each method is selected.

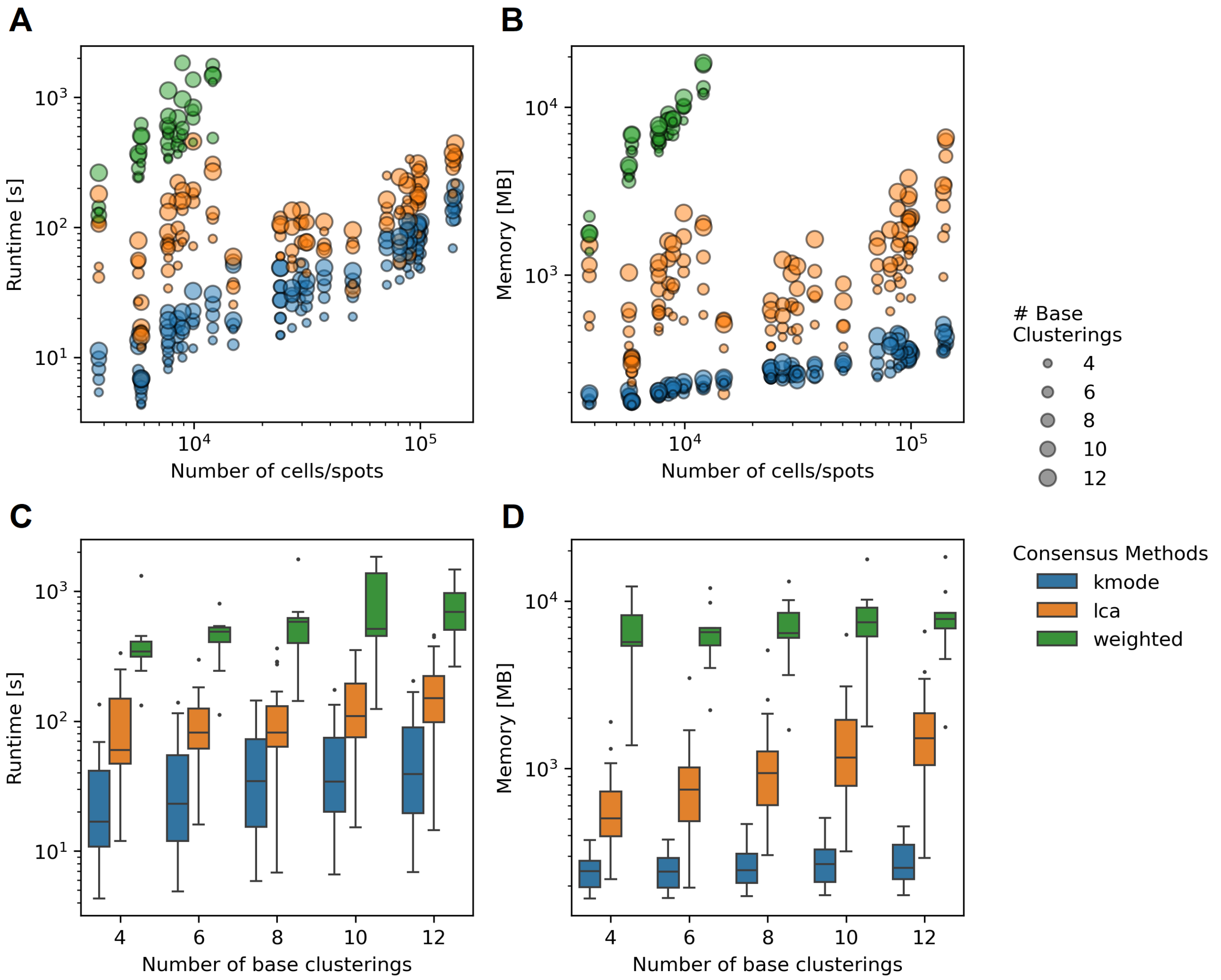


**Figure S23**: **Scalability analysis of consensus algorithms. (A, B)** Consensus algorithm runtime (A) and memory (B) w.r.t. number of cells and spots in the dataset. **(C, D)** Consensus algorithm runtime (C) and memory (B) w.r.t. number of base clustering input.

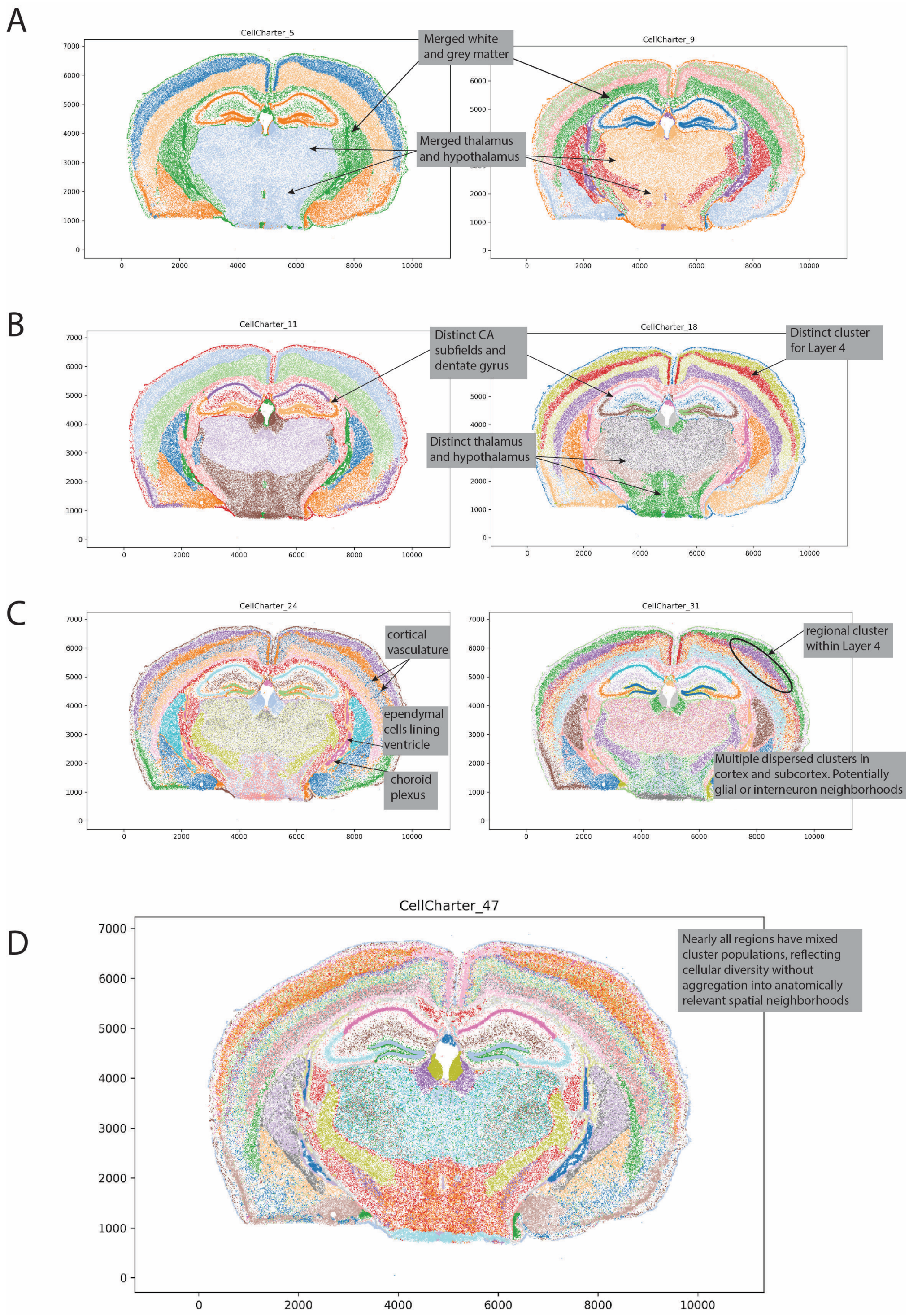


**Figure S24**: SAC cluster sweep encompasses high- and low-resolution clustering results. In this annotated example using CellCharter on the Xenium mouse brain data, we covered a wide range of cluster parameters to illustrate how spatial resolution is related to cluster number. Subplot titles indicate the number of clusters. A. Low numbers of clusters fail to resolve even coarse anatomical regions. B. Moderate cluster numbers include highly resolved anatomical structures, including cortical layers, structures within the hippocampus and distinct domains for thalamus and hypothalamus. C. High cluster numbers showing further resolution, which may be desirable for some studies, but some anatomical regions begin to include multiple clusters built on distinct cellular populations. D. Very high cluster number showing salt-and-pepper distributions of clusters across the brain, reflecting cellular diversity rather than contiguous spatial domains.

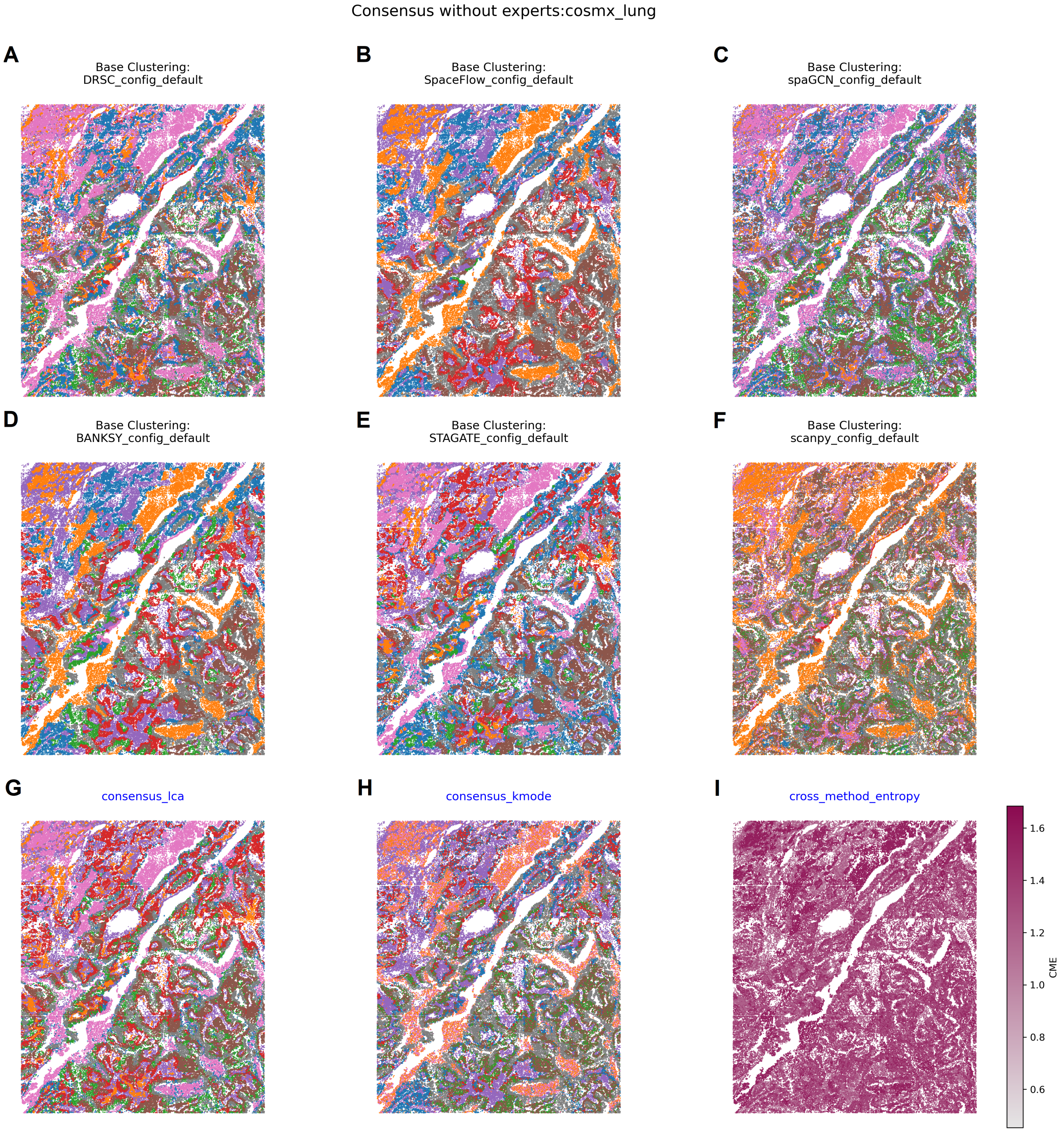


**Figure S25**: Base and consensus clusterings for the CosMX lung dataset. **(A-F)** Base clustering from methods using default configurations set to 8 clusters. **(G-H)** Consensus clustering using LCA and K-modes consensus building. Results for weighted are not shown as the computation did not complete. **(I)** Cross method entropy heatmap.

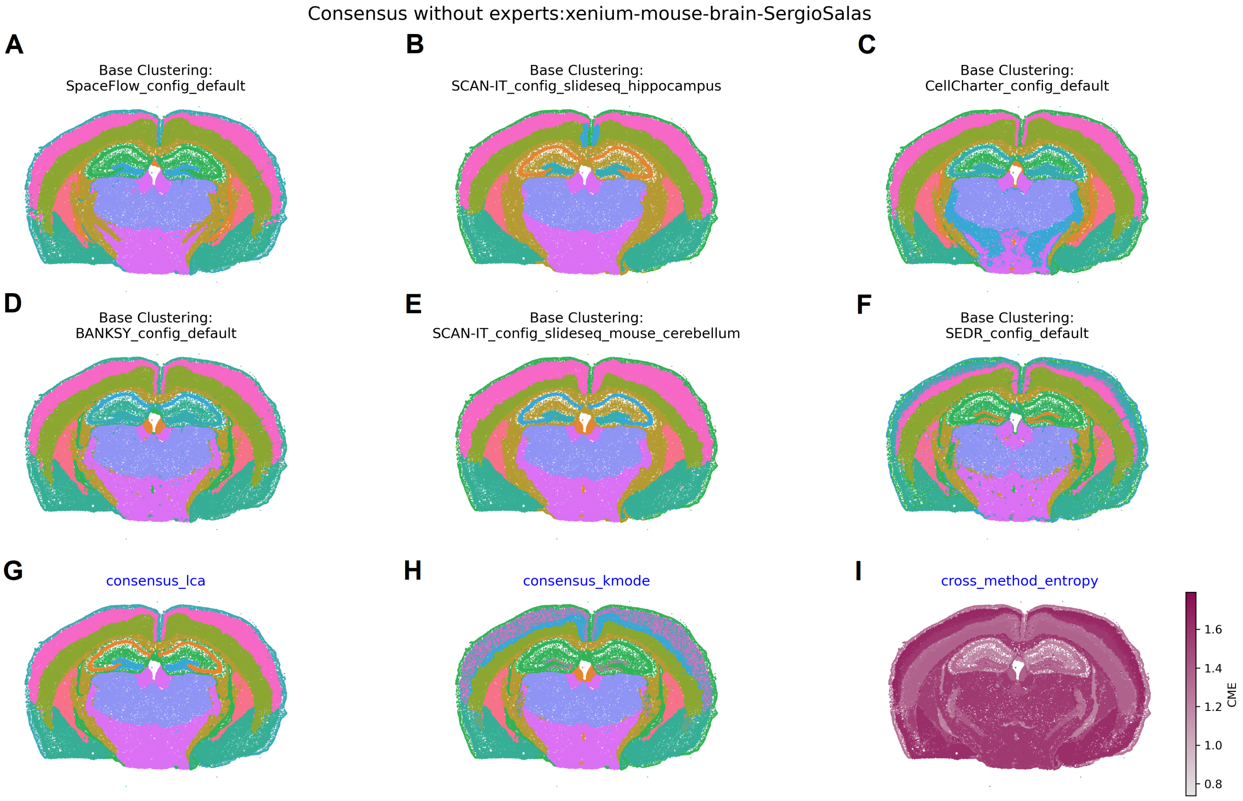


**Figure S26**: Base and consensus clusterings for the Xenium mouse brain dataset. **(A-F)** Base clustering from methods using default configurations set to 11 clusters. **(G-H)** Consensus clustering using LCA and K-modes consensus building. Results for weighted are not shown as the computation did not complete. **(I)** Cross method entropy heatmap.

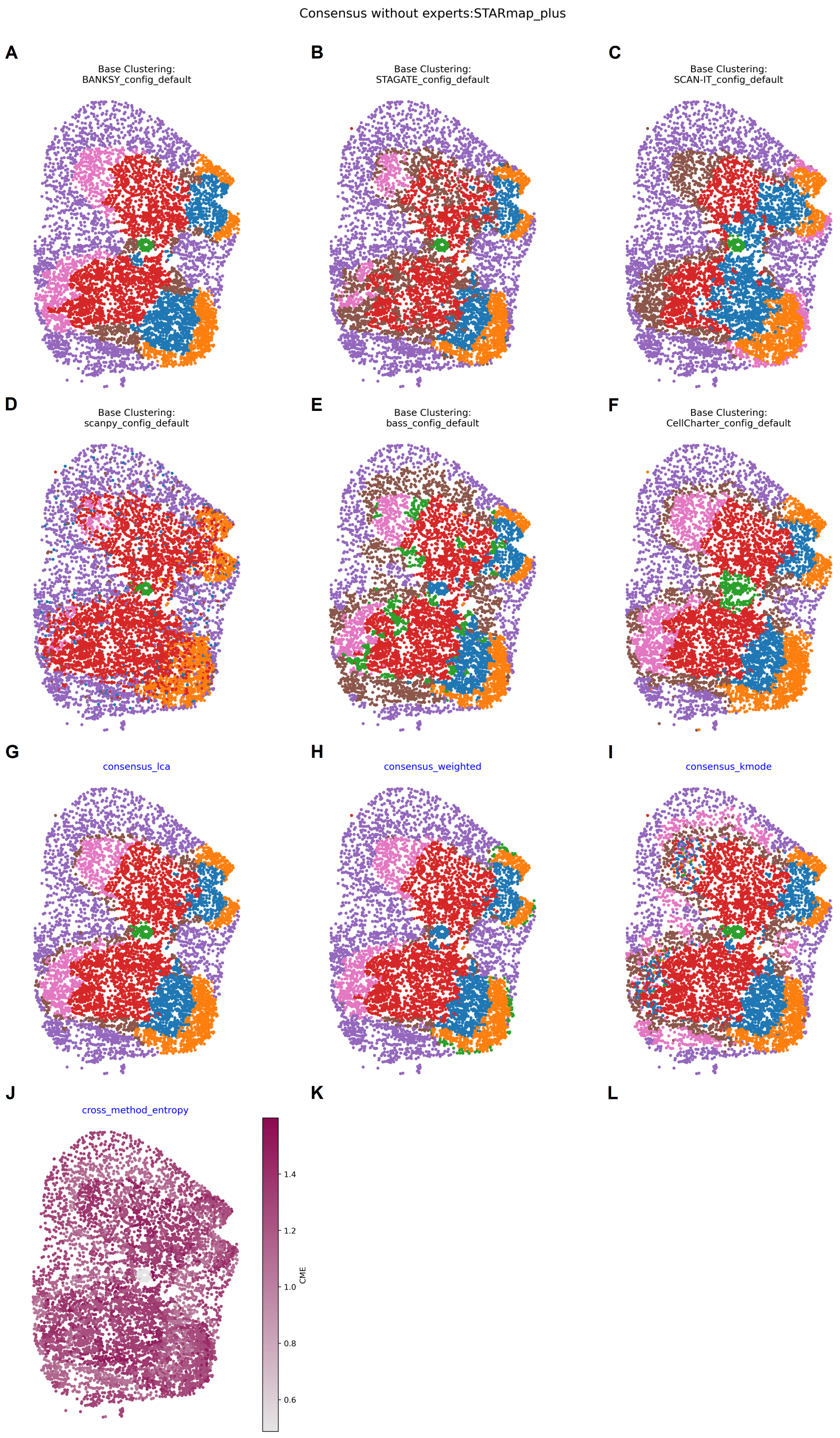


**Figure S27**: Base and consensus clusterings for the STARmap plus mouse brain dataset. **(A-F)** Base clustering from methods using default configurations set to 7 clusters. **(G-I)** Consensus clustering using LCA, K-modes and weighted consensus building. **(J)** Cross method entropy heatmap.

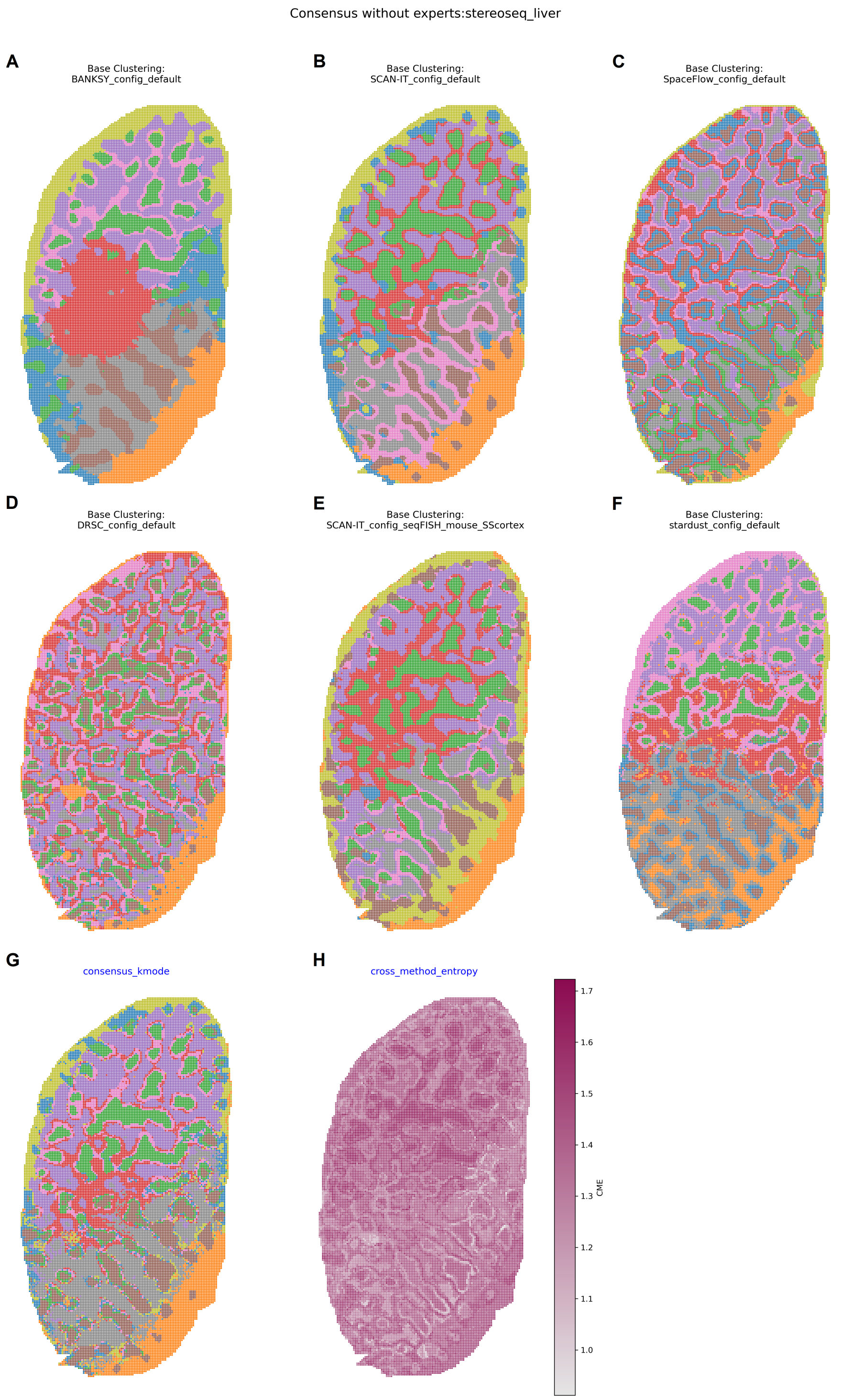


**Figure S28**: Base and consensus clusterings for the Stero-seq liver dataset. **(A-F)** Base clustering from methods using default configurations set to 9 clusters. **(G)** Consensus clustering using K-modes. Results for LCA and weighted are not shown as the computation did not complete. **(H)** Cross method entropy heatmap.

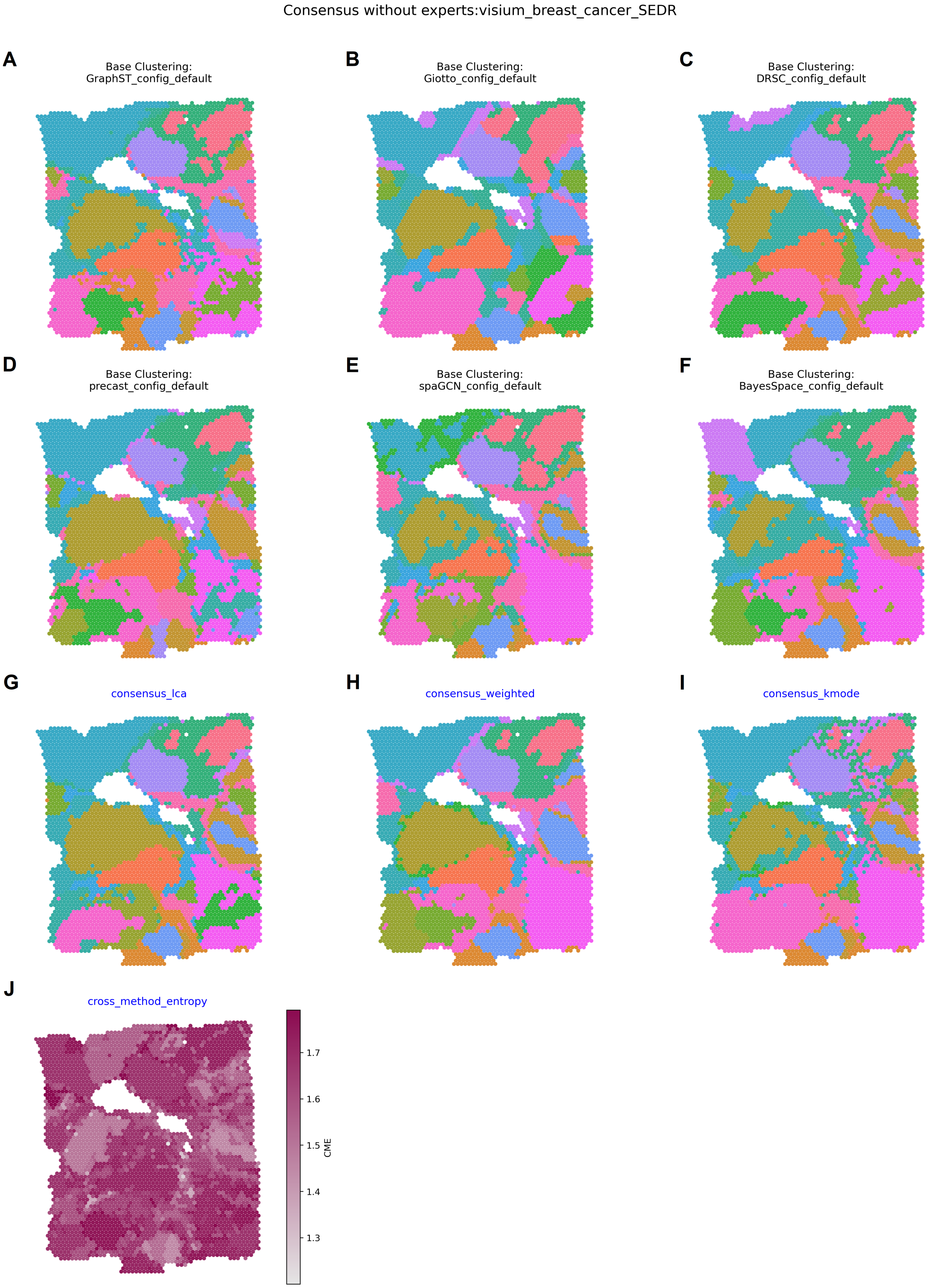


**Figure S29**: Base and consensus clusterings for the Visum breast cancer dataset. **(A-F)** Base clustering from methods using default configurations set to 20 clusters. **(G-I)** Consensus clustering using LCA, K-modes and weighted consensus building. **(J)** Cross method entropy heatmap.

#### Supplementary Tables

**Table S1**: Overview of methods. A GitHub commit is only included for methods installed from GitHub without releases.

| Name | Version (commit) | Year | Language | Citation |
|---|---|---|---|---|
| MAPLE | 0.99.1 (b173e89) | 2024 | R | Jeon et al. (2024) |
| BANKSY | 0.99.7 (dbda6fd) | 2023 | R (+ Python) | Singhal et al. (2023) |
| PRECAST | 1.6.3 | 2023 | R | Liu et al. (2023) |
| GraphST | 1.1.1 | 2023 | Python | Long et al. (2023) |
| SPICEMIX | 0.0.71 | 2023 | Python | Chidester et al. (2023) |
| CellCharter | 0.2.0 | 2023 | Python | Varrone et al. (2023) |
| STAGATE | 1.0.1 (48ce7f8) | 2022 | Python | Dong & Zhang (2022) |
| scMEB | 1.1 | 2022 | R | Yang et al. (2022) |
| SpaceFlow | 1.0.4 | 2022 | Python | Ren et al. (2022) |
| DR-SC | 3.3 | 2022 | R | Liu et al. (2022) |
| SCAN-IT | 0.1 (ebf3894) | 2022 | Python | Cang et al. (2022) |
| SOTIP | 1.0 (d3b762c) | 2022 | Python | Yuan et al. (2022) |
| Stardust | 2.0 (f1b5417) | 2022 | R | Avesani et al. (2022) |
| BASS | 1.1.0.016 (37980c9) | 2022 | R | Li & Zhou (2022) |
| spatialGE | 1.2.0.0000 | 2022 | R | Ospina et al. (2022) |
| SpaGCN | 1.2.7 | 2021 | Python | Hu et al. (2021) |
| BayesSpace | 1.10.1 | 2021 | R | Zhao et al. (2021) |
| SEDR | 1.0.0 (8c273af) | 2021 | Python | Fu et al. (2021) |
| MERINGUE | 1.0 (ca9e2cc) | 2021 | R | Miller et al. (2021) |
| Giotto | 4.0.0 | 2021 | R (+ Python) | Dries et al. (2021) |
| Scanpy | 1.10.0 | 2018 | Python | Wolf et al. (2018) |
| Seurat | 5.0.1 | 2018 | R | Butler et al. (2018) |

**Table S2**: Overview of method configurations.

**Table S3**: Overview of metrics.

| Name | Requires GT labels | Uses spatial information | Language | Citation |
|---|---|---|---|---|
| Adjusted Rand Index (ARI) | yes | no | Python | Hubert & Arabie (1985) |
| Spatial chaos score (CHAOS) | no | yes | R | Shang & Zhou (2022) |
| Calinski–Harabasz index | yes | no | Python | Calinski & Harabasz (1974) |
| Completeness | yes | no | Python | Rosenberg & Hirschberg (2007) |
| Davies–Bouldin index | no | no | Python | Davies & Bouldin (1979) |
| Entropy | yes | no | Python | Shannon (1948) |
| Fowlkes–Mallows Index (FMI) | yes | no | Python | Fowlkes & Mallows (1983) |
| Homogeneity | yes | no | Python | Rosenberg & Hirschberg (2007) |
| Local Inverse Simpson's Index (LISI) | yes | no | R | Korsunsky et al. (2019) |
| Mathew's Correlation Coefficient (MCC) | yes | no | Python | Chicco & Jurman (2020) |
| Normalized Mutual Information (NMI) | yes | no | R | Strehl & Ghosh (2002) |
| Percentage of Abnormal Spots (PAS) | no | yes | R | Shang & Zhou (2022) |
| SpatialARI | yes | yes | R | Luo et al. (2025) |
| V measure | yes | no | Python | Rosenberg & Hirschberg (2007) |
| Cluster specific silhouette | no | no | R | Rousseeuw (1987) |
| Domain specific F1 | yes | no | R | Rijsbergen (1979) |
| Jaccard index | yes | no | Python | Jaccard (1912) |

**Table S4**: SpaceHack 2.0 participants.
