## Supplementary File 1 for "Beyond benchmarking: an expert-guided consensus approach to spatially aware clustering"

| | |
|---|---|
| **Biological Question** | • **Define the biological question** or hypothesis guiding the clustering: identifying tissue domains, subtypes, or microenvironments. For example:<br>○ Does spatial transcriptomics support subdividing anatomical regions of the thalamus?<br>○ Can spatial transcriptomics identify neoplastic progression stages in colorectal cancer? |
| **Data Wrangling** | • Confirm **data format compatibility** with SACCELERATOR.<br>• Identify **labels of interest** already in the data: manual annotations and cell type labels can be used for comparison and validation.<br>• Identify **genes of interest** for downstream comparisons. |
| **Configure Spatially Aware Clustering** | • **Estimate** how many spatial clusters are present in the tissue<br>• Using the **expected number of spatial clusters** as a guide, configure the cluster sweep to span at least ±5 numbers of clusters around the expected number.<br>• **Run** all, or selected methods suitable for the data. By default the downstream consensus workflow will select the 8 most concordant methods based on cross-method ARI. This will generate the base clusterings |
| **Validation and Feedback** | • **Review SACCELERATOR outputs**: and compare to biologically relevant structures.<br>○ Validate scale and number of clusters in the context of the biological topic.<br>○ Does one base clustering result address biological questions?<br>○ For example, in the brain: do some outputs include laminar organization of cortex? In cancer cases, do some outputs correctly identify tumor tissue?<br>• If biological features are not included, **review SACCELLERATOR configuration** and broaden the parameter sweep. |
| **Consensus Clustering, CME and Reference Labels** | • If needed, identify the **subset of base clusterings** to use for cross-method entropy (CME), e.g., filtering base clusterings based on spatial continuity measures (such as smoothness entropy).<br>• Generate **visual summaries** (spatial consensus plot, cross-method entropy map, weight distribution plot, pairwise ARI heatmaps with clustering method class) to accompany expert interpretation.<br>• Using CME heatmap and consensus, check if any single base **clustering method/class dominates the consensus**.<br>• Verify that spatial resolution and granularity of the **final consensus** includes the biological scales of interest.<br>• **Interpretation of CME maps:**<br>○ Identify low entropy areas, where labels are consistent across methods.<br>○ Areas of high entropy often correspond to boundaries between regions where multiple methods give slightly different results. |
| **Gene Expression in SAC context** | • **Review gene expression** in the context of consensus clustering.<br>○ Link **consensus clustering results with biological regions of interest** based on gene expression.<br>○ Gene expression or population **gradients may not be captured by discrete domains**.<br>○ Do genes of interest show differential expression between consensus clusters? |
| **Important Notes** | • SACCELLERATOR assumes cellular **quality control** has already been performed.<br>• Even within a single tissue sample, genes of interest, relevant length scales, and expected numbers of clusters can **vary based on biological questions**. Independent runs of SACCELERATOR may be required to address different biological questions.<br>• We recommend a discussion to **review interpretations** of the consensus clusterings, including, for example: choice of a single consensus output or application of CME to identify domain boundaries. |
