## Supplementary File 2 for "Beyond benchmarking: an expert-guided consensus approach to spatially aware clustering"

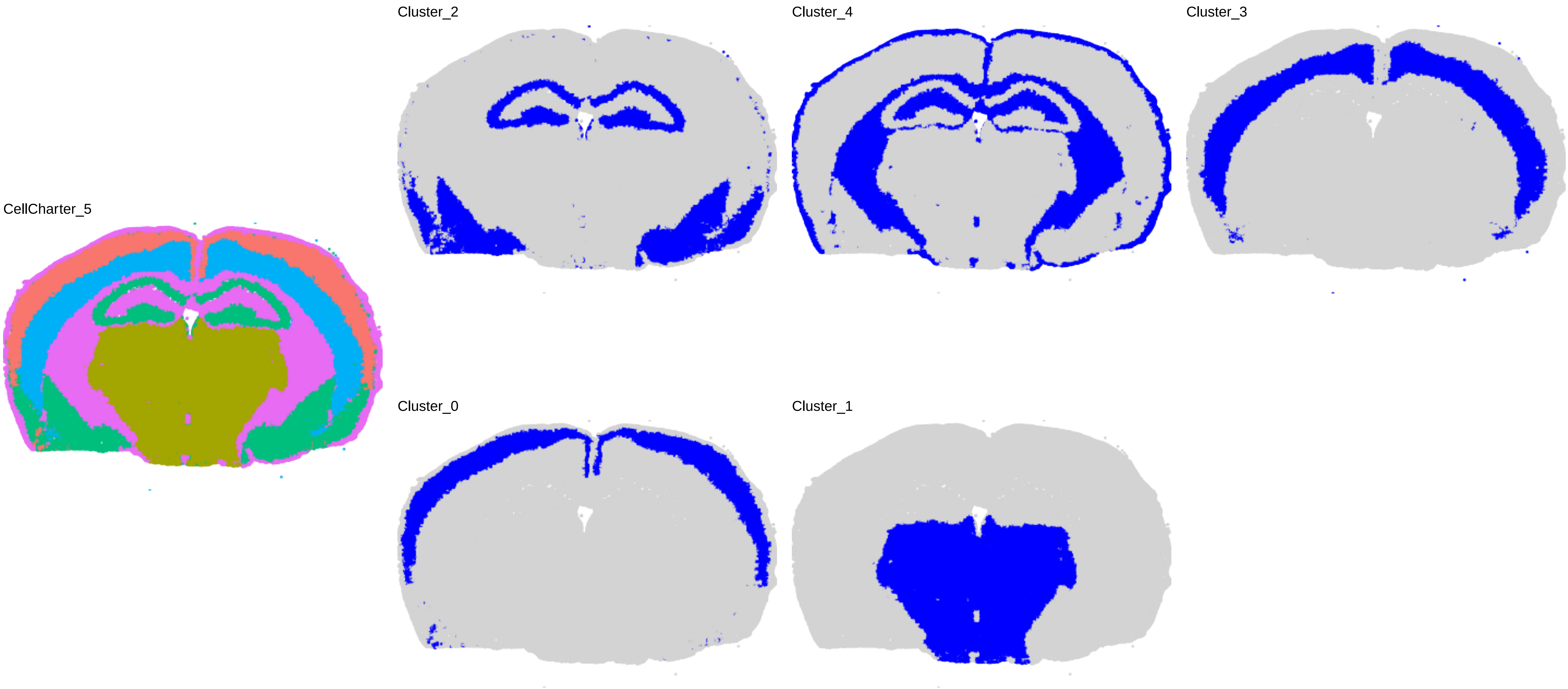

CellCharter_5
Cluster_2
Cluster_4
Cluster_3
Cluster_0
Cluster_1

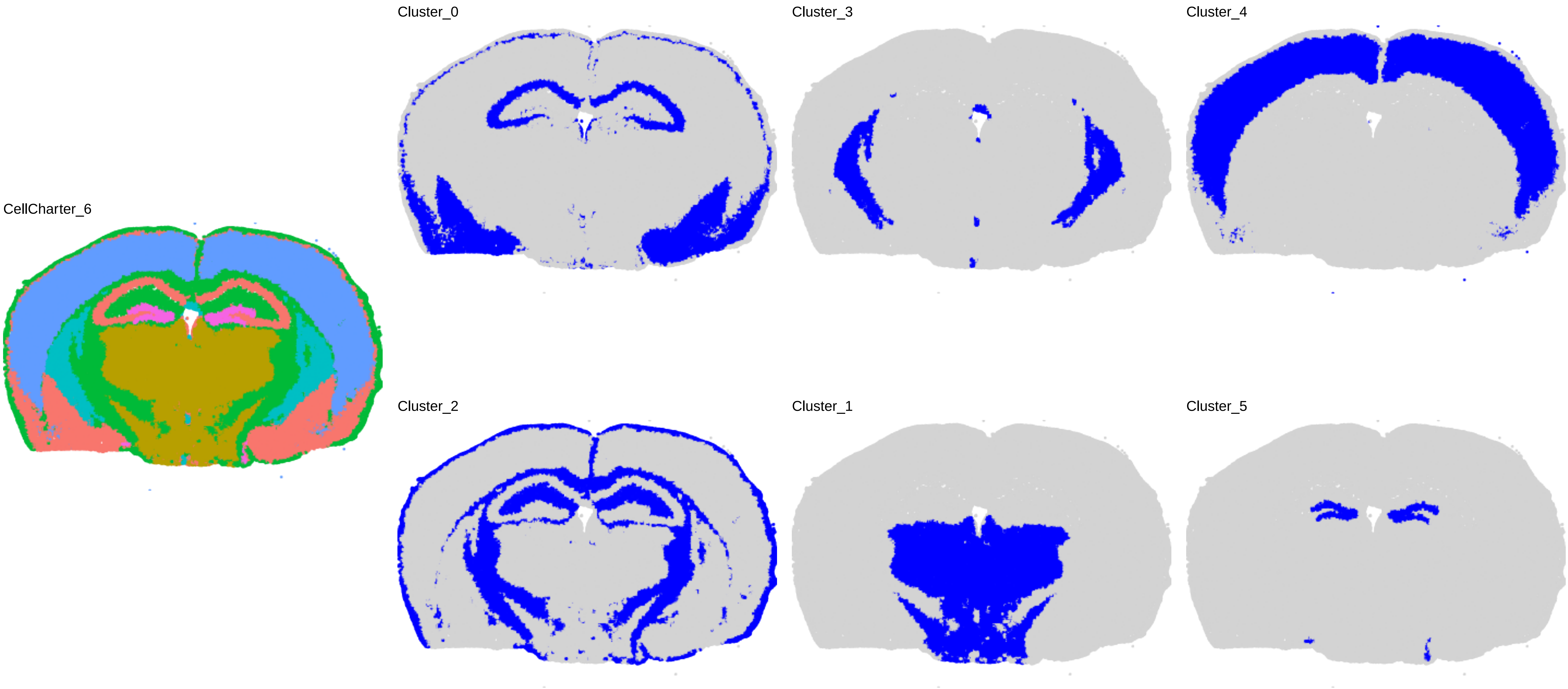
CellCharter_6
Cluster_0
Cluster_3
Cluster_4
Cluster_2
Cluster_1
Cluster_5

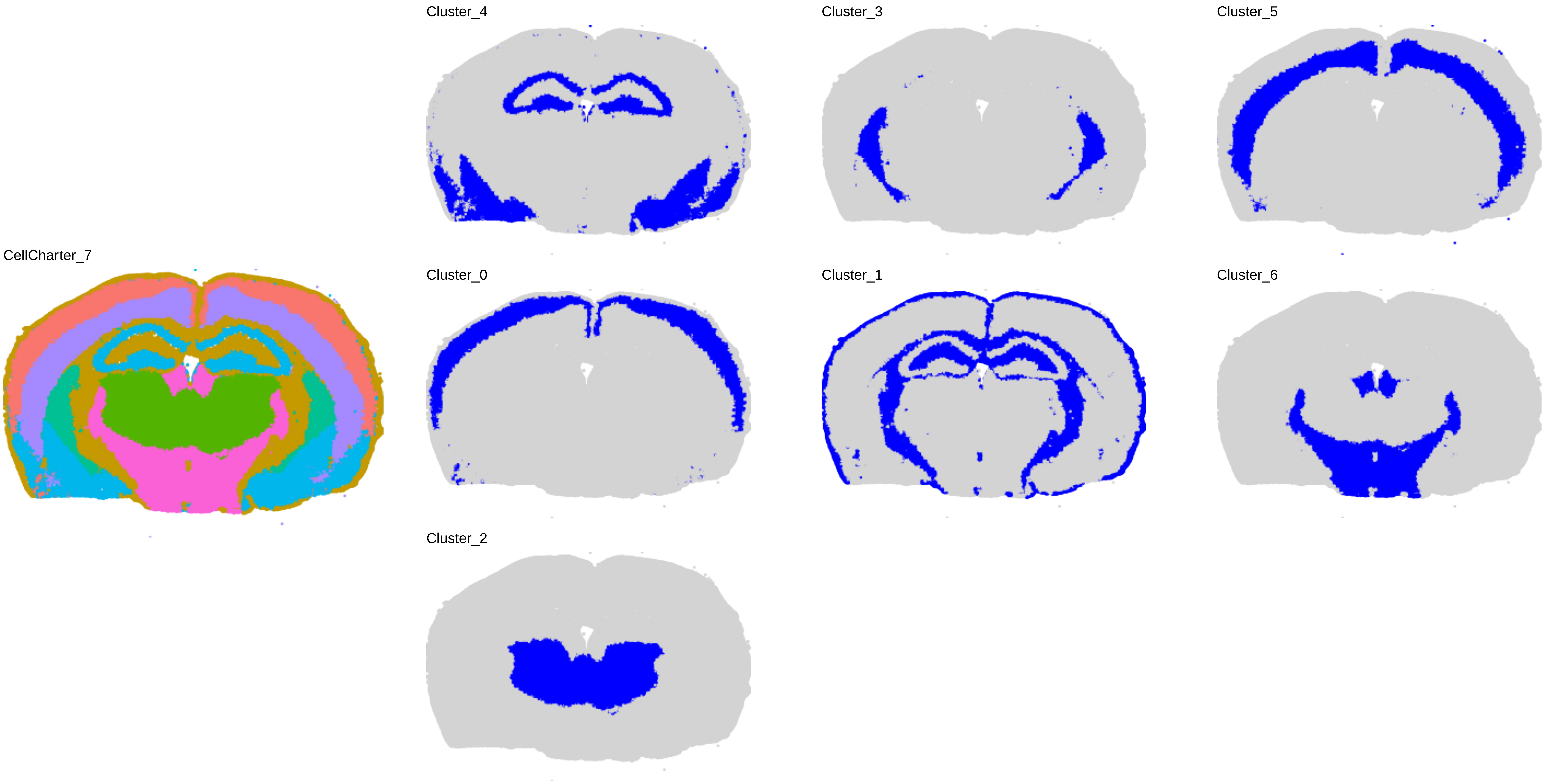

Cluster_4
Cluster_3
Cluster_5
CellCharter_7
Cluster_0
Cluster_1
Cluster_6
Cluster_2

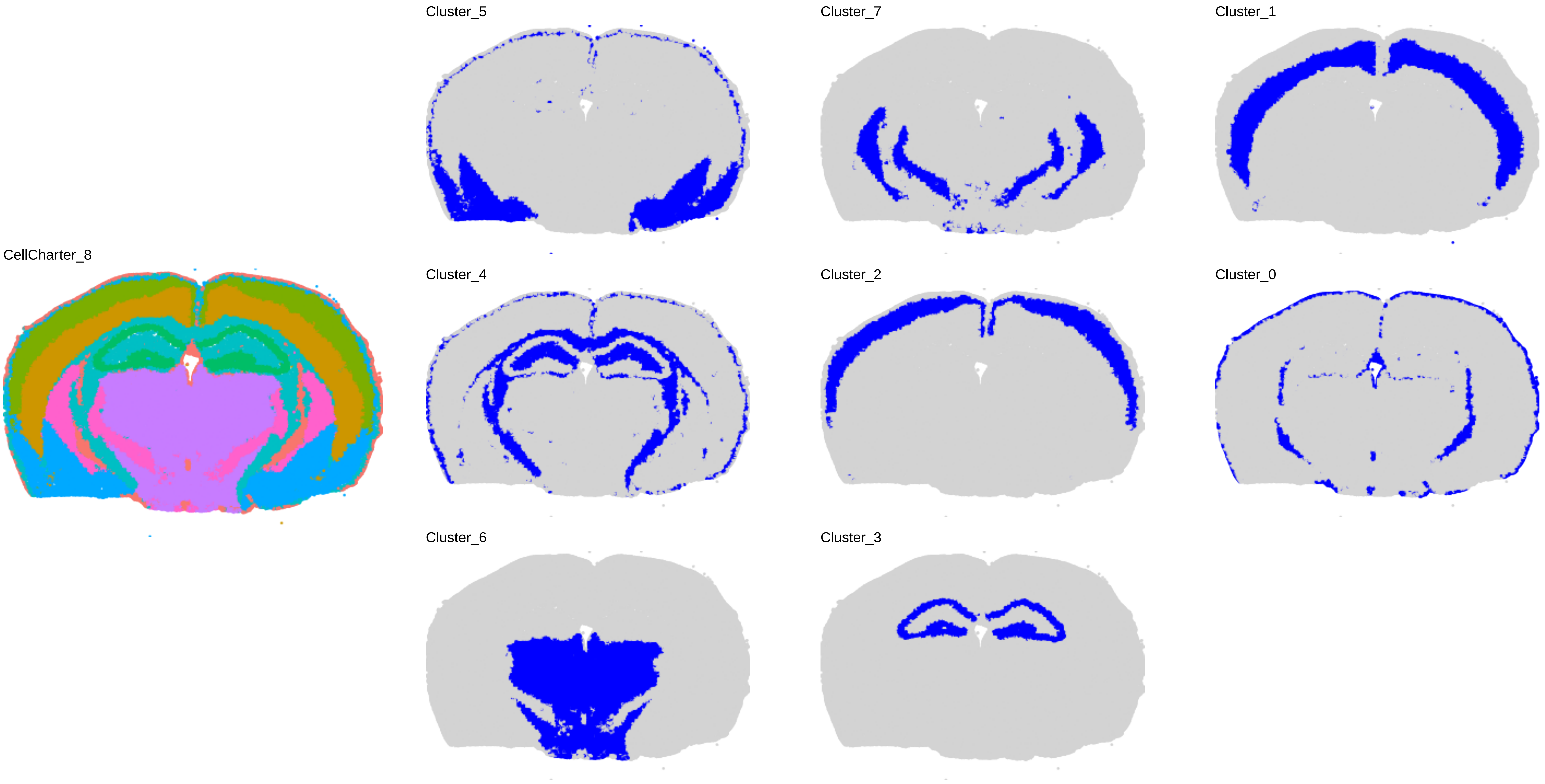
CellCharter_8
Cluster_5
Cluster_7
Cluster_1
Cluster_4
Cluster_2
Cluster_0
Cluster_6
Cluster_3

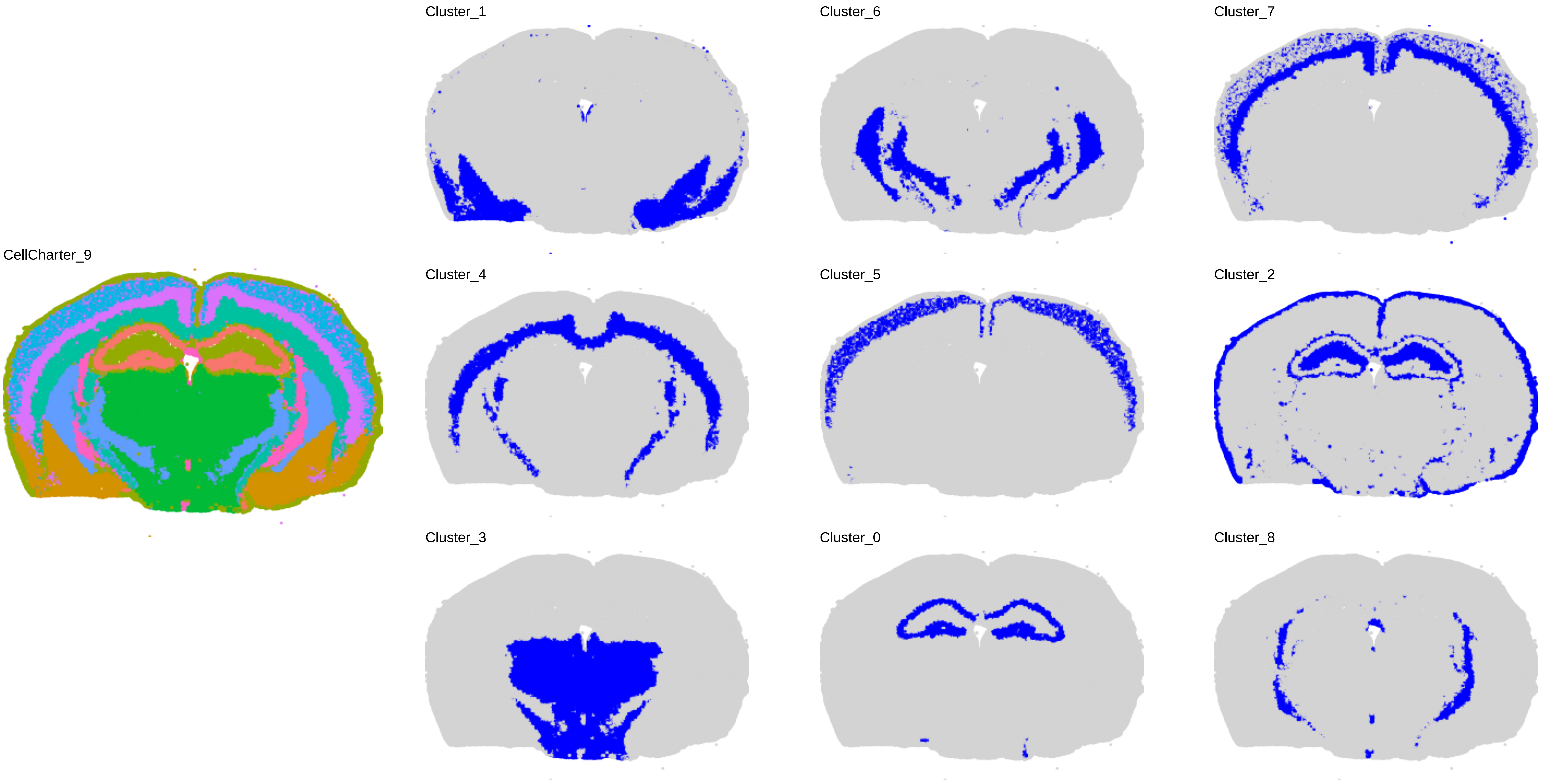
CellCharter_9
Cluster_1
Cluster_6
Cluster_7
Cluster_4
Cluster_5
Cluster_2
Cluster_3
Cluster_0
Cluster_8

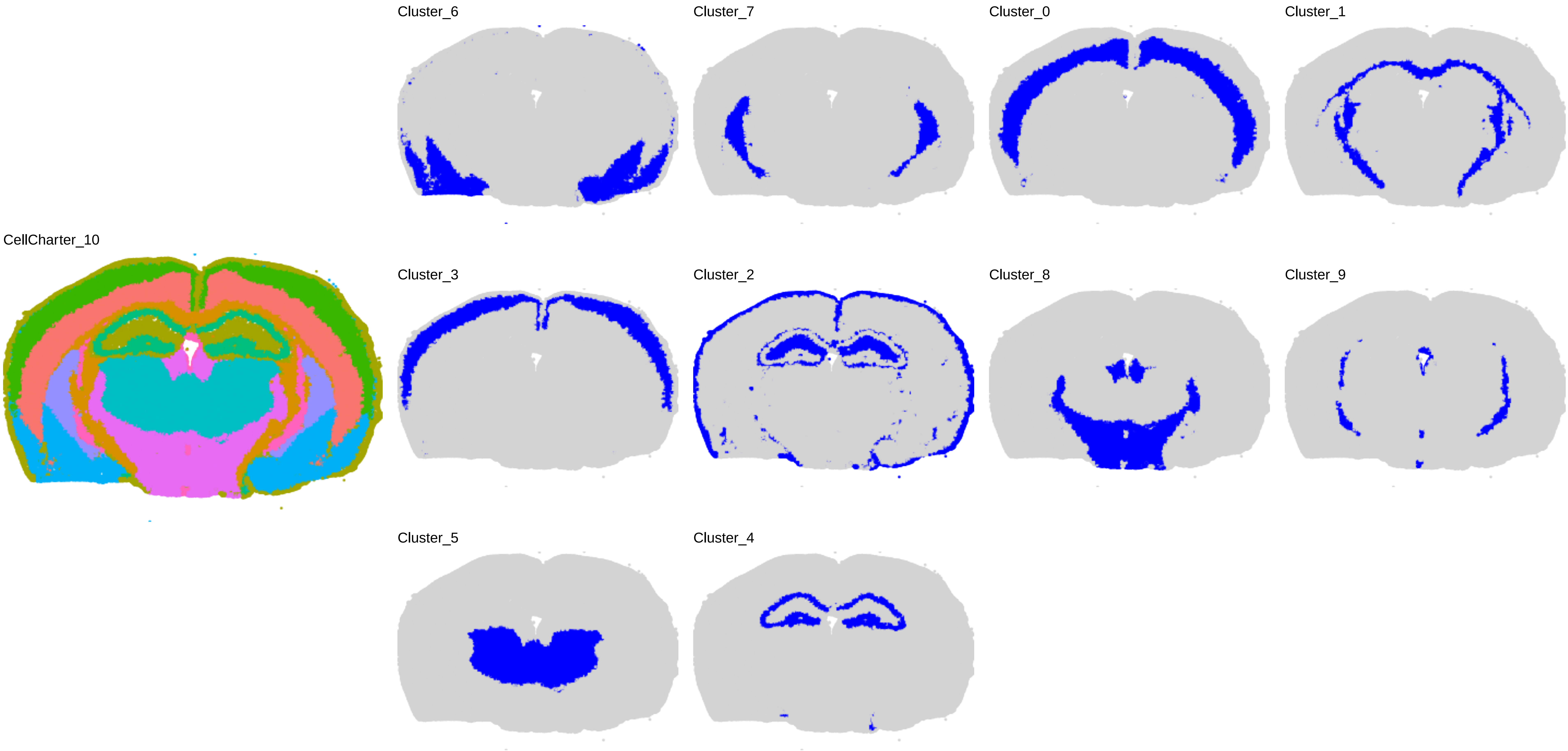
CellCharter_10
Cluster_6
Cluster_7
Cluster_0
Cluster_1
Cluster_3
Cluster_2
Cluster_8
Cluster_9
Cluster_5
Cluster_4

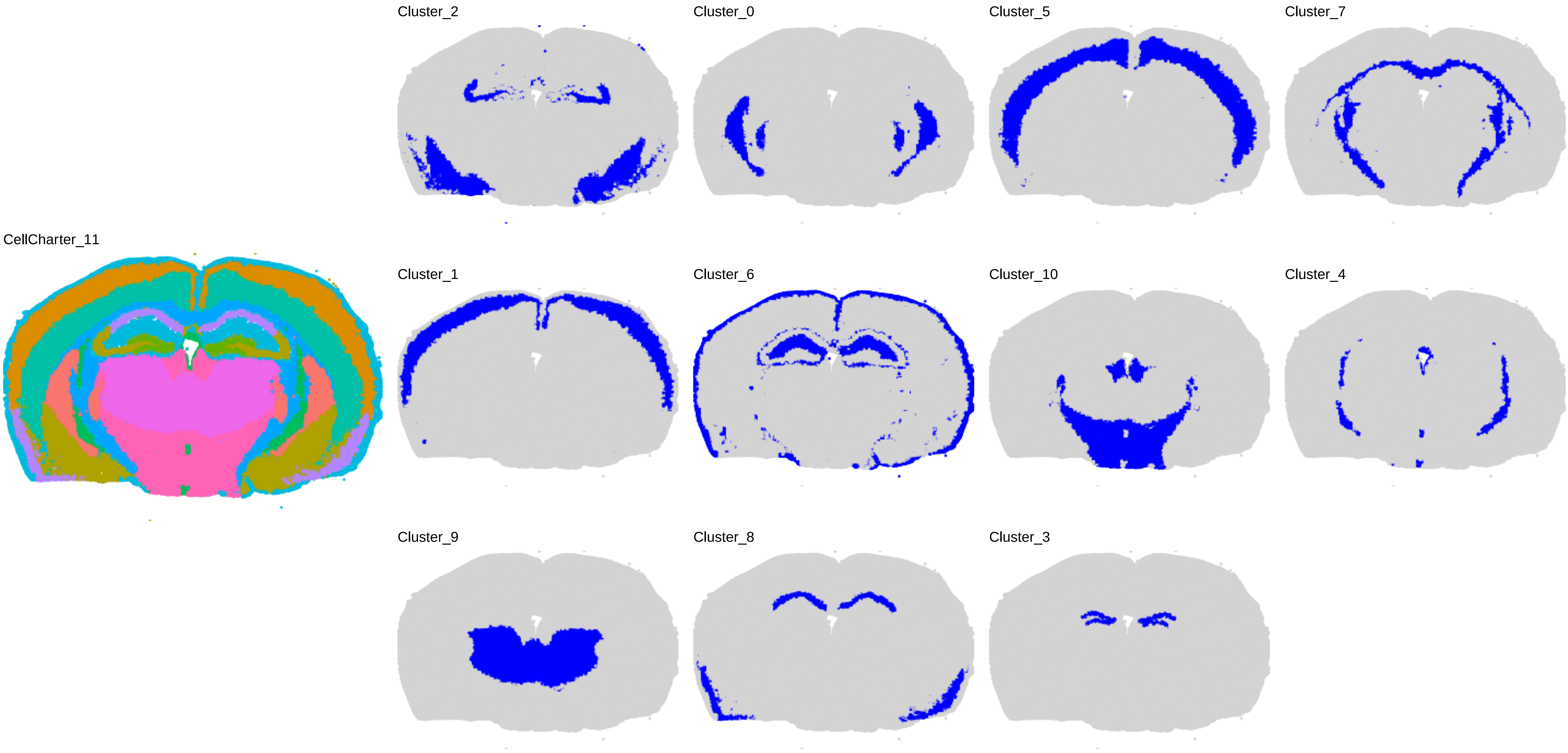

CellCharter_11
Cluster_2
Cluster_0
Cluster_5
Cluster_7
Cluster_1
Cluster_6
Cluster_10
Cluster_4
Cluster_9
Cluster_8
Cluster_3

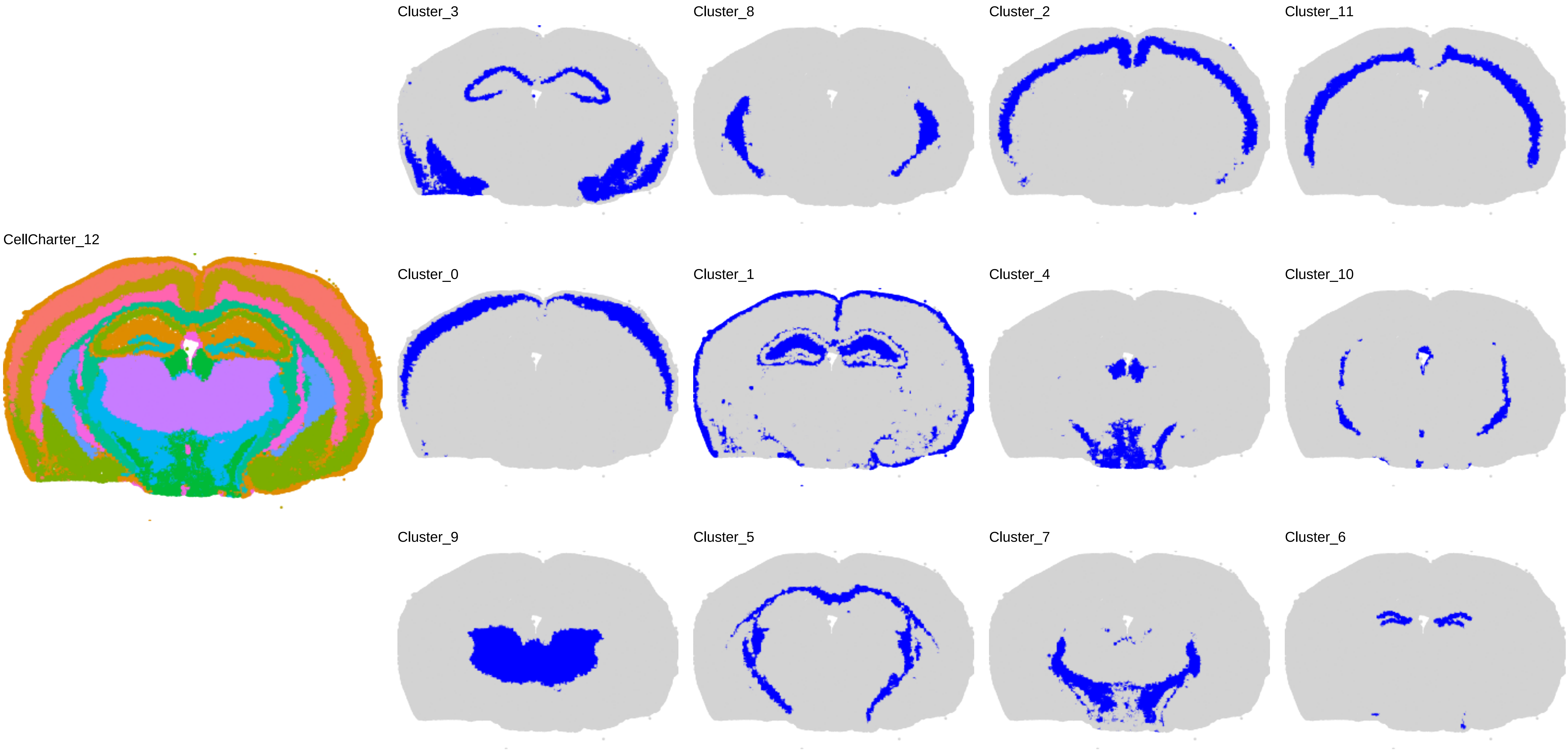
CellCharter_12
Cluster_3
Cluster_8
Cluster_2
Cluster_11
Cluster_0
Cluster_1
Cluster_4
Cluster_10
Cluster_9
Cluster_5
Cluster_7
Cluster_6

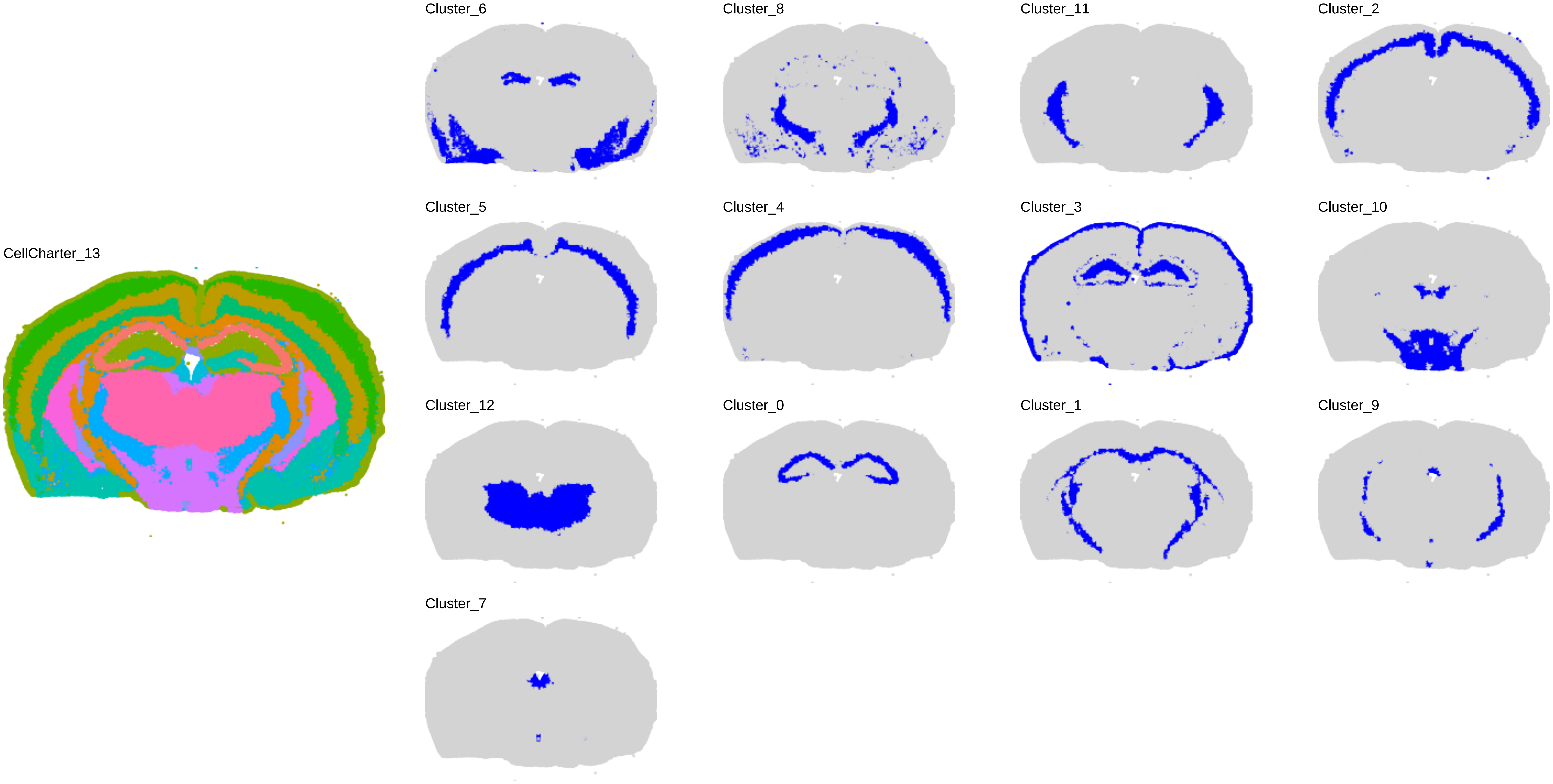
CellCharter_13
Cluster_6
Cluster_8
Cluster_11
Cluster_2
Cluster_5
Cluster_4
Cluster_3
Cluster_10
Cluster_12
Cluster_0
Cluster_1
Cluster_9
Cluster_7

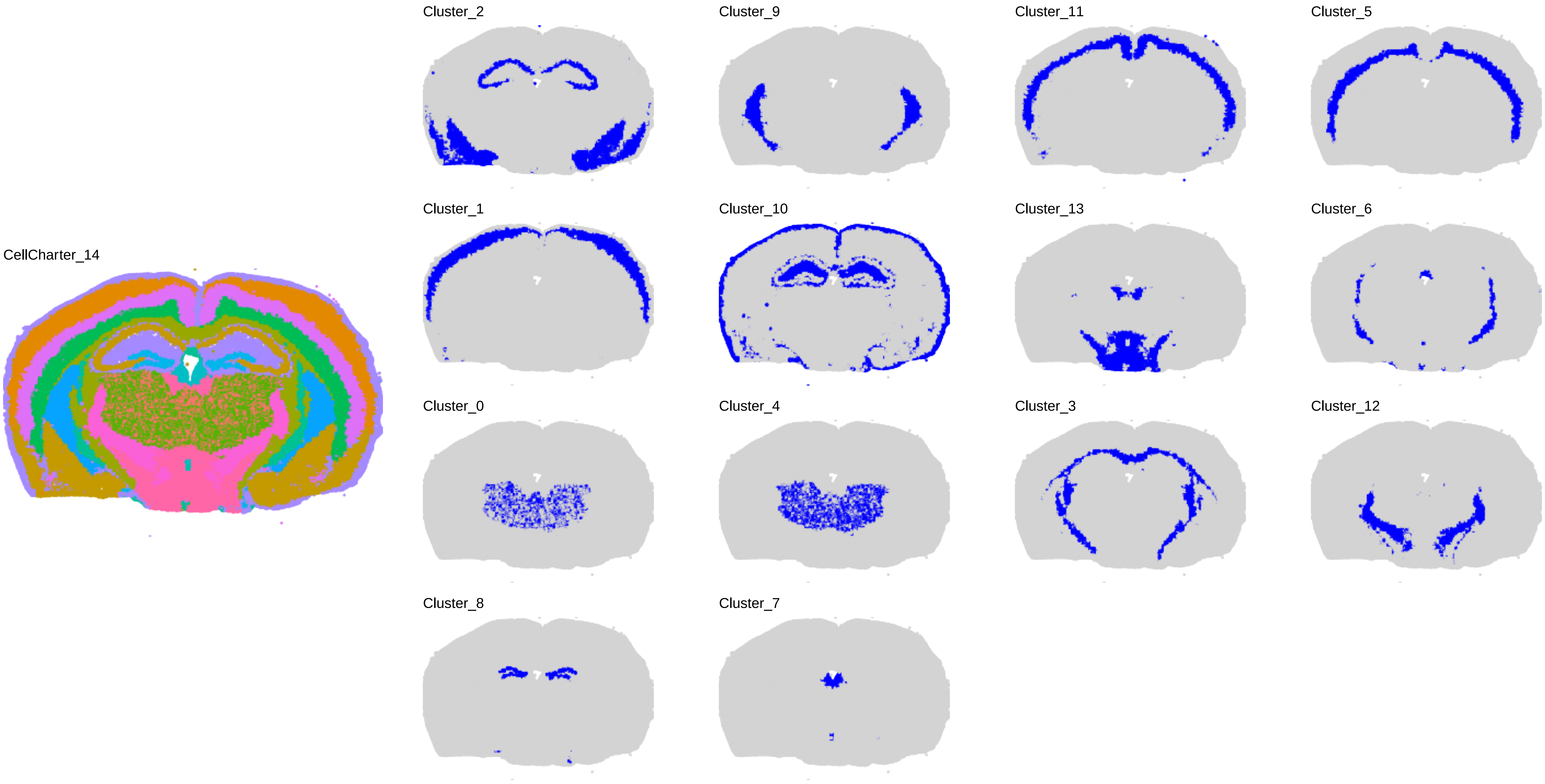
CellCharter_14
Cluster_2
Cluster_9
Cluster_11
Cluster_5
Cluster_1
Cluster_10
Cluster_13
Cluster_6
Cluster_0
Cluster_4
Cluster_3
Cluster_12
Cluster_8
Cluster_7

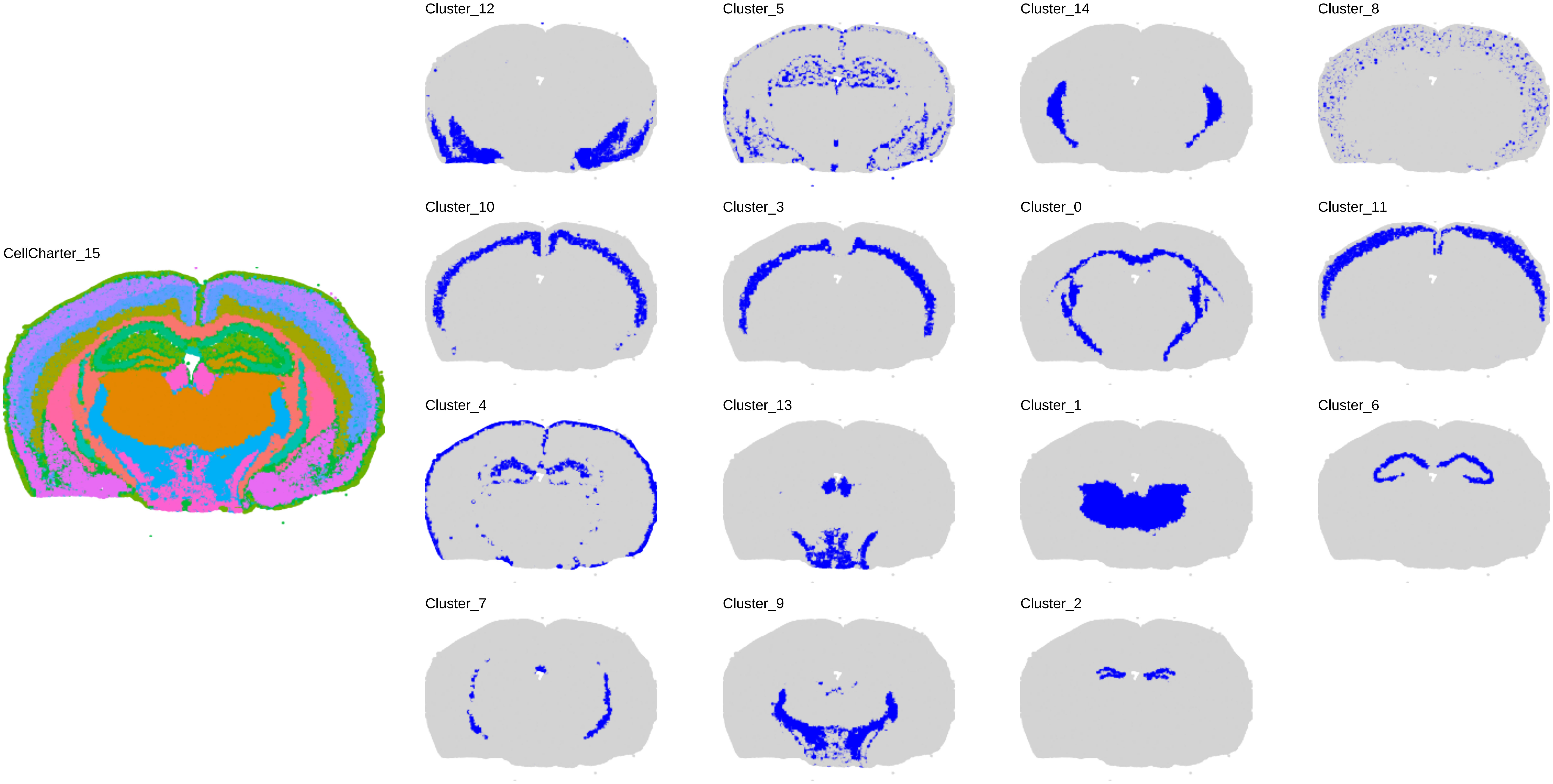
CellCharter_15
Cluster_12
Cluster_5
Cluster_14
Cluster_8
Cluster_10
Cluster_3
Cluster_0
Cluster_11
Cluster_4
Cluster_13
Cluster_1
Cluster_6
Cluster_7
Cluster_9
Cluster_2

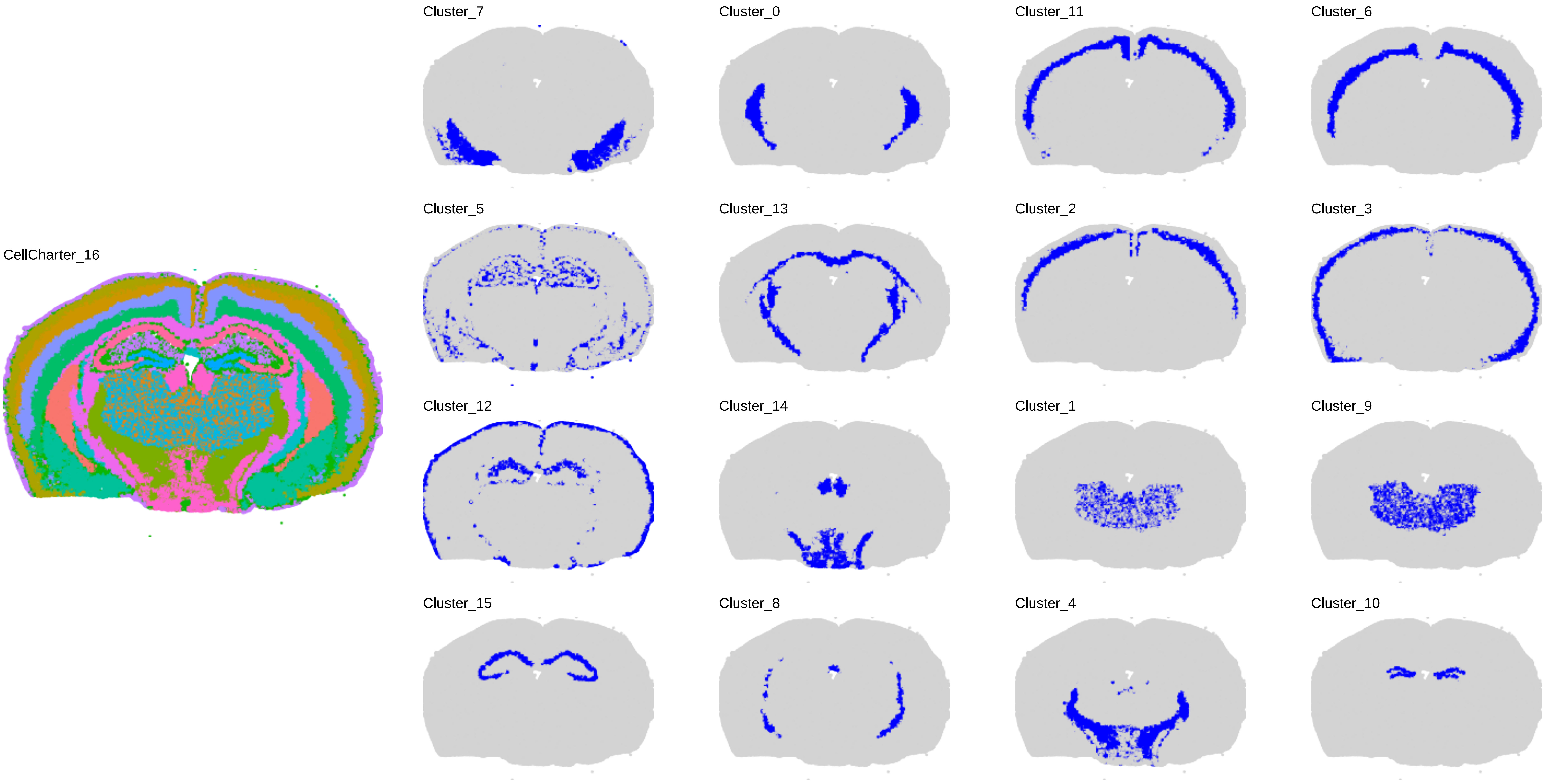
CellCharter_16
Cluster_7
Cluster_0
Cluster_11
Cluster_6
Cluster_5
Cluster_13
Cluster_2
Cluster_3
Cluster_12
Cluster_14
Cluster_1
Cluster_9
Cluster_15
Cluster_8
Cluster_4
Cluster_10

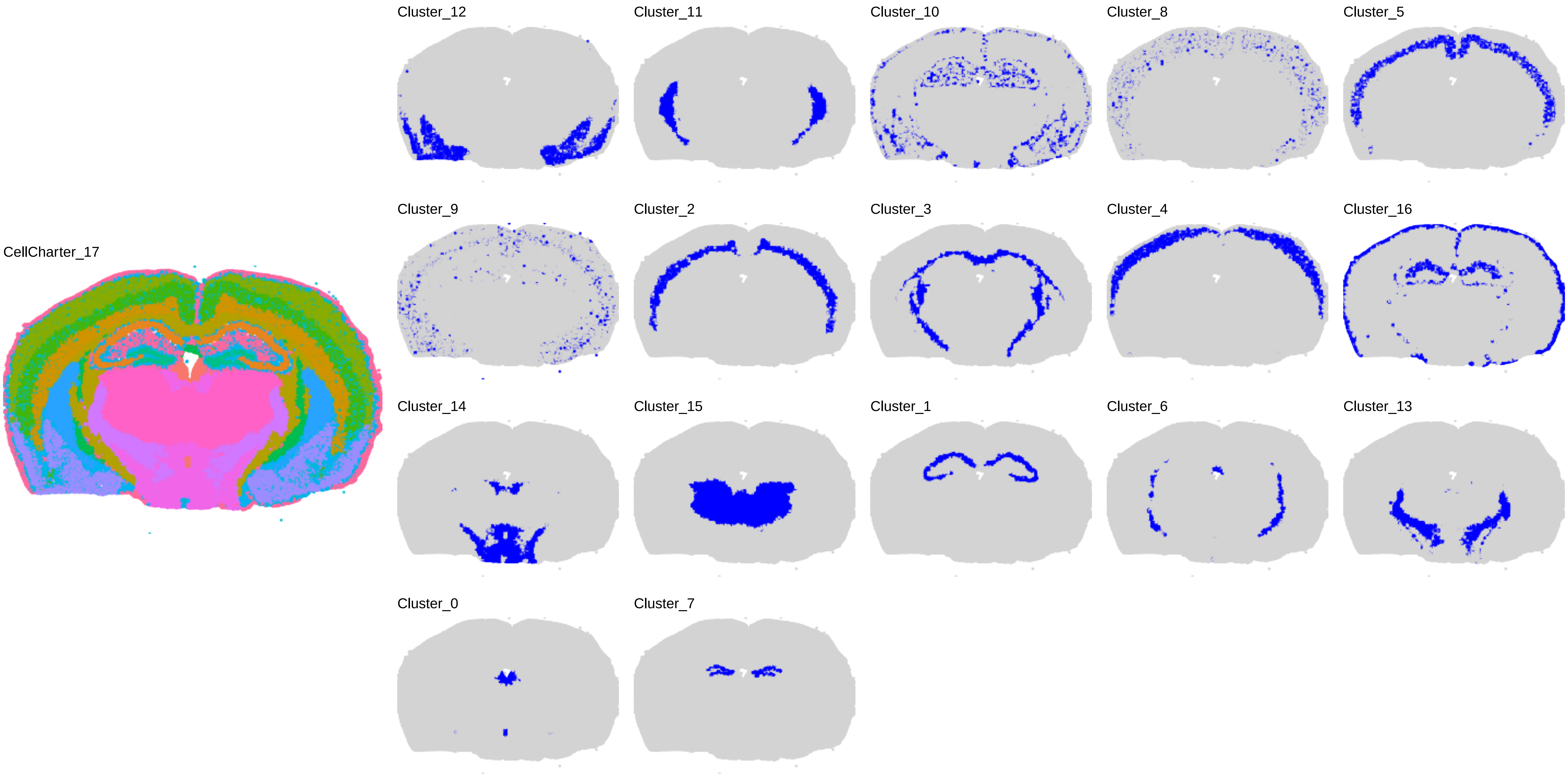
CellCharter_17
Cluster_12
Cluster_11
Cluster_10
Cluster_8
Cluster_5
Cluster_9
Cluster_2
Cluster_3
Cluster_4
Cluster_16
Cluster_14
Cluster_15
Cluster_1
Cluster_6
Cluster_13
Cluster_0
Cluster_7

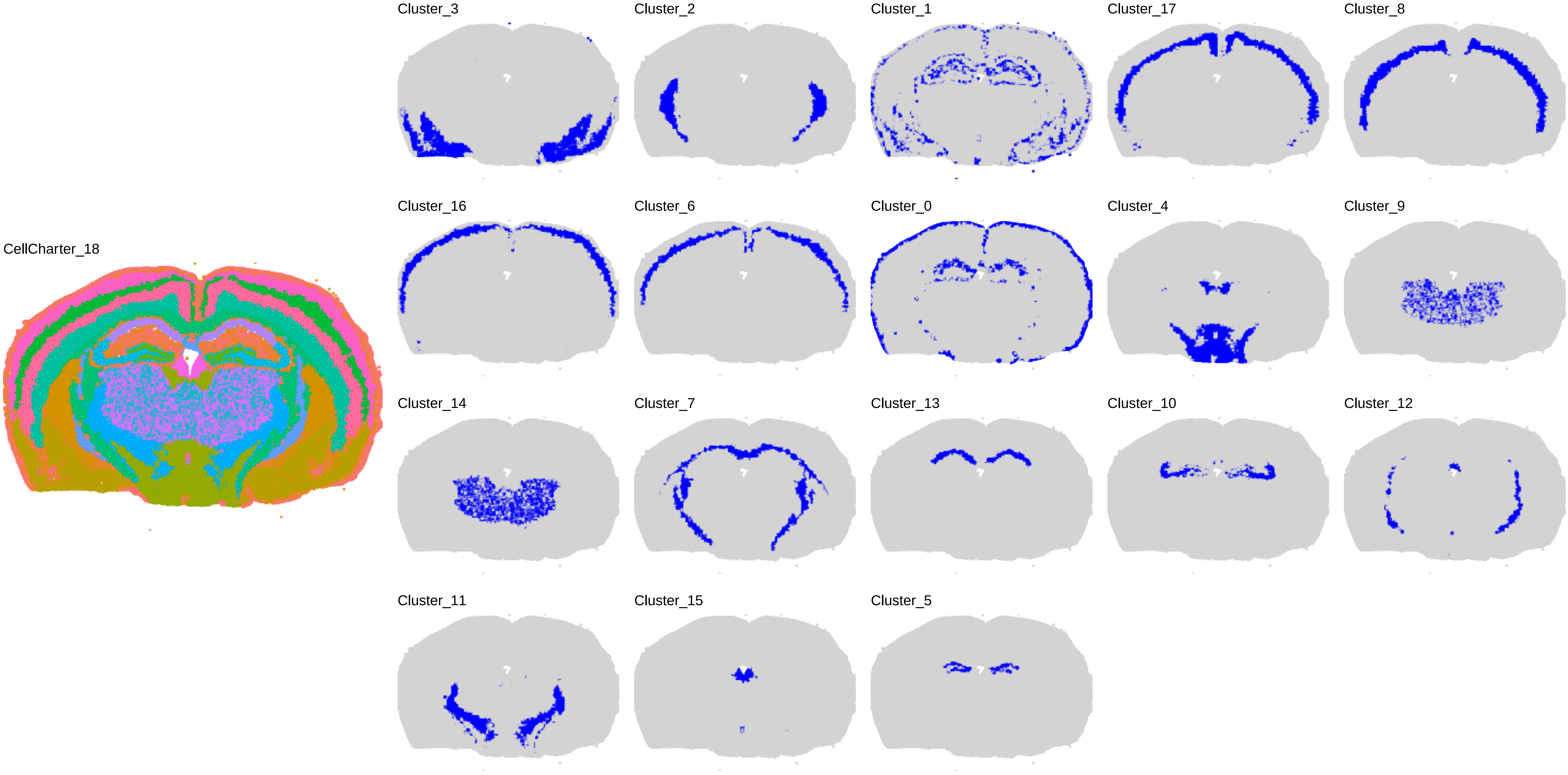

CellCharter_18
Cluster_3
Cluster_2
Cluster_1
Cluster_17
Cluster_8
Cluster_16
Cluster_6
Cluster_0
Cluster_4
Cluster_9
Cluster_14
Cluster_7
Cluster_13
Cluster_10
Cluster_12
Cluster_11
Cluster_15
Cluster_5

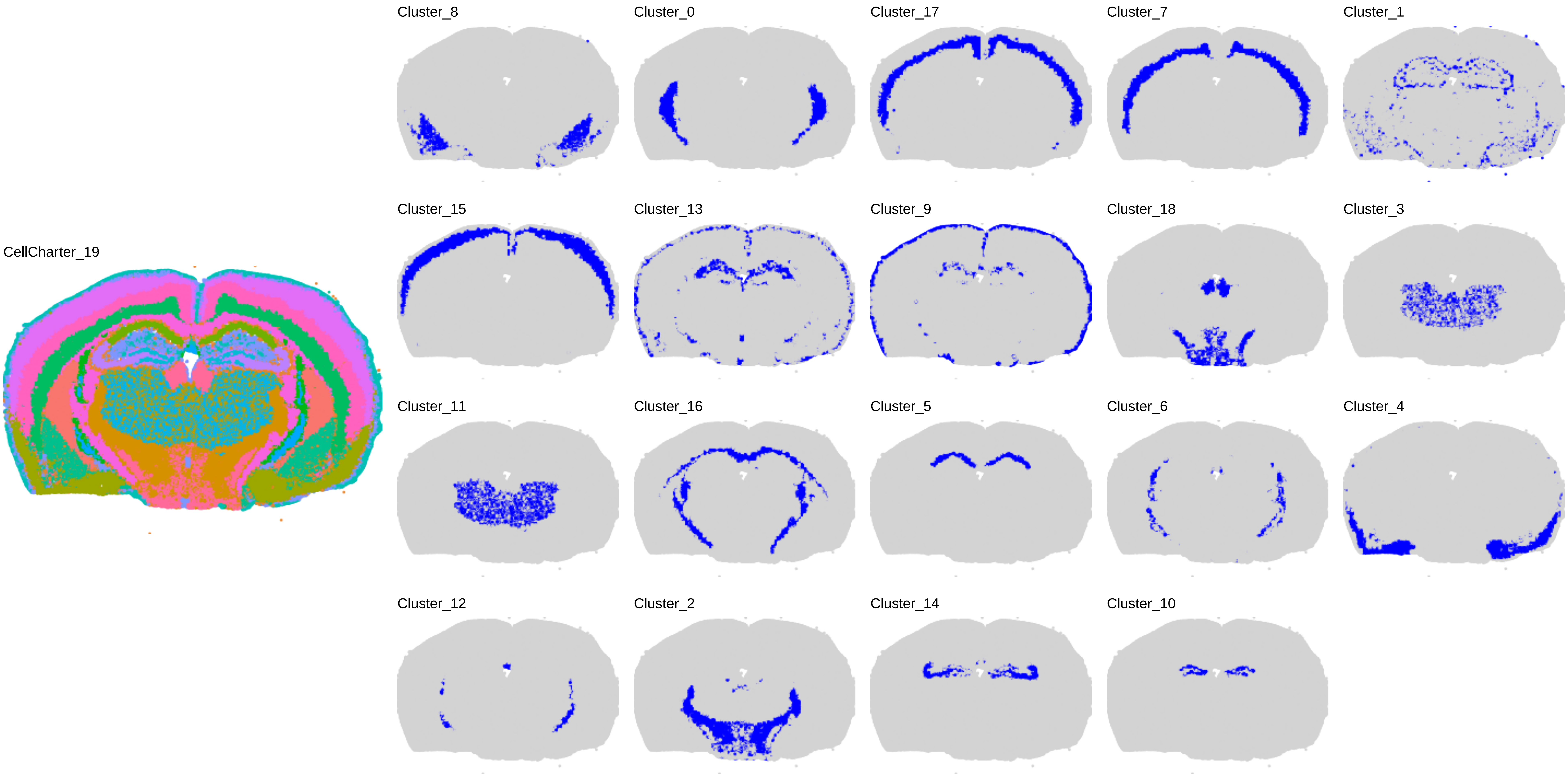
CellCharter_19
Cluster_8
Cluster_0
Cluster_17
Cluster_7
Cluster_1
Cluster_15
Cluster_13
Cluster_9
Cluster_18
Cluster_3
Cluster_11
Cluster_16
Cluster_5
Cluster_6
Cluster_4
Cluster_12
Cluster_2
Cluster_14
Cluster_10

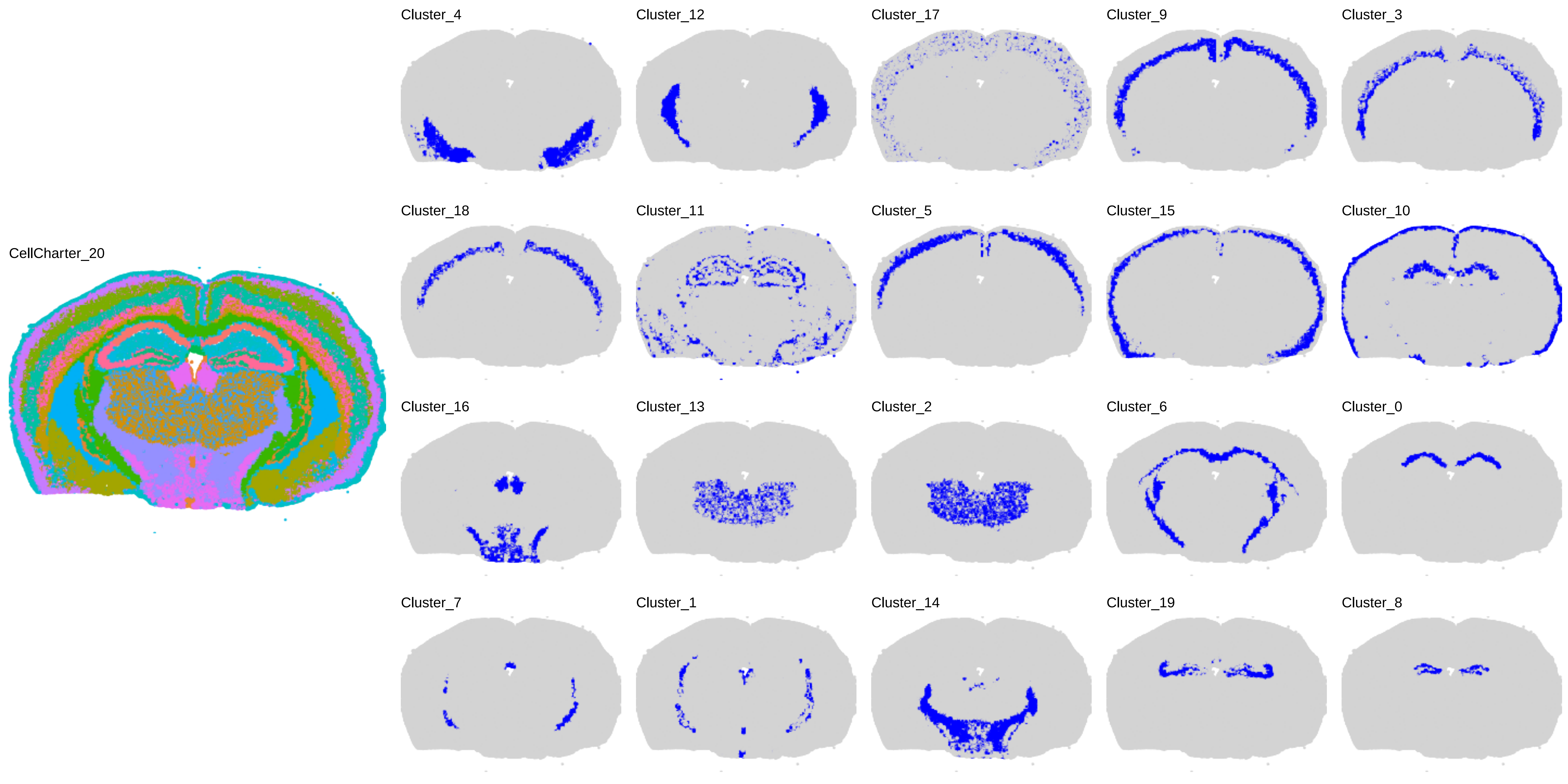
CellCharter_20
Cluster_4
Cluster_12
Cluster_17
Cluster_9
Cluster_3
Cluster_18
Cluster_11
Cluster_5
Cluster_15
Cluster_10
Cluster_16
Cluster_13
Cluster_2
Cluster_6
Cluster_0
Cluster_7
Cluster_1
Cluster_14
Cluster_19
Cluster_8

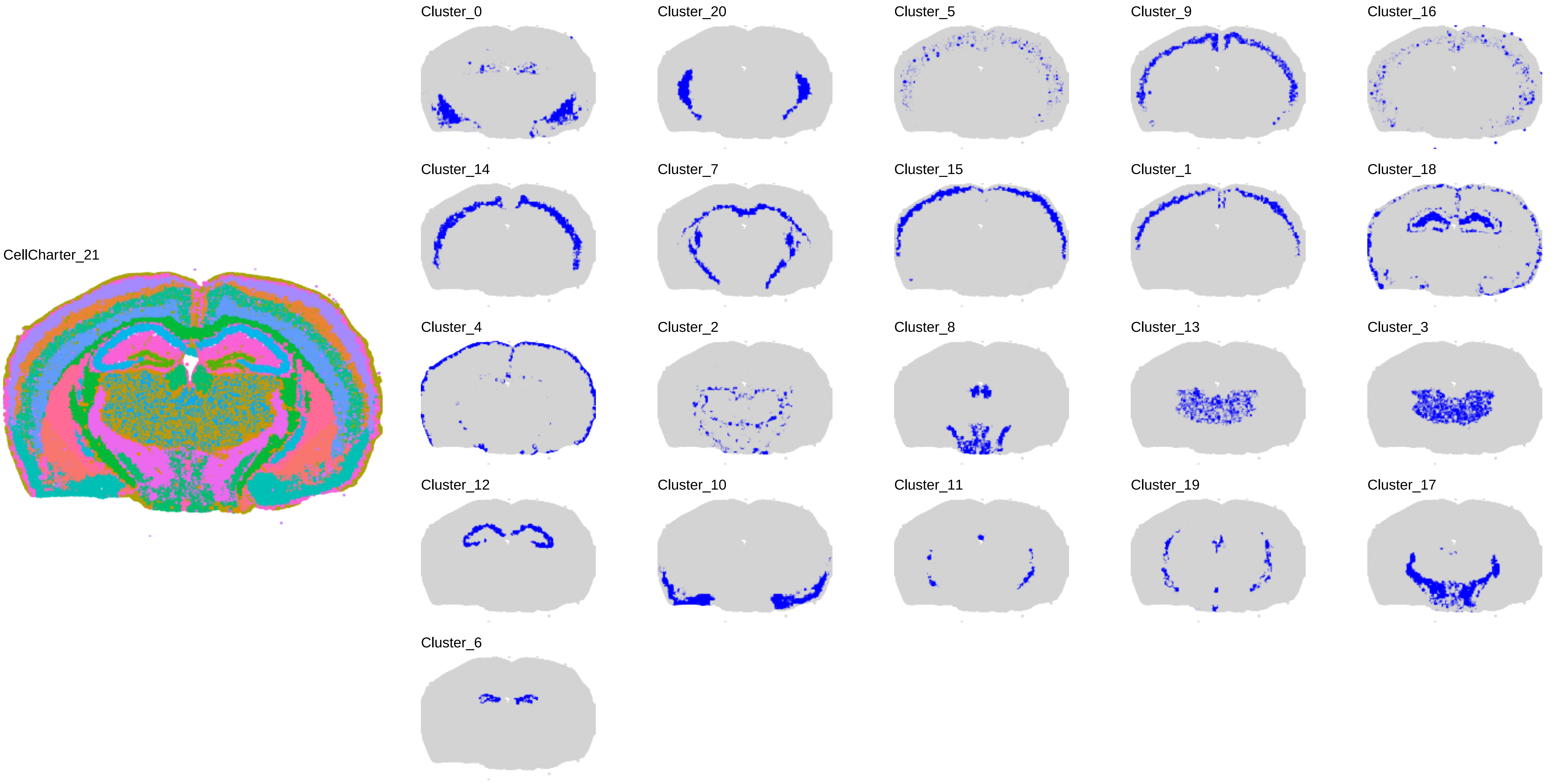
CellCharter_21
Cluster_0
Cluster_20
Cluster_5
Cluster_9
Cluster_16
Cluster_14
Cluster_7
Cluster_15
Cluster_1
Cluster_18
Cluster_4
Cluster_2
Cluster_8
Cluster_13
Cluster_3
Cluster_12
Cluster_10
Cluster_11
Cluster_19
Cluster_17
Cluster_6

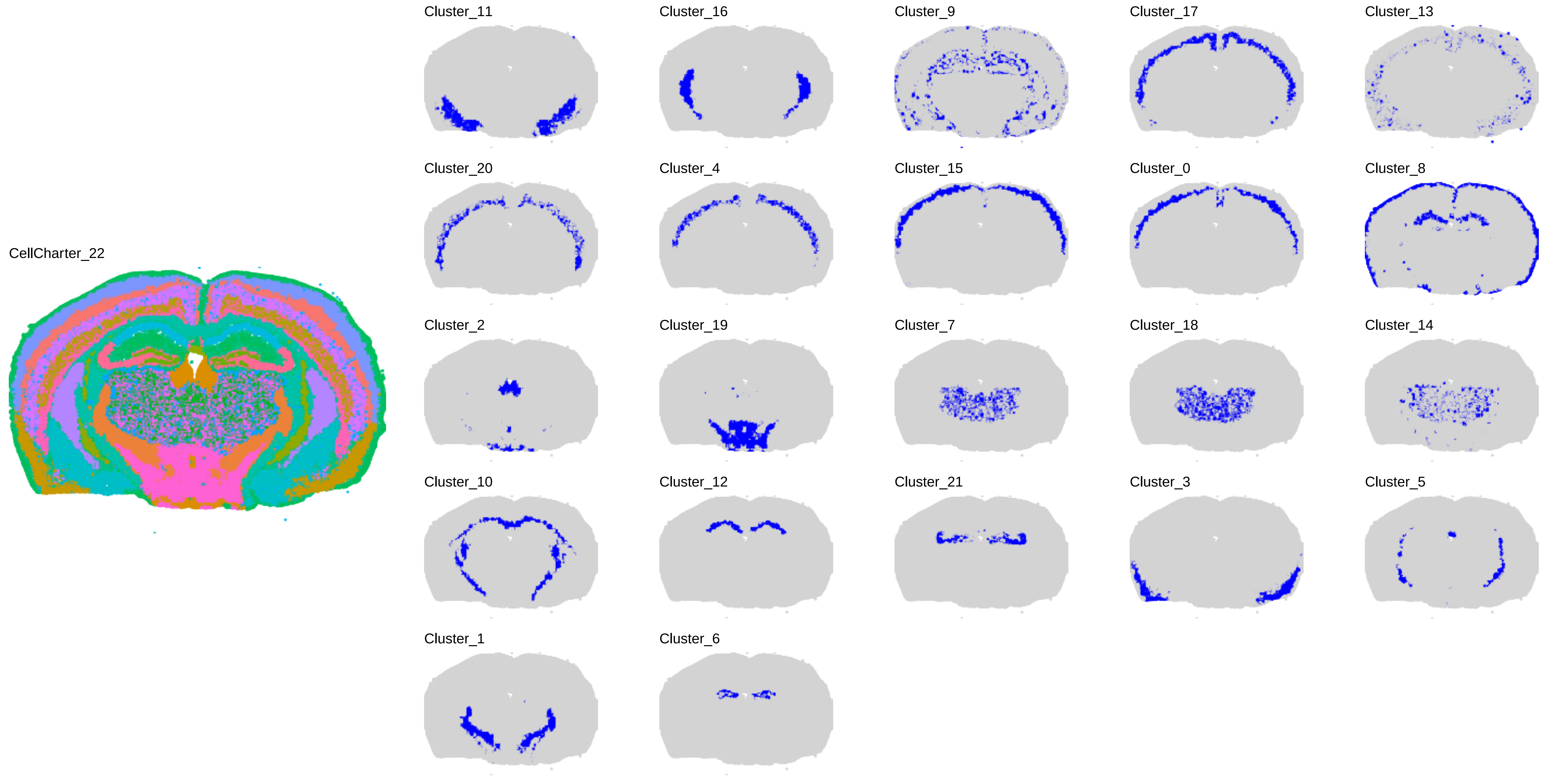
CellCharter_22
Cluster_11
Cluster_16
Cluster_9
Cluster_17
Cluster_13
Cluster_20
Cluster_4
Cluster_15
Cluster_0
Cluster_8
Cluster_2
Cluster_19
Cluster_7
Cluster_18
Cluster_14
Cluster_10
Cluster_12
Cluster_21
Cluster_3
Cluster_5
Cluster_1
Cluster_6

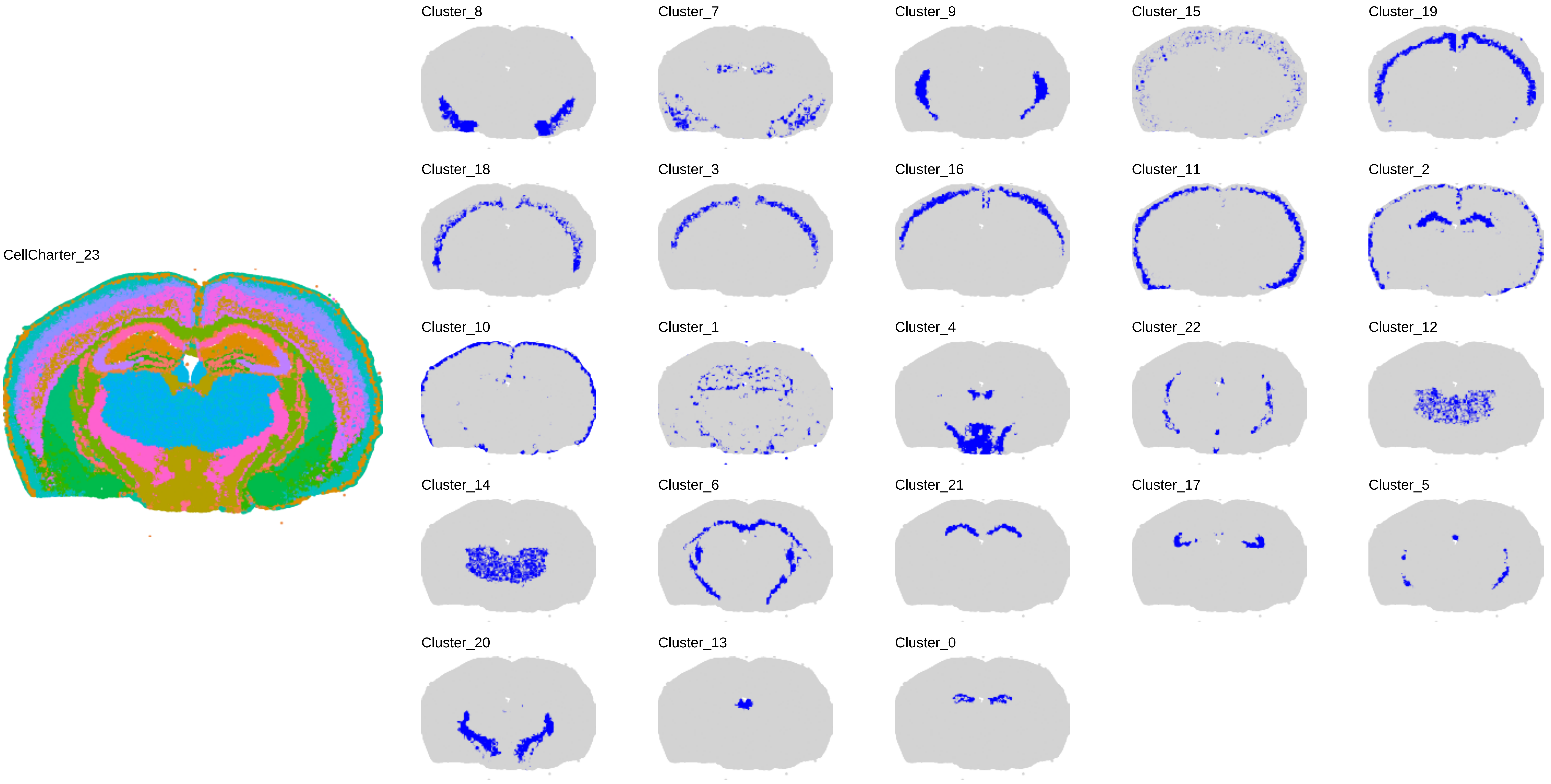
CellCharter_23
Cluster_8
Cluster_7
Cluster_9
Cluster_15
Cluster_19
Cluster_18
Cluster_3
Cluster_16
Cluster_11
Cluster_2
Cluster_10
Cluster_1
Cluster_4
Cluster_22
Cluster_12
Cluster_14
Cluster_6
Cluster_21
Cluster_17
Cluster_5
Cluster_20
Cluster_13
Cluster_0

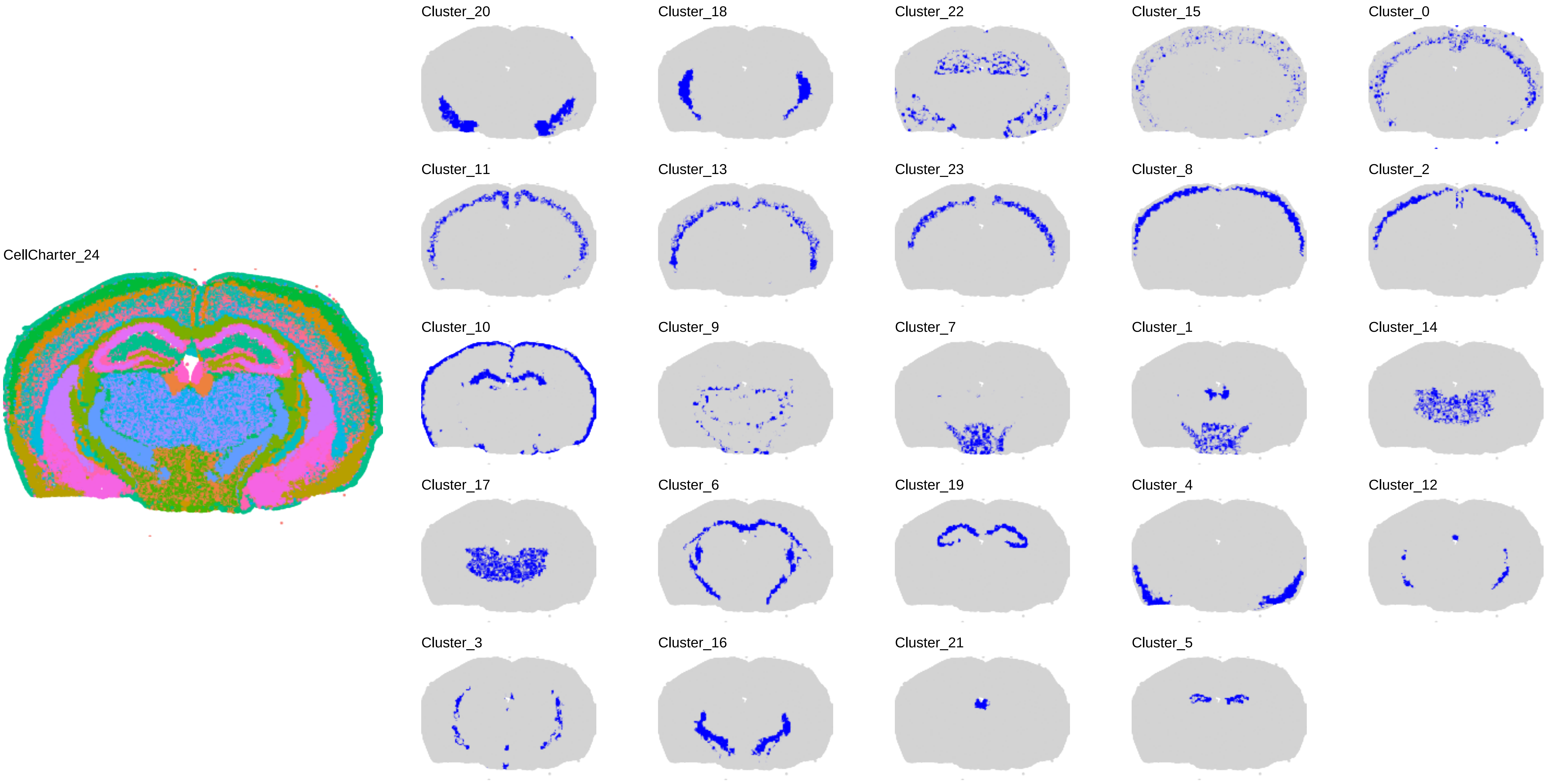
CellCharter_24
Cluster_20
Cluster_18
Cluster_22
Cluster_15
Cluster_0
Cluster_11
Cluster_13
Cluster_23
Cluster_8
Cluster_2
Cluster_10
Cluster_9
Cluster_7
Cluster_1
Cluster_14
Cluster_17
Cluster_6
Cluster_19
Cluster_4
Cluster_12
Cluster_3
Cluster_16
Cluster_21
Cluster_5

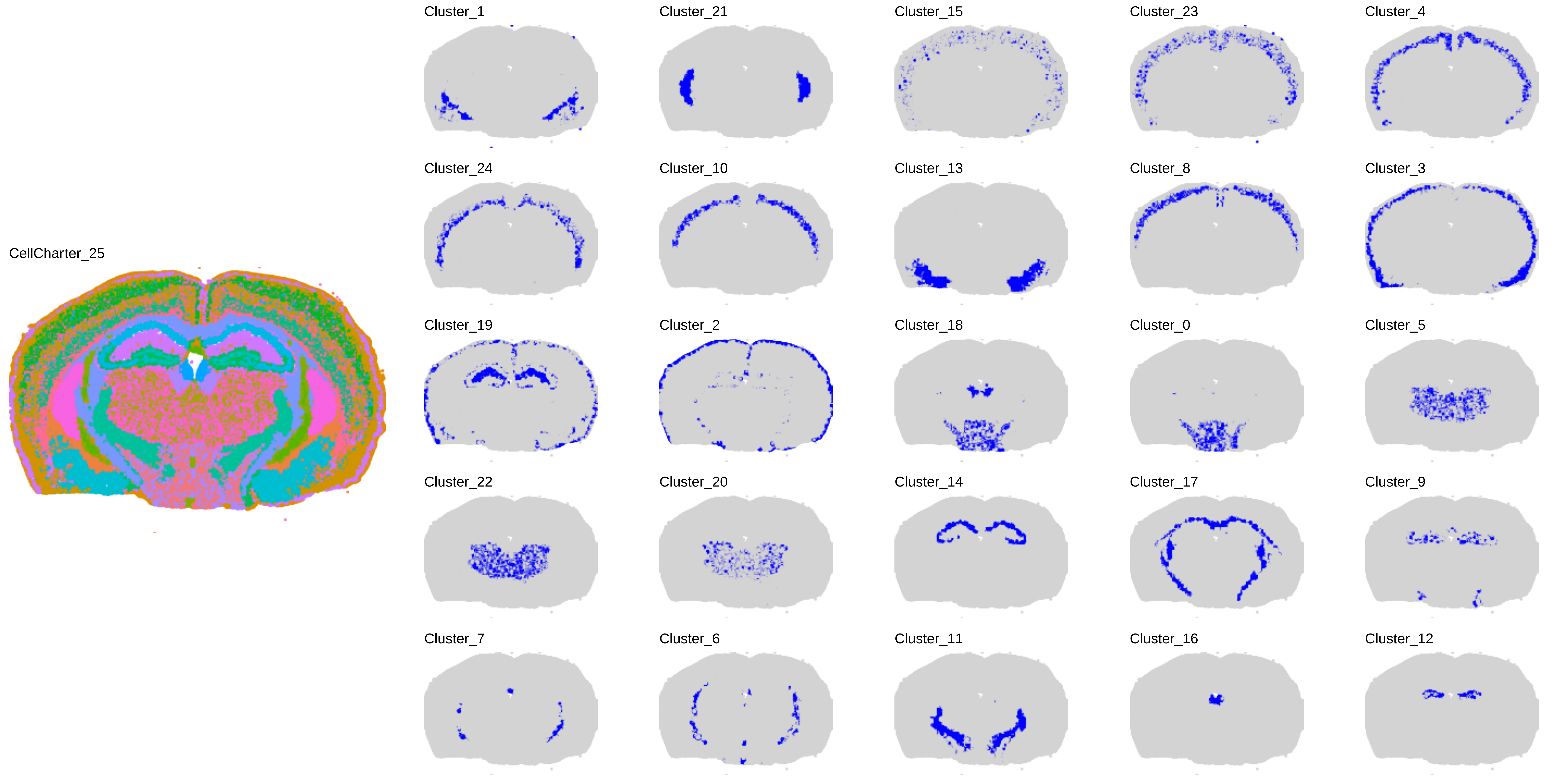
CellCharter_25
Cluster_1
Cluster_21
Cluster_15
Cluster_23
Cluster_4
Cluster_24
Cluster_10
Cluster_13
Cluster_8
Cluster_3
Cluster_19
Cluster_2
Cluster_18
Cluster_0
Cluster_5
Cluster_22
Cluster_20
Cluster_14
Cluster_17
Cluster_9
Cluster_7
Cluster_6
Cluster_11
Cluster_16
Cluster_12

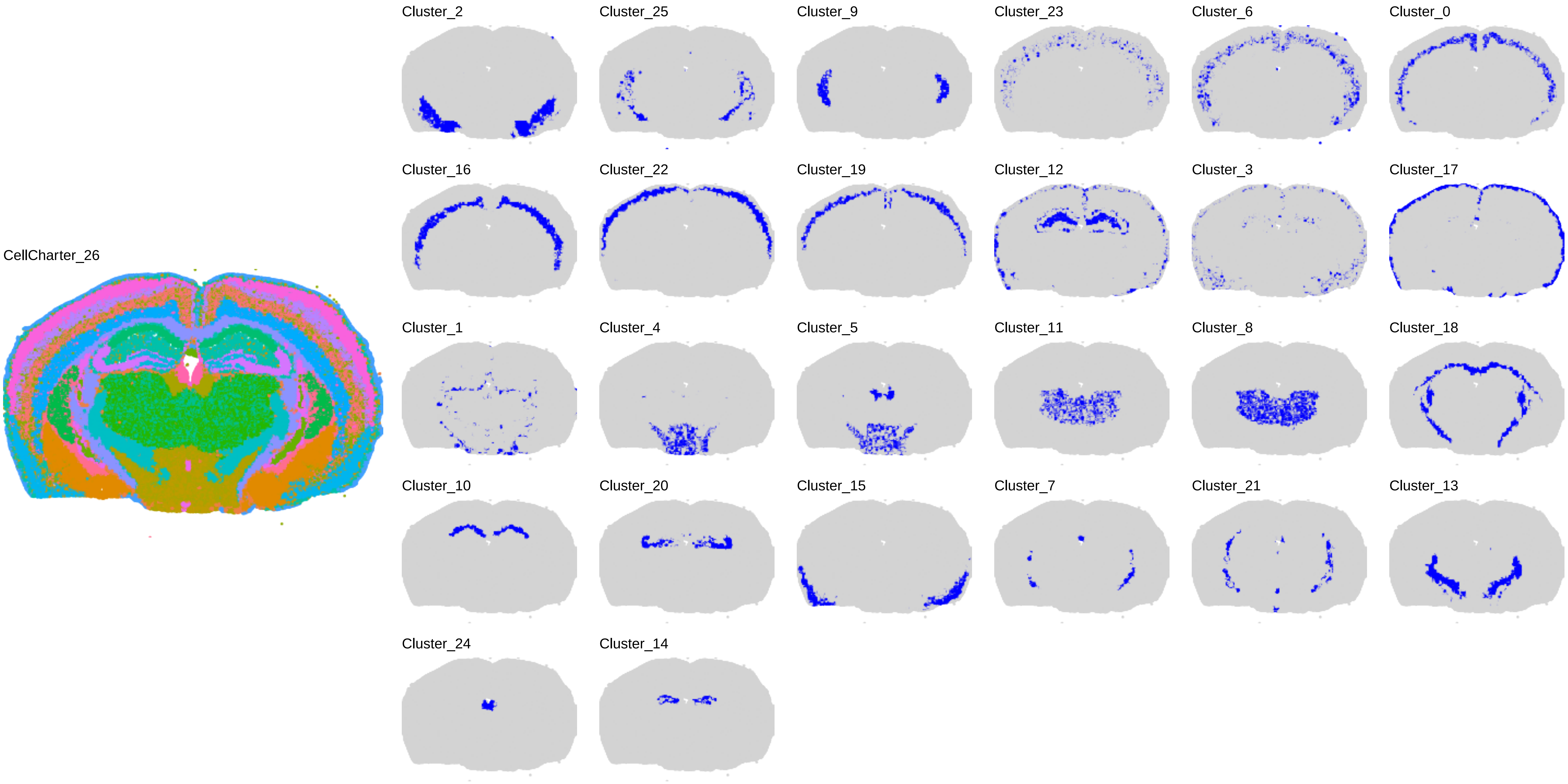
CellCharter_26
Cluster_2
Cluster_25
Cluster_9
Cluster_23
Cluster_6
Cluster_0
Cluster_16
Cluster_22
Cluster_19
Cluster_12
Cluster_3
Cluster_17
Cluster_1
Cluster_4
Cluster_5
Cluster_11
Cluster_8
Cluster_18
Cluster_10
Cluster_20
Cluster_15
Cluster_7
Cluster_21
Cluster_13
Cluster_24
Cluster_14

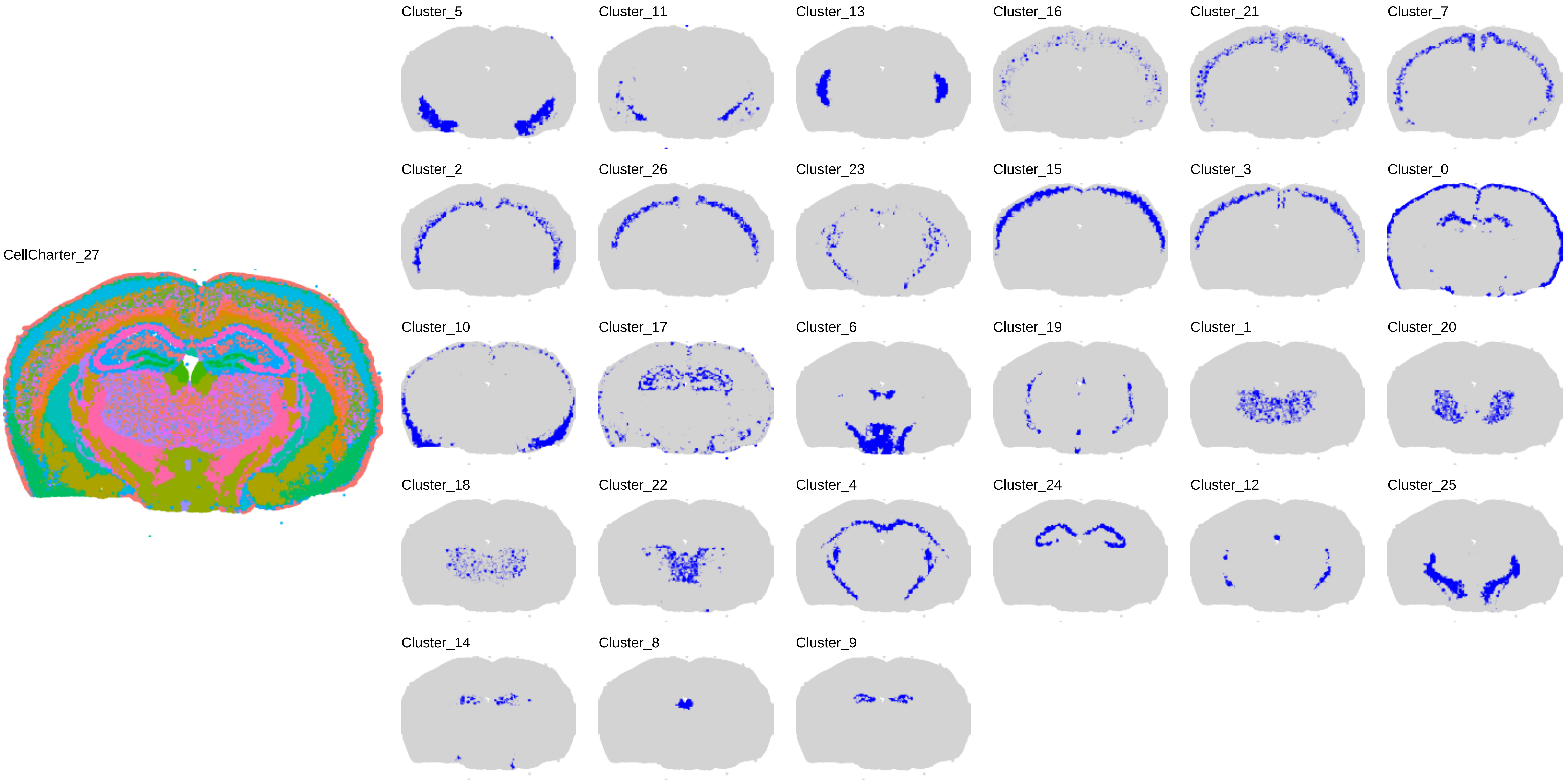

CellCharter_27
Cluster_5
Cluster_11
Cluster_13
Cluster_16
Cluster_21
Cluster_7
Cluster_2
Cluster_26
Cluster_23
Cluster_15
Cluster_3
Cluster_0
Cluster_10
Cluster_17
Cluster_6
Cluster_19
Cluster_1
Cluster_20
Cluster_18
Cluster_22
Cluster_4
Cluster_24
Cluster_12
Cluster_25
Cluster_14
Cluster_8
Cluster_9

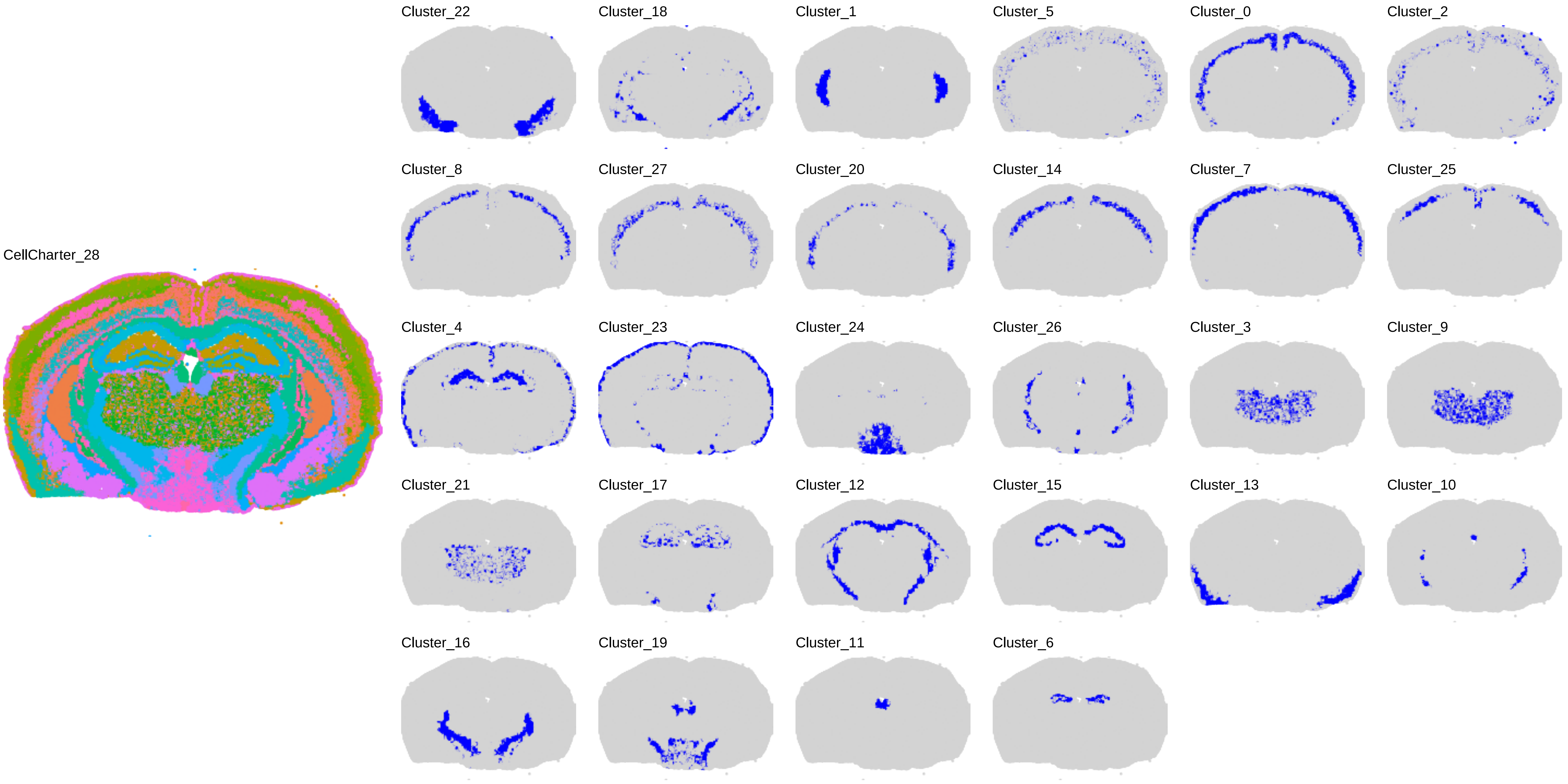
CellCharter_28
Cluster_22
Cluster_18
Cluster_1
Cluster_5
Cluster_0
Cluster_2
Cluster_8
Cluster_27
Cluster_20
Cluster_14
Cluster_7
Cluster_25
Cluster_4
Cluster_23
Cluster_24
Cluster_26
Cluster_3
Cluster_9
Cluster_21
Cluster_17
Cluster_12
Cluster_15
Cluster_13
Cluster_10
Cluster_16
Cluster_19
Cluster_11
Cluster_6

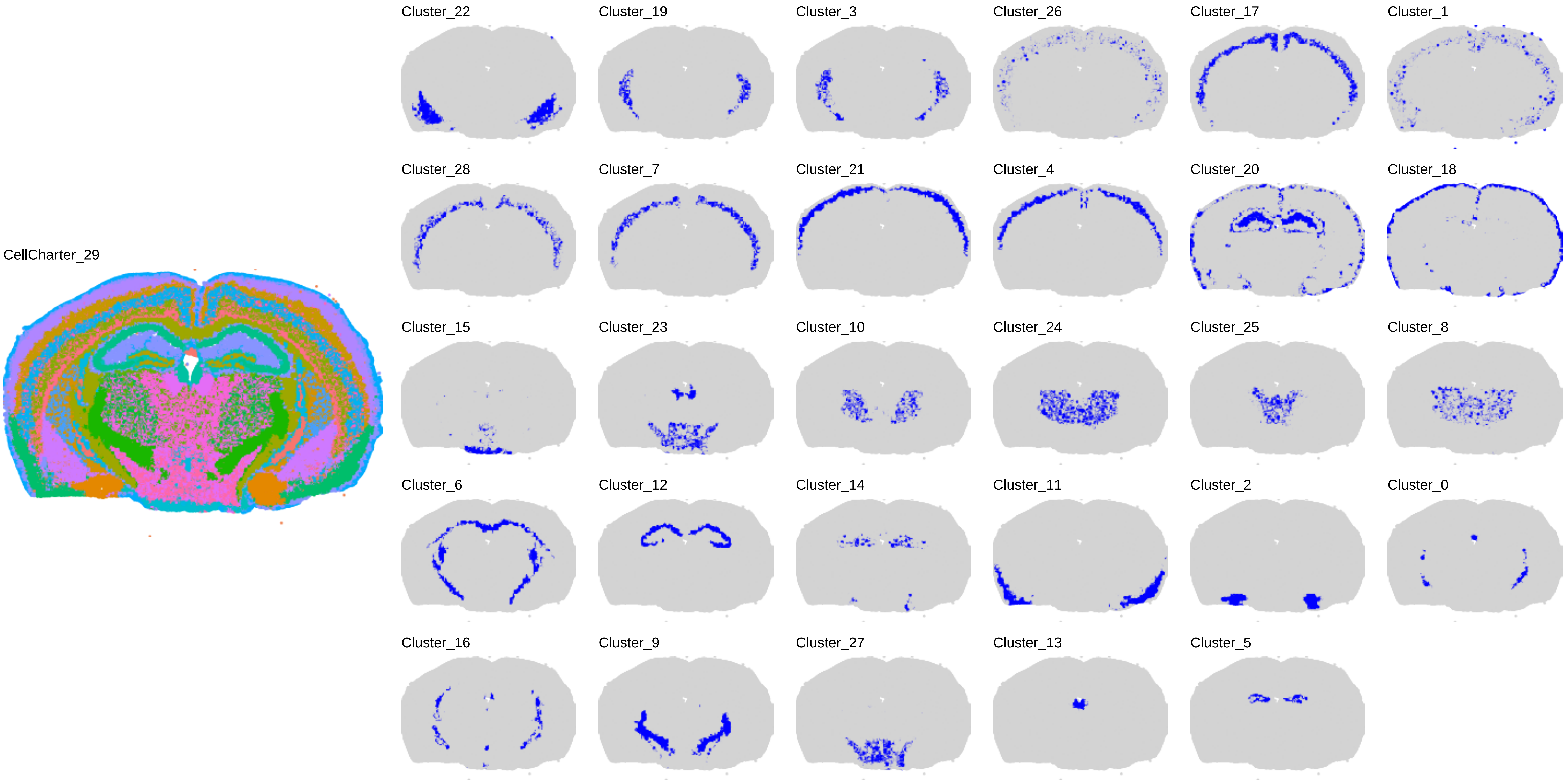
CellCharter_29
Cluster_22
Cluster_19
Cluster_3
Cluster_26
Cluster_17
Cluster_1
Cluster_28
Cluster_7
Cluster_21
Cluster_4
Cluster_20
Cluster_18
Cluster_15
Cluster_23
Cluster_10
Cluster_24
Cluster_25
Cluster_8
Cluster_6
Cluster_12
Cluster_14
Cluster_11
Cluster_2
Cluster_0
Cluster_16
Cluster_9
Cluster_27
Cluster_13
Cluster_5

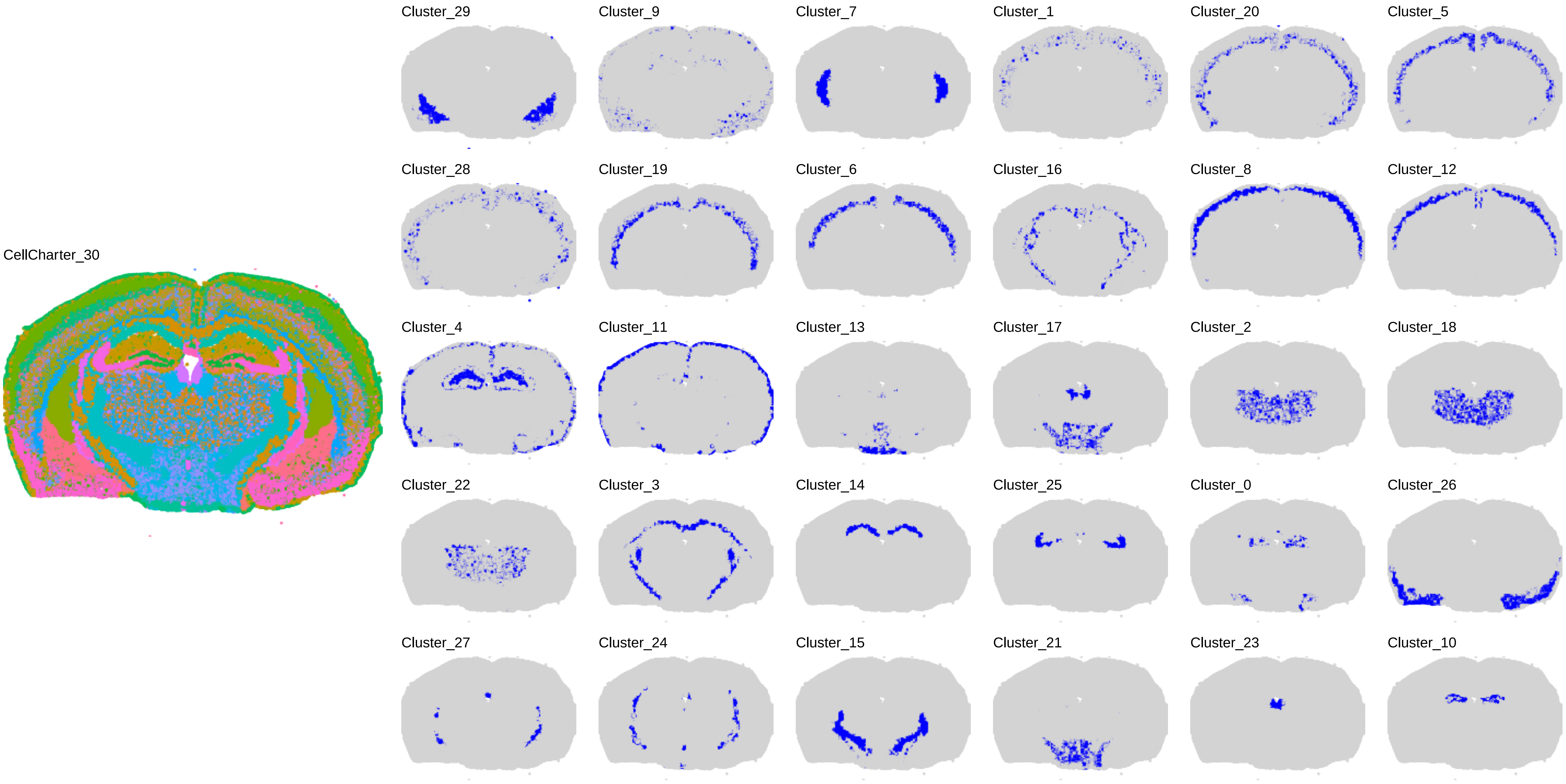

CellCharter_30
Cluster_29
Cluster_9
Cluster_7
Cluster_1
Cluster_20
Cluster_5
Cluster_28
Cluster_19
Cluster_6
Cluster_16
Cluster_8
Cluster_12
Cluster_4
Cluster_11
Cluster_13
Cluster_17
Cluster_2
Cluster_18
Cluster_22
Cluster_3
Cluster_14
Cluster_25
Cluster_0
Cluster_26
Cluster_27
Cluster_24
Cluster_15
Cluster_21
Cluster_23
Cluster_10

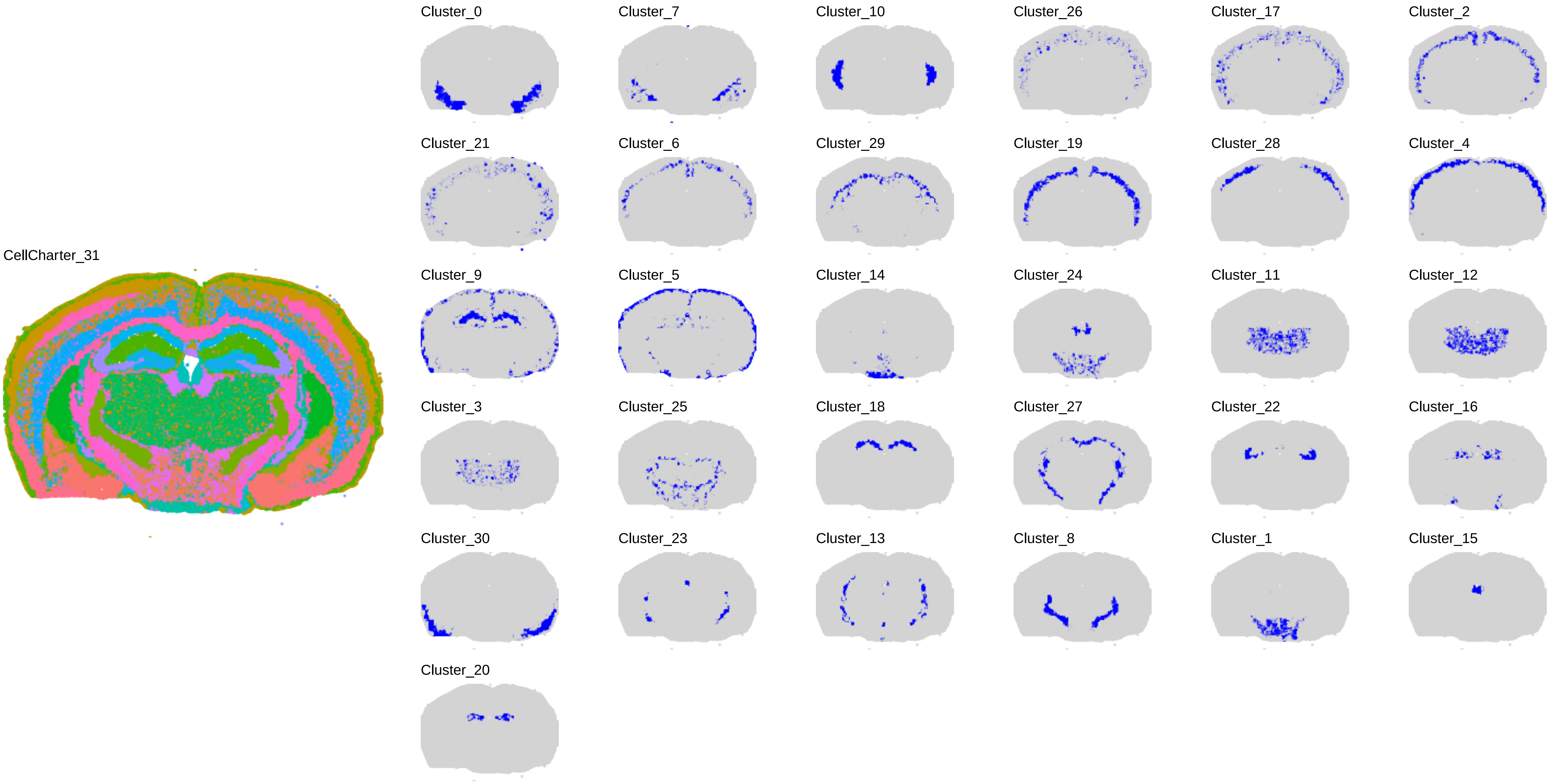
CellCharter_31
Cluster_0
Cluster_7
Cluster_10
Cluster_26
Cluster_17
Cluster_2
Cluster_21
Cluster_6
Cluster_29
Cluster_19
Cluster_28
Cluster_4
Cluster_9
Cluster_5
Cluster_14
Cluster_24
Cluster_11
Cluster_12
Cluster_3
Cluster_25
Cluster_18
Cluster_27
Cluster_22
Cluster_16
Cluster_30
Cluster_23
Cluster_13
Cluster_8
Cluster_1
Cluster_15
Cluster_20

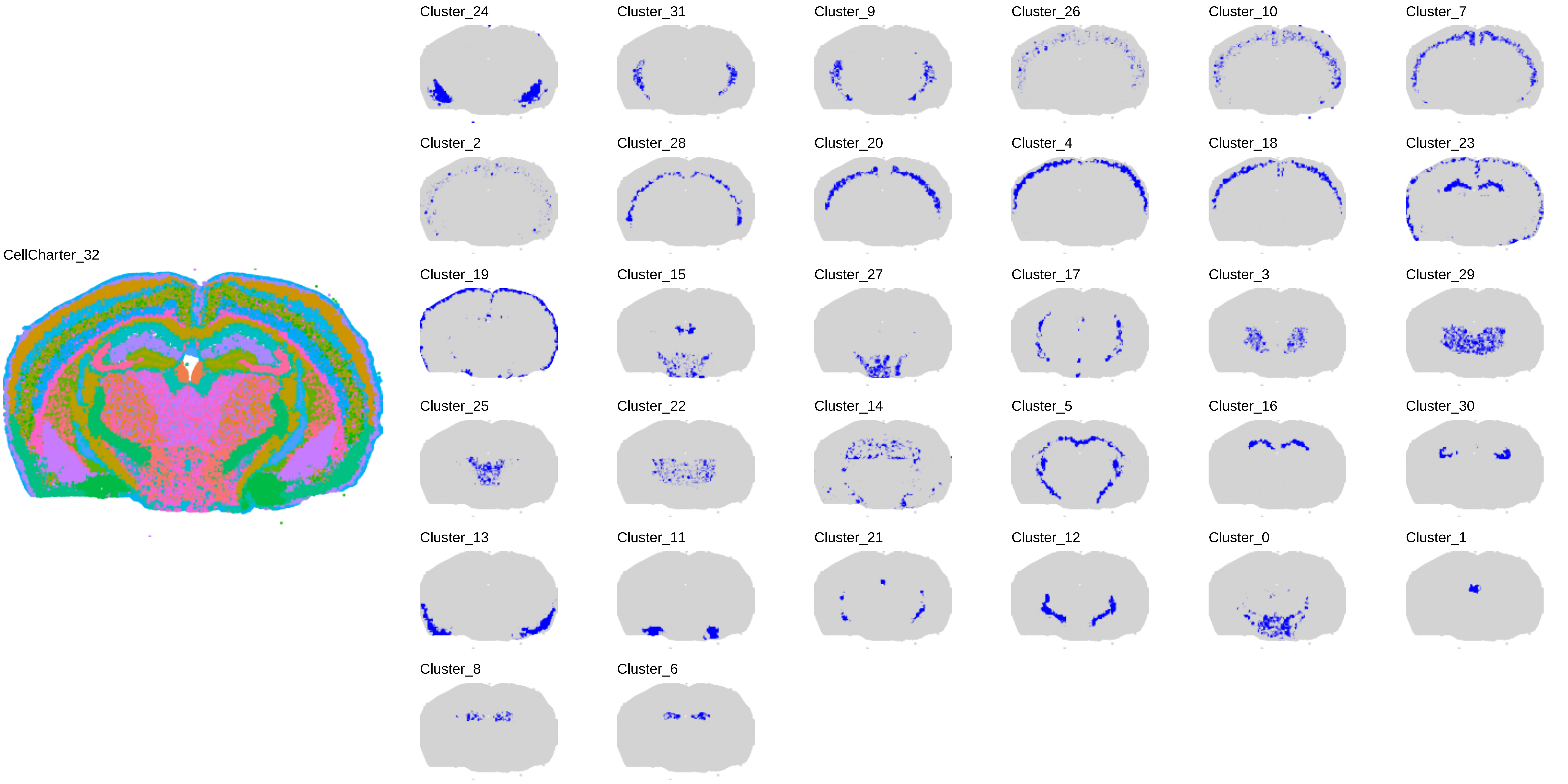
CellCharter_32
Cluster_24
Cluster_31
Cluster_9
Cluster_26
Cluster_10
Cluster_7
Cluster_2
Cluster_28
Cluster_20
Cluster_4
Cluster_18
Cluster_23
Cluster_19
Cluster_15
Cluster_27
Cluster_17
Cluster_3
Cluster_29
Cluster_25
Cluster_22
Cluster_14
Cluster_5
Cluster_16
Cluster_30
Cluster_13
Cluster_11
Cluster_21
Cluster_12
Cluster_0
Cluster_1
Cluster_8
Cluster_6

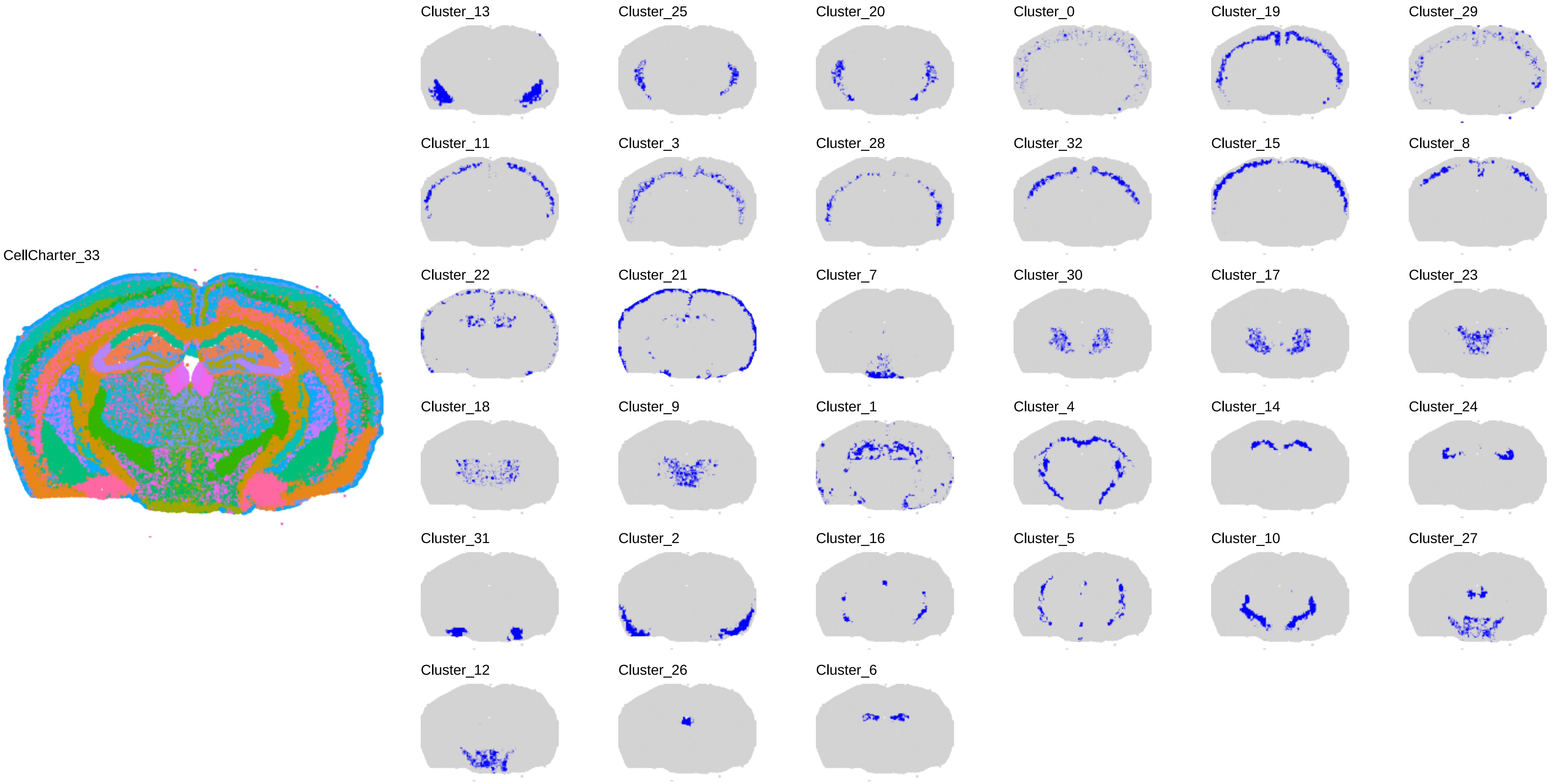
CellCharter_33
Cluster_13
Cluster_25
Cluster_20
Cluster_0
Cluster_19
Cluster_29
Cluster_11
Cluster_3
Cluster_28
Cluster_32
Cluster_15
Cluster_8
Cluster_22
Cluster_21
Cluster_7
Cluster_30
Cluster_17
Cluster_23
Cluster_18
Cluster_9
Cluster_1
Cluster_4
Cluster_14
Cluster_24
Cluster_31
Cluster_2
Cluster_16
Cluster_5
Cluster_10
Cluster_27
Cluster_12
Cluster_26
Cluster_6

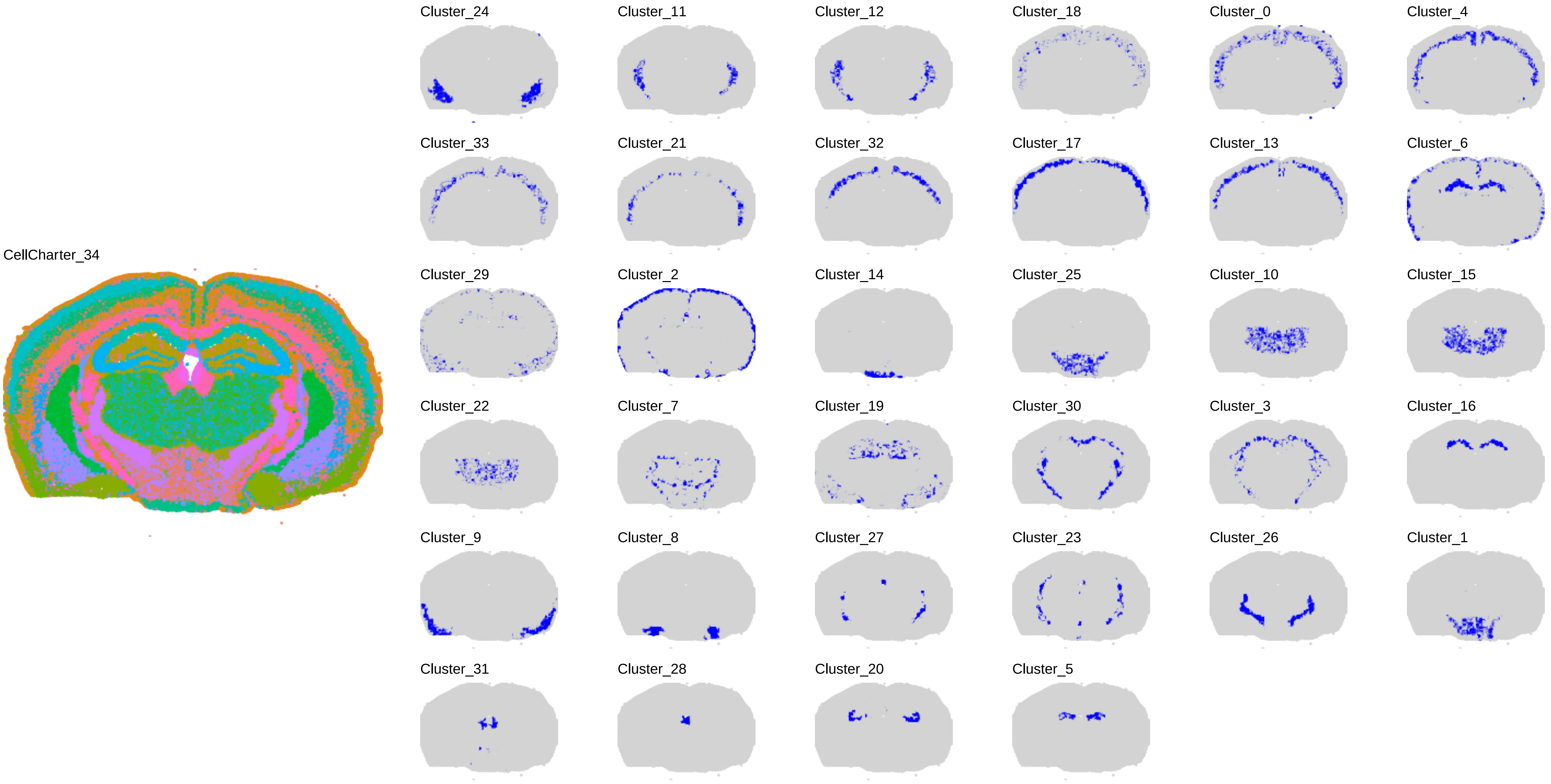

CellCharter_34
Cluster_24
Cluster_11
Cluster_12
Cluster_18
Cluster_0
Cluster_4
Cluster_33
Cluster_21
Cluster_32
Cluster_17
Cluster_13
Cluster_6
Cluster_29
Cluster_2
Cluster_14
Cluster_25
Cluster_10
Cluster_15
Cluster_22
Cluster_7
Cluster_19
Cluster_30
Cluster_3
Cluster_16
Cluster_9
Cluster_8
Cluster_27
Cluster_23
Cluster_26
Cluster_1
Cluster_31
Cluster_28
Cluster_20
Cluster_5

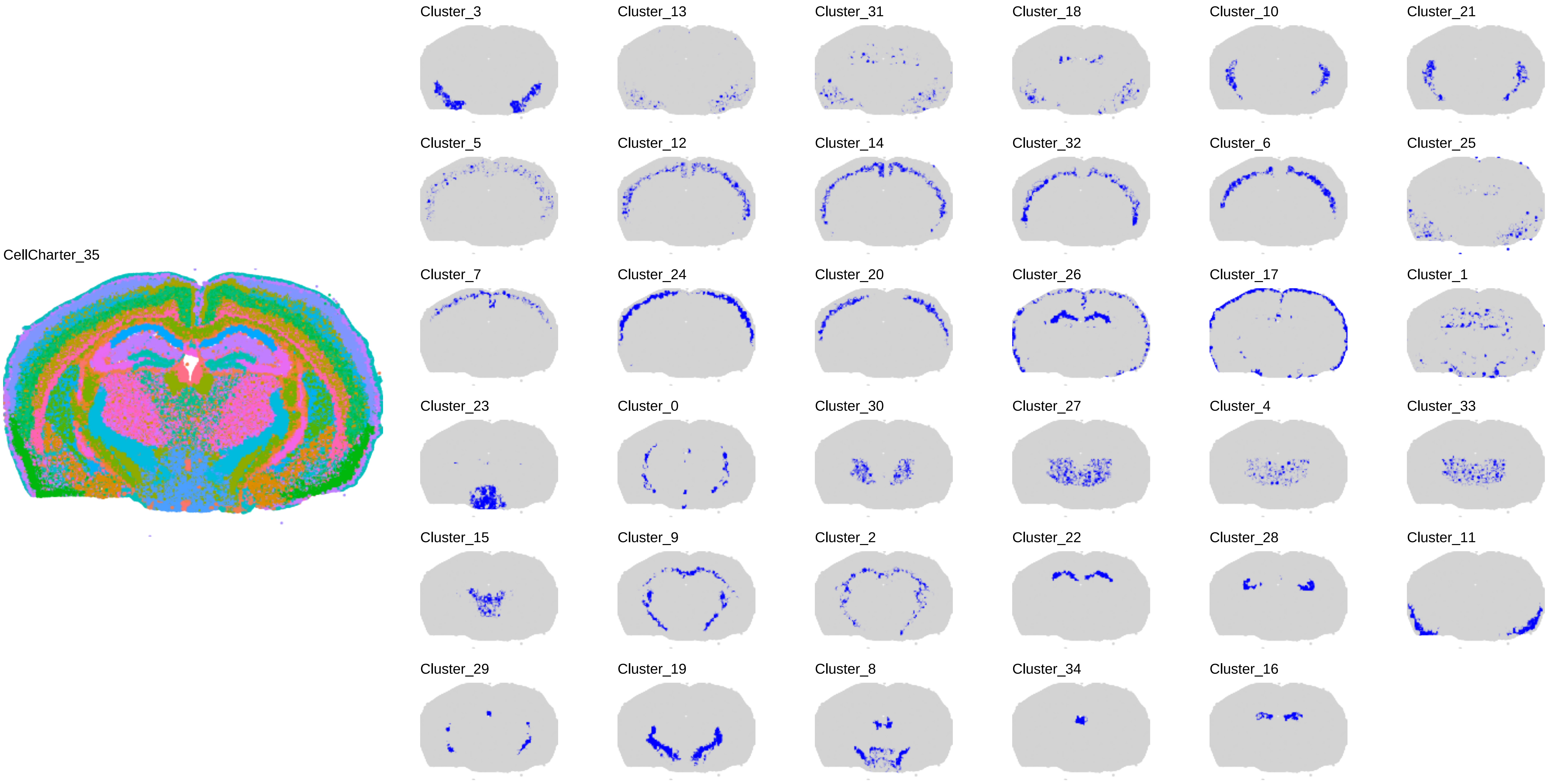
CellCharter_35
Cluster_3
Cluster_13
Cluster_31
Cluster_18
Cluster_10
Cluster_21
Cluster_5
Cluster_12
Cluster_14
Cluster_32
Cluster_6
Cluster_25
Cluster_7
Cluster_24
Cluster_20
Cluster_26
Cluster_17
Cluster_1
Cluster_23
Cluster_0
Cluster_30
Cluster_27
Cluster_4
Cluster_33
Cluster_15
Cluster_9
Cluster_2
Cluster_22
Cluster_28
Cluster_11
Cluster_29
Cluster_19
Cluster_8
Cluster_34
Cluster_16

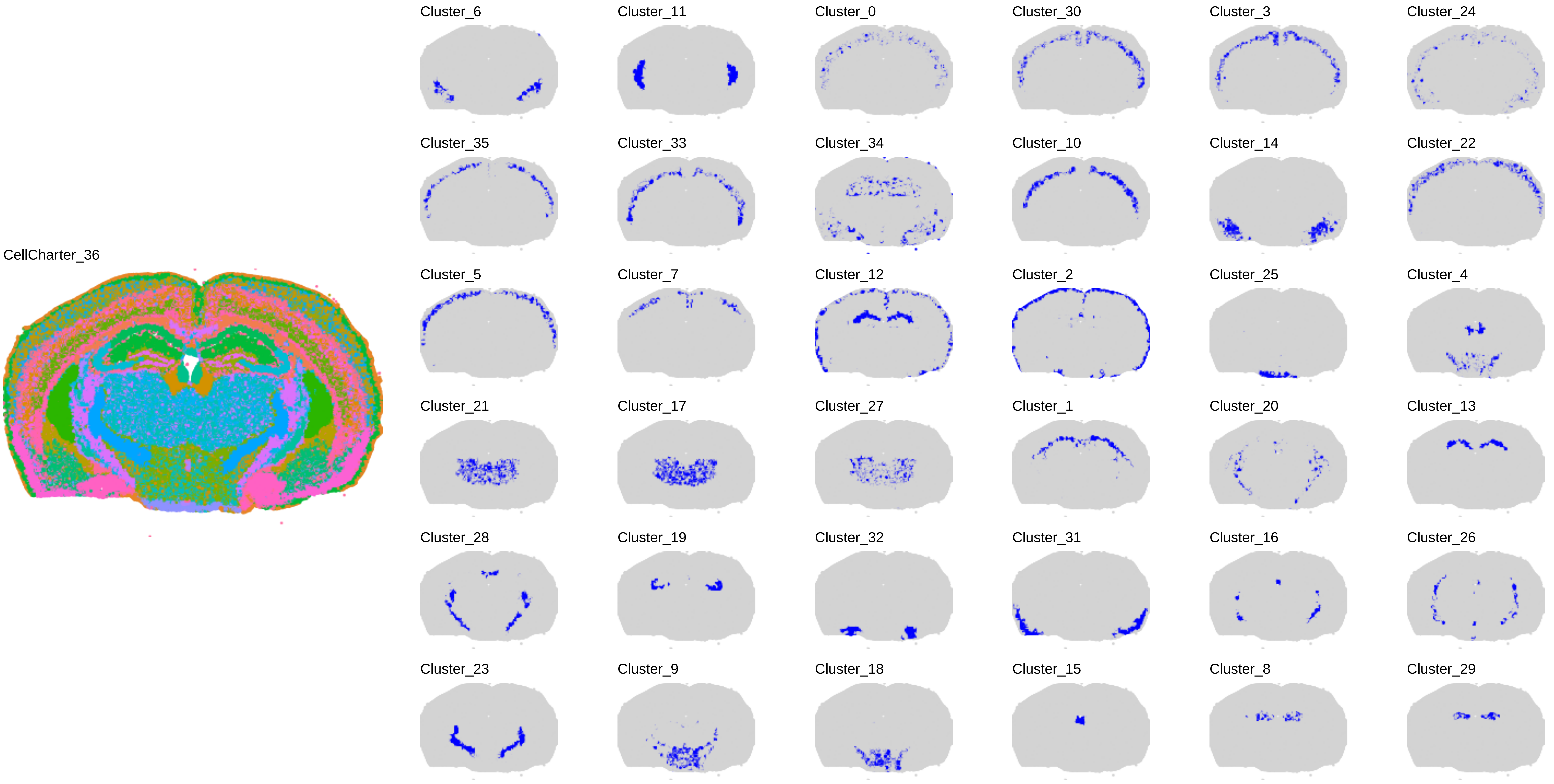
CellCharter_36
Cluster_6
Cluster_11
Cluster_0
Cluster_30
Cluster_3
Cluster_24
Cluster_35
Cluster_33
Cluster_34
Cluster_10
Cluster_14
Cluster_22
Cluster_5
Cluster_7
Cluster_12
Cluster_2
Cluster_25
Cluster_4
Cluster_21
Cluster_17
Cluster_27
Cluster_1
Cluster_20
Cluster_13
Cluster_28
Cluster_19
Cluster_32
Cluster_31
Cluster_16
Cluster_26
Cluster_23
Cluster_9
Cluster_18
Cluster_15
Cluster_8
Cluster_29

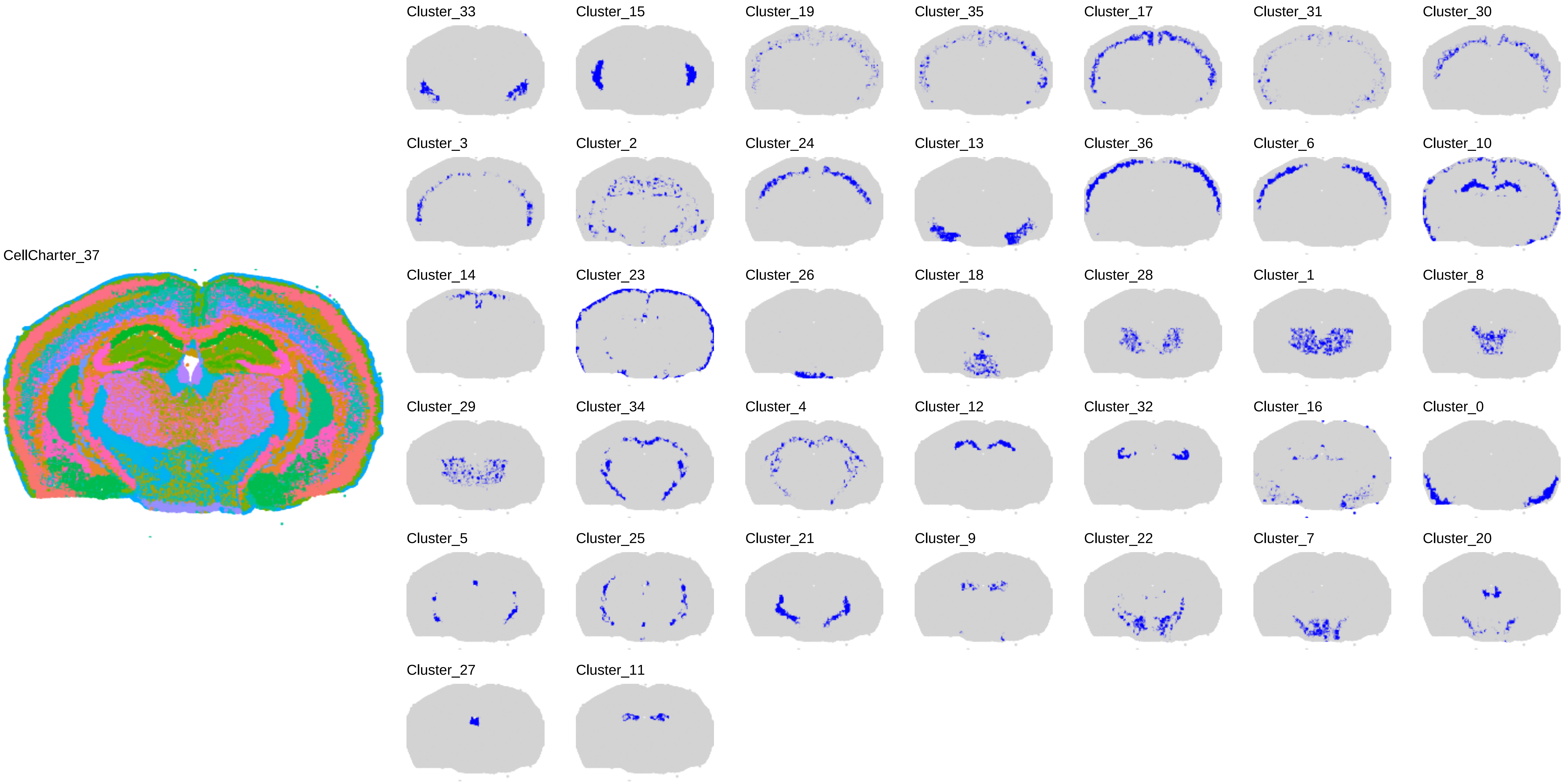
CellCharter_37
Cluster_33
Cluster_15
Cluster_19
Cluster_35
Cluster_17
Cluster_31
Cluster_30
Cluster_3
Cluster_2
Cluster_24
Cluster_13
Cluster_36
Cluster_6
Cluster_10
Cluster_14
Cluster_23
Cluster_26
Cluster_18
Cluster_28
Cluster_1
Cluster_8
Cluster_29
Cluster_34
Cluster_4
Cluster_12
Cluster_32
Cluster_16
Cluster_0
Cluster_5
Cluster_25
Cluster_21
Cluster_9
Cluster_22
Cluster_7
Cluster_20
Cluster_27
Cluster_11

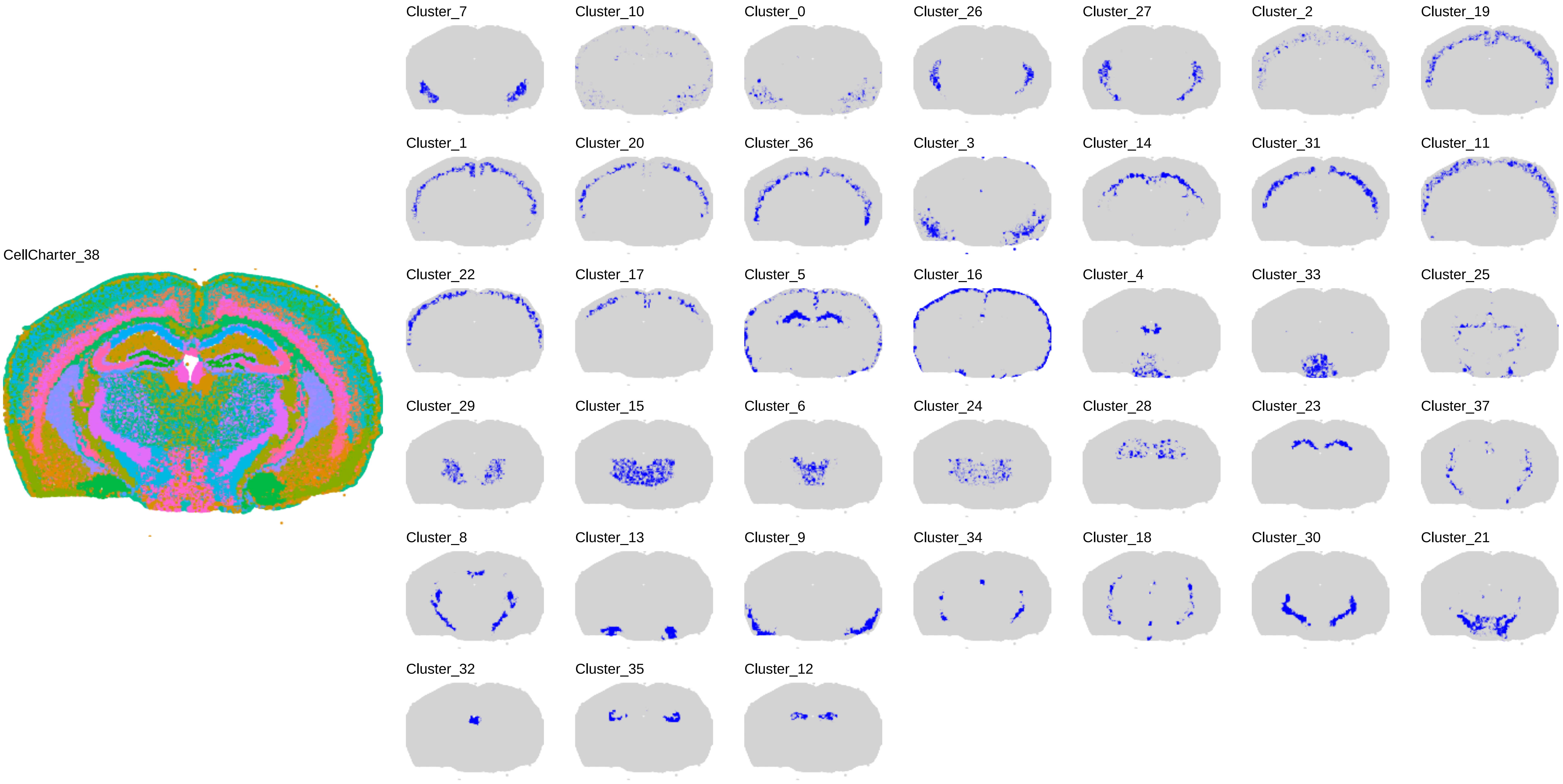
CellCharter_38
Cluster_7
Cluster_10
Cluster_0
Cluster_26
Cluster_27
Cluster_2
Cluster_19
Cluster_1
Cluster_20
Cluster_36
Cluster_3
Cluster_14
Cluster_31
Cluster_11
Cluster_22
Cluster_17
Cluster_5
Cluster_16
Cluster_4
Cluster_33
Cluster_25
Cluster_29
Cluster_15
Cluster_6
Cluster_24
Cluster_28
Cluster_23
Cluster_37
Cluster_8
Cluster_13
Cluster_9
Cluster_34
Cluster_18
Cluster_30
Cluster_21
Cluster_32
Cluster_35
Cluster_12

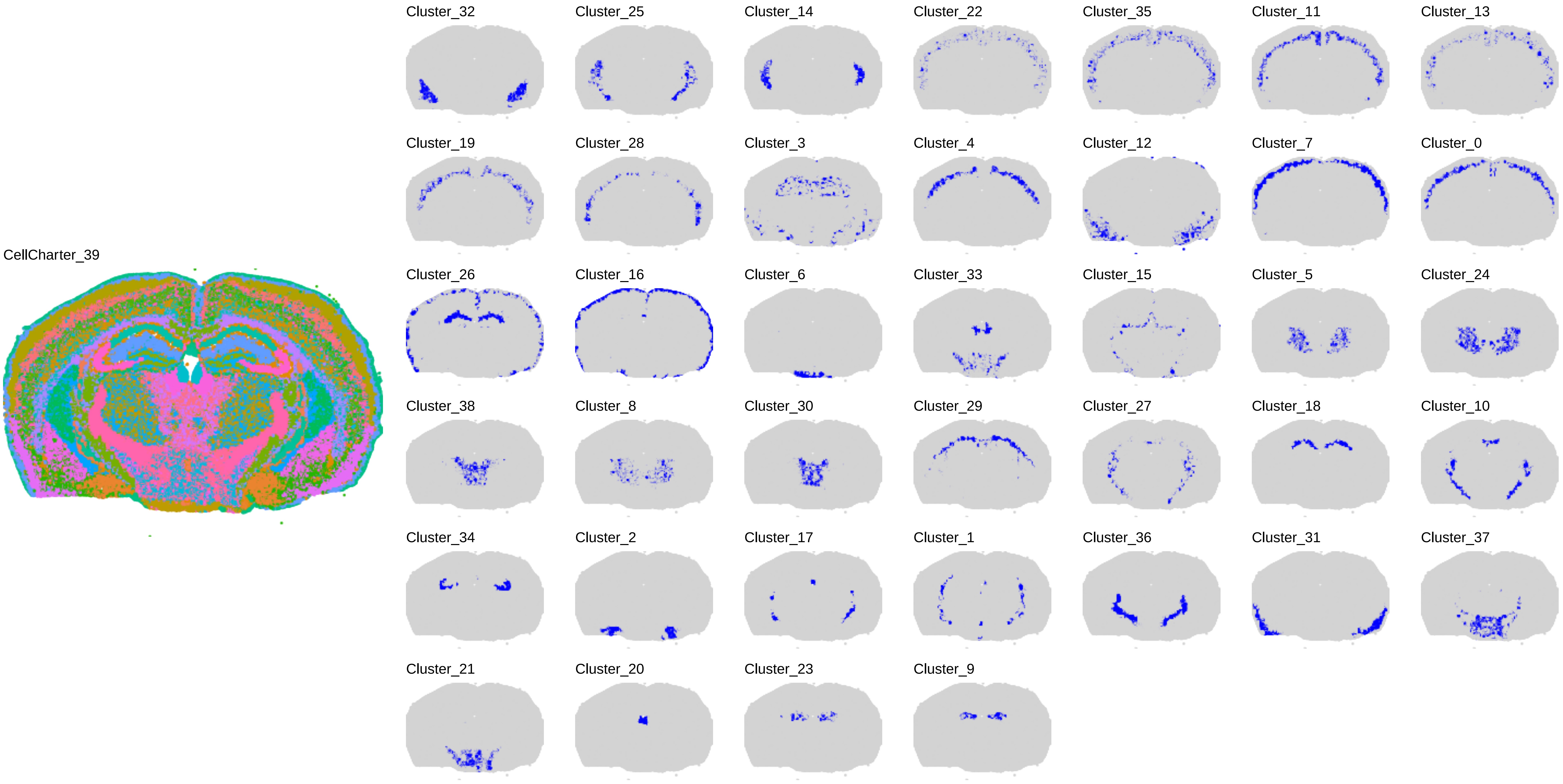
CellCharter_39
Cluster_32
Cluster_25
Cluster_14
Cluster_22
Cluster_35
Cluster_11
Cluster_13
Cluster_19
Cluster_28
Cluster_3
Cluster_4
Cluster_12
Cluster_7
Cluster_0
Cluster_26
Cluster_16
Cluster_6
Cluster_33
Cluster_15
Cluster_5
Cluster_24
Cluster_38
Cluster_8
Cluster_30
Cluster_29
Cluster_27
Cluster_18
Cluster_10
Cluster_34
Cluster_2
Cluster_17
Cluster_1
Cluster_36
Cluster_31
Cluster_37
Cluster_21
Cluster_20
Cluster_23
Cluster_9

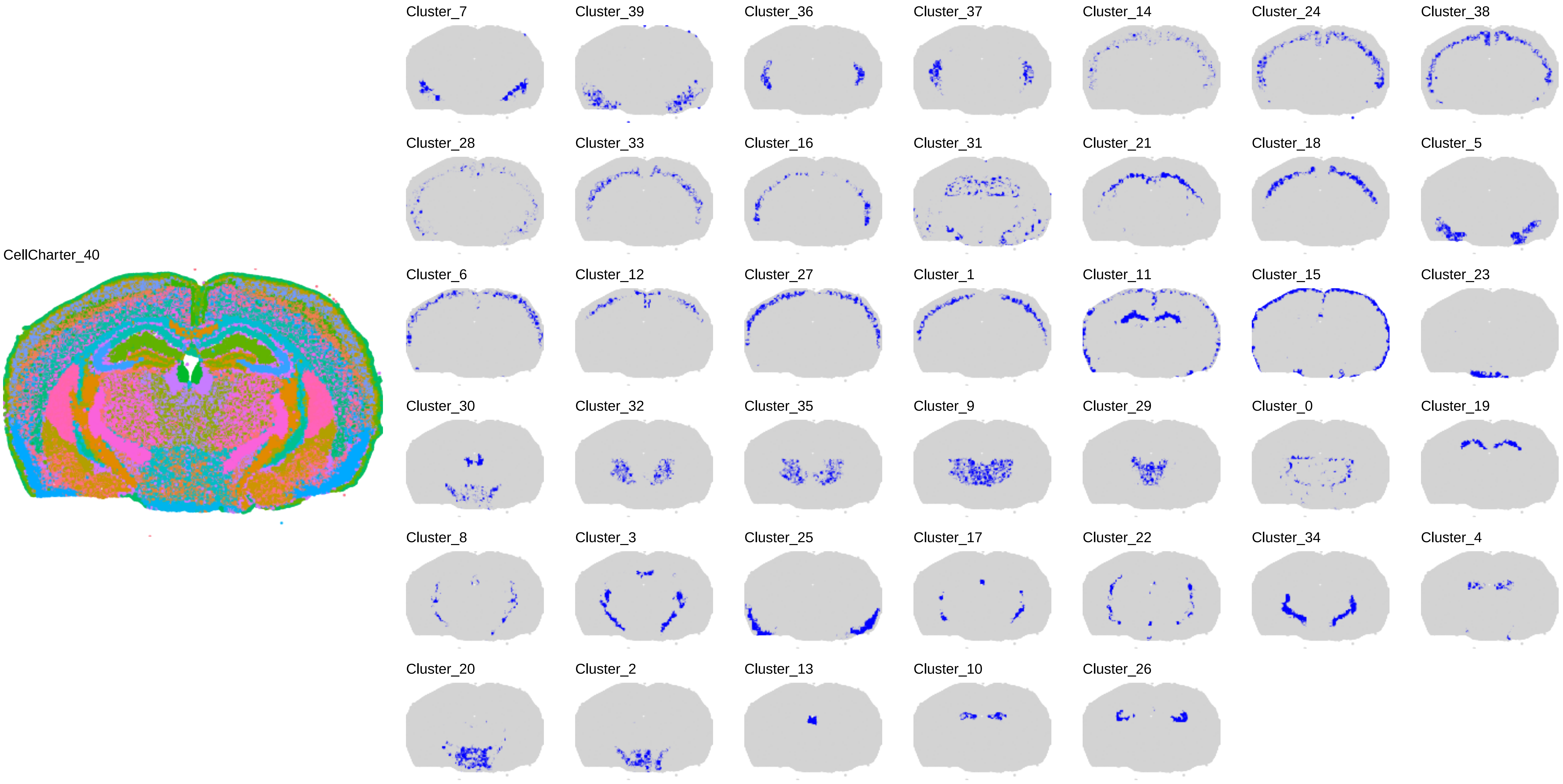
CellCharter_40
Cluster_7
Cluster_39
Cluster_36
Cluster_37
Cluster_14
Cluster_24
Cluster_38
Cluster_28
Cluster_33
Cluster_16
Cluster_31
Cluster_21
Cluster_18
Cluster_5
Cluster_6
Cluster_12
Cluster_27
Cluster_1
Cluster_11
Cluster_15
Cluster_23
Cluster_30
Cluster_32
Cluster_35
Cluster_9
Cluster_29
Cluster_0
Cluster_19
Cluster_8
Cluster_3
Cluster_25
Cluster_17
Cluster_22
Cluster_34
Cluster_4
Cluster_20
Cluster_2
Cluster_13
Cluster_10
Cluster_26

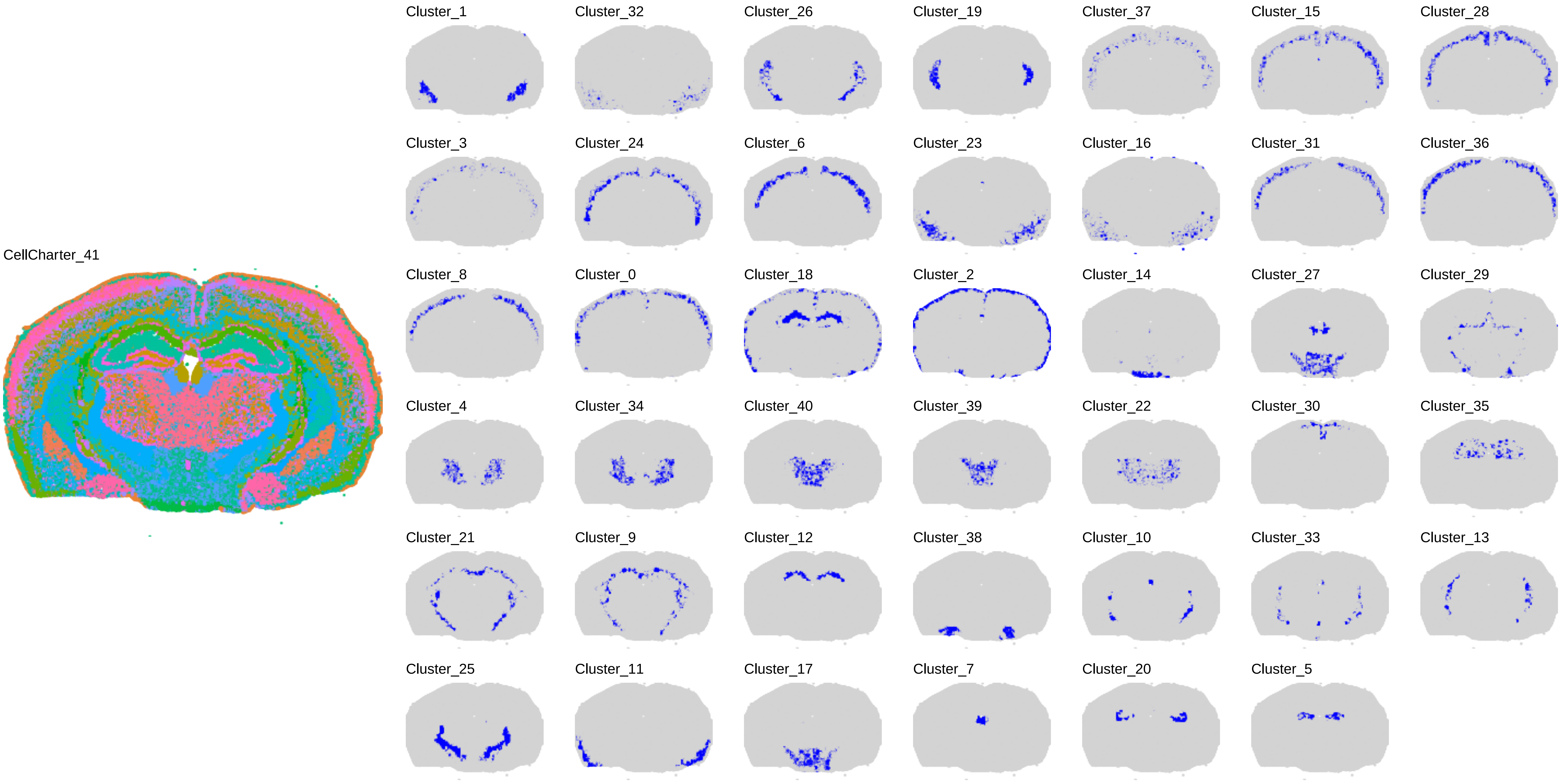

CellCharter_41
Cluster_1
Cluster_32
Cluster_26
Cluster_19
Cluster_37
Cluster_15
Cluster_28
Cluster_3
Cluster_24
Cluster_6
Cluster_23
Cluster_16
Cluster_31
Cluster_36
Cluster_8
Cluster_0
Cluster_18
Cluster_2
Cluster_14
Cluster_27
Cluster_29
Cluster_4
Cluster_34
Cluster_40
Cluster_39
Cluster_22
Cluster_30
Cluster_35
Cluster_21
Cluster_9
Cluster_12
Cluster_38
Cluster_10
Cluster_33
Cluster_13
Cluster_25
Cluster_11
Cluster_17
Cluster_7
Cluster_20
Cluster_5

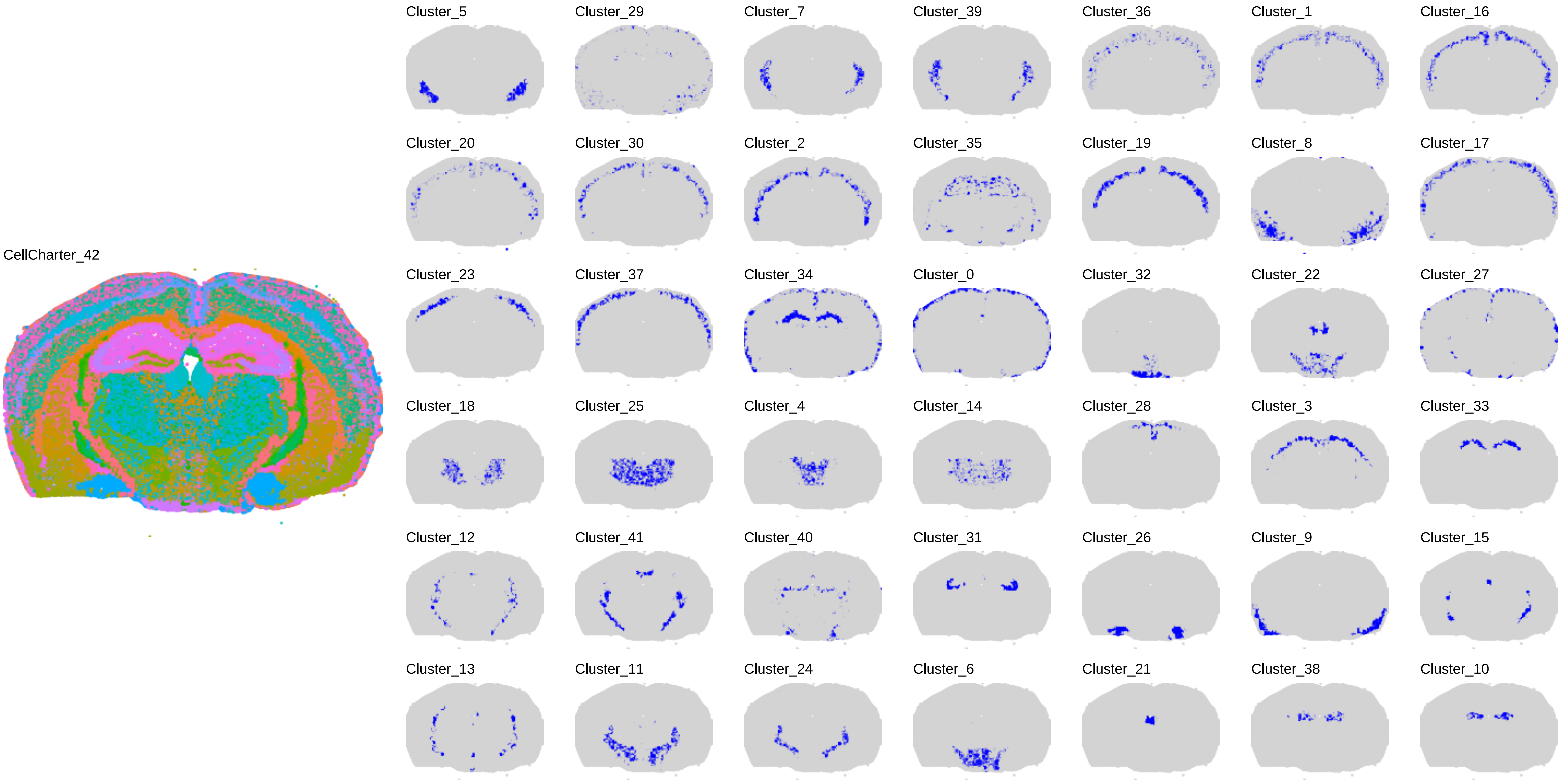
CellCharter_42
Cluster_5
Cluster_29
Cluster_7
Cluster_39
Cluster_36
Cluster_1
Cluster_16
Cluster_20
Cluster_30
Cluster_2
Cluster_35
Cluster_19
Cluster_8
Cluster_17
Cluster_23
Cluster_37
Cluster_34
Cluster_0
Cluster_32
Cluster_22
Cluster_27
Cluster_18
Cluster_25
Cluster_4
Cluster_14
Cluster_28
Cluster_3
Cluster_33
Cluster_12
Cluster_41
Cluster_40
Cluster_31
Cluster_26
Cluster_9
Cluster_15
Cluster_13
Cluster_11
Cluster_24
Cluster_6
Cluster_21
Cluster_38
Cluster_10

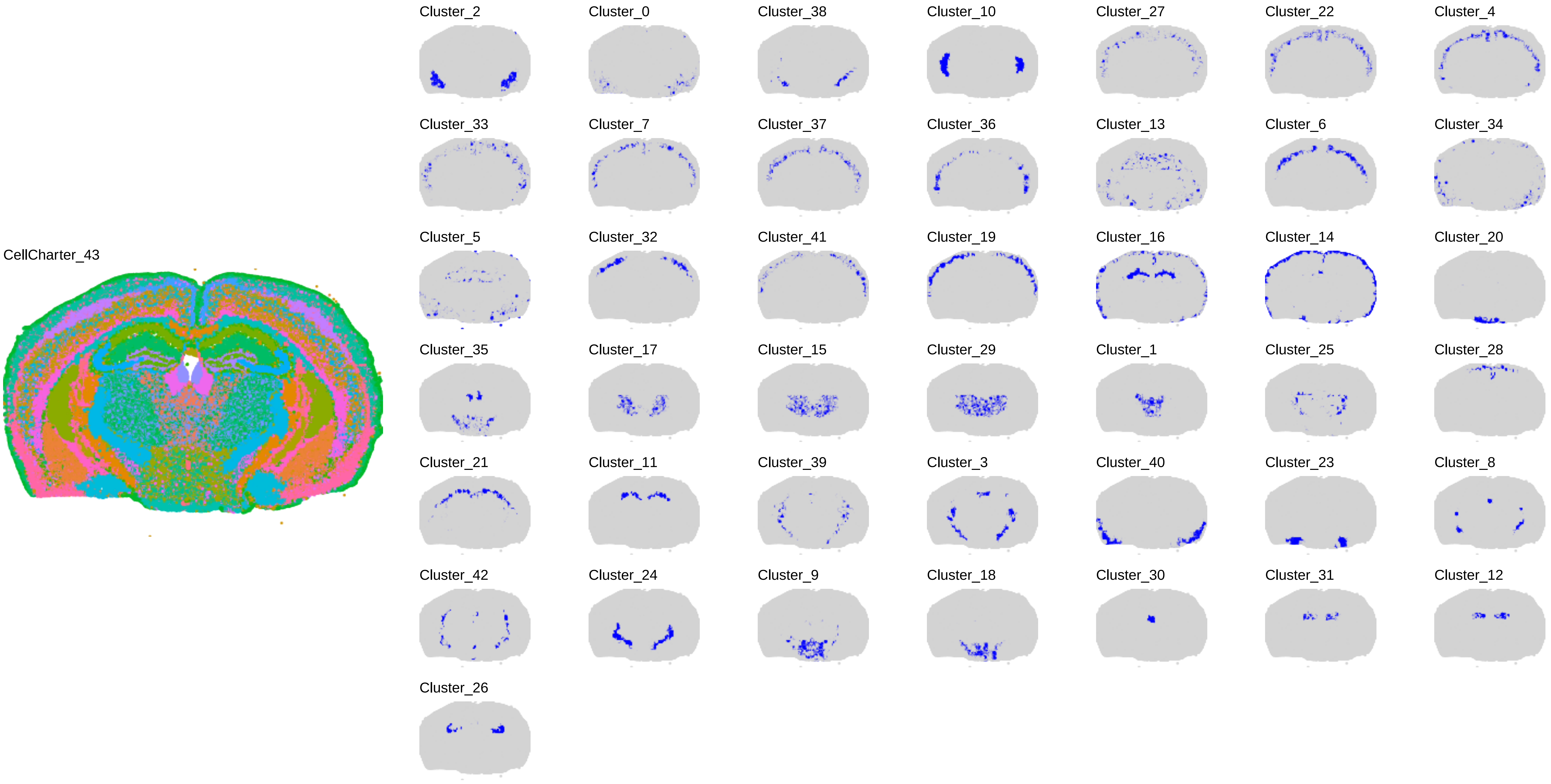

CellCharter_43
Cluster_2
Cluster_0
Cluster_38
Cluster_10
Cluster_27
Cluster_22
Cluster_4
Cluster_33
Cluster_7
Cluster_37
Cluster_36
Cluster_13
Cluster_6
Cluster_34
Cluster_5
Cluster_32
Cluster_41
Cluster_19
Cluster_16
Cluster_14
Cluster_20
Cluster_35
Cluster_17
Cluster_15
Cluster_29
Cluster_1
Cluster_25
Cluster_28
Cluster_21
Cluster_11
Cluster_39
Cluster_3
Cluster_40
Cluster_23
Cluster_8
Cluster_42
Cluster_24
Cluster_9
Cluster_18
Cluster_30
Cluster_31
Cluster_12
Cluster_26

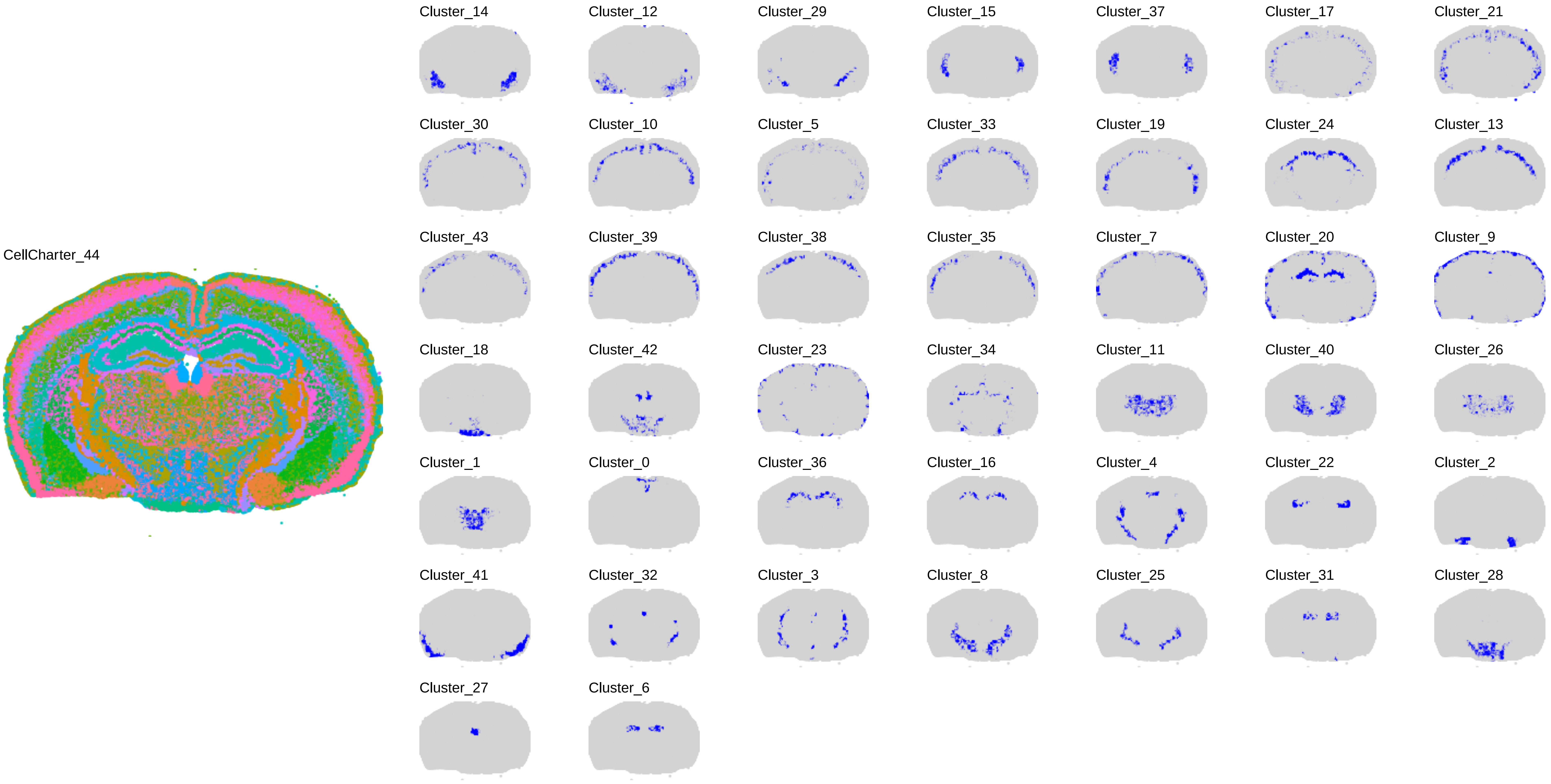
CellCharter_44
Cluster_14
Cluster_12
Cluster_29
Cluster_15
Cluster_37
Cluster_17
Cluster_21
Cluster_30
Cluster_10
Cluster_5
Cluster_33
Cluster_19
Cluster_24
Cluster_13
Cluster_43
Cluster_39
Cluster_38
Cluster_35
Cluster_7
Cluster_20
Cluster_9
Cluster_18
Cluster_42
Cluster_23
Cluster_34
Cluster_11
Cluster_40
Cluster_26
Cluster_1
Cluster_0
Cluster_36
Cluster_16
Cluster_4
Cluster_22
Cluster_2
Cluster_41
Cluster_32
Cluster_3
Cluster_8
Cluster_25
Cluster_31
Cluster_28
Cluster_27
Cluster_6

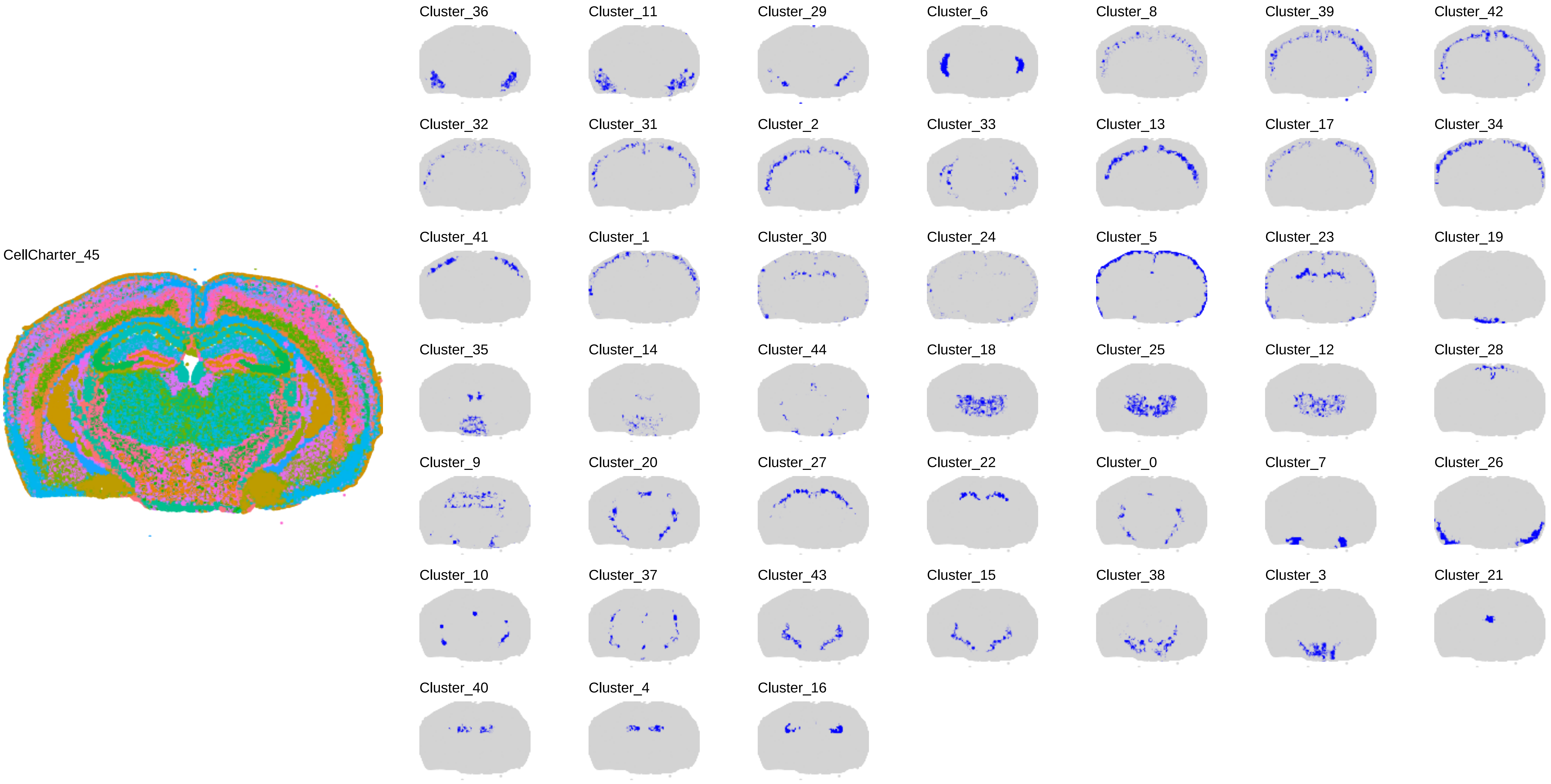
CellCharter_45
Cluster_36
Cluster_11
Cluster_29
Cluster_6
Cluster_8
Cluster_39
Cluster_42
Cluster_32
Cluster_31
Cluster_2
Cluster_33
Cluster_13
Cluster_17
Cluster_34
Cluster_41
Cluster_1
Cluster_30
Cluster_24
Cluster_5
Cluster_23
Cluster_19
Cluster_35
Cluster_14
Cluster_44
Cluster_18
Cluster_25
Cluster_12
Cluster_28
Cluster_9
Cluster_20
Cluster_27
Cluster_22
Cluster_0
Cluster_7
Cluster_26
Cluster_10
Cluster_37
Cluster_43
Cluster_15
Cluster_38
Cluster_3
Cluster_21
Cluster_40
Cluster_4
Cluster_16

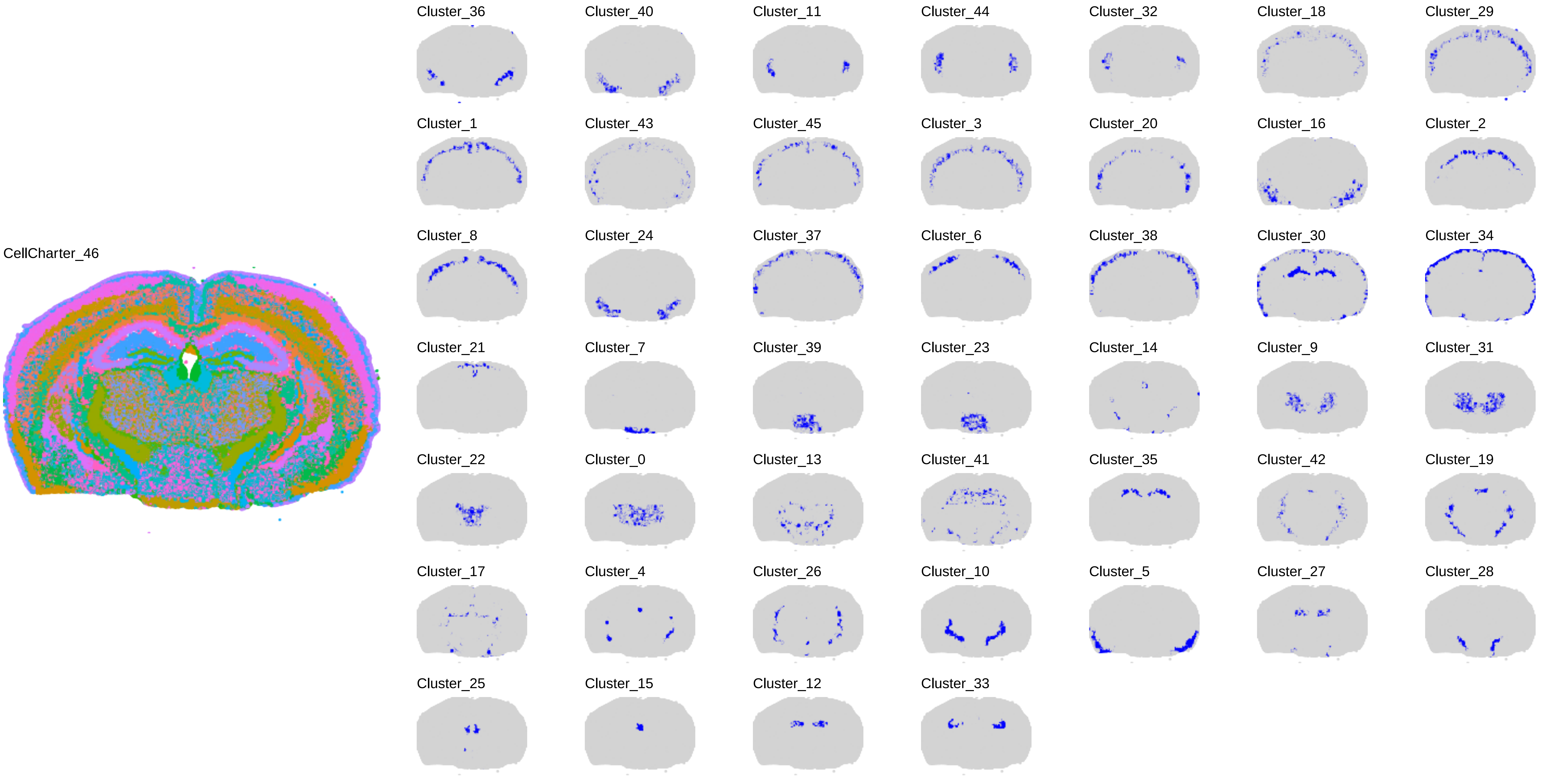

CellCharter_46
Cluster_36
Cluster_40
Cluster_11
Cluster_44
Cluster_32
Cluster_18
Cluster_29
Cluster_1
Cluster_43
Cluster_45
Cluster_3
Cluster_20
Cluster_16
Cluster_2
Cluster_8
Cluster_24
Cluster_37
Cluster_6
Cluster_38
Cluster_30
Cluster_34
Cluster_21
Cluster_7
Cluster_39
Cluster_23
Cluster_14
Cluster_9
Cluster_31
Cluster_22
Cluster_0
Cluster_13
Cluster_41
Cluster_35
Cluster_42
Cluster_19
Cluster_17
Cluster_4
Cluster_26
Cluster_10
Cluster_5
Cluster_27
Cluster_28
Cluster_25
Cluster_15
Cluster_12
Cluster_33

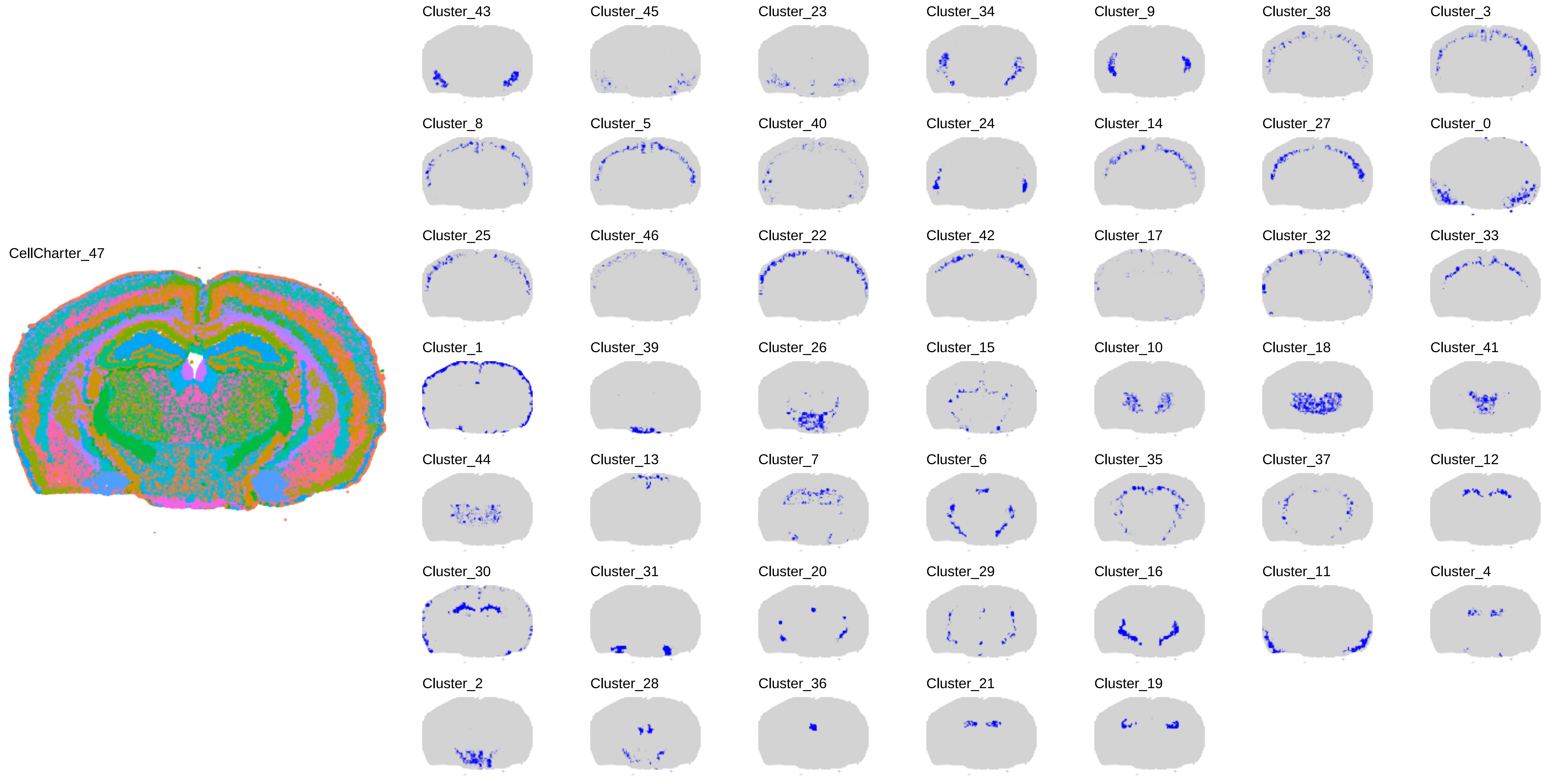
CellCharter_47
Cluster_43
Cluster_45
Cluster_23
Cluster_34
Cluster_9
Cluster_38
Cluster_3
Cluster_8
Cluster_5
Cluster_40
Cluster_24
Cluster_14
Cluster_27
Cluster_0
Cluster_25
Cluster_46
Cluster_22
Cluster_42
Cluster_17
Cluster_32
Cluster_33
Cluster_1
Cluster_39
Cluster_26
Cluster_15
Cluster_10
Cluster_18
Cluster_41
Cluster_44
Cluster_13
Cluster_7
Cluster_6
Cluster_35
Cluster_37
Cluster_12
Cluster_30
Cluster_31
Cluster_20
Cluster_29
Cluster_16
Cluster_11
Cluster_4
Cluster_2
Cluster_28
Cluster_36
Cluster_21
Cluster_19

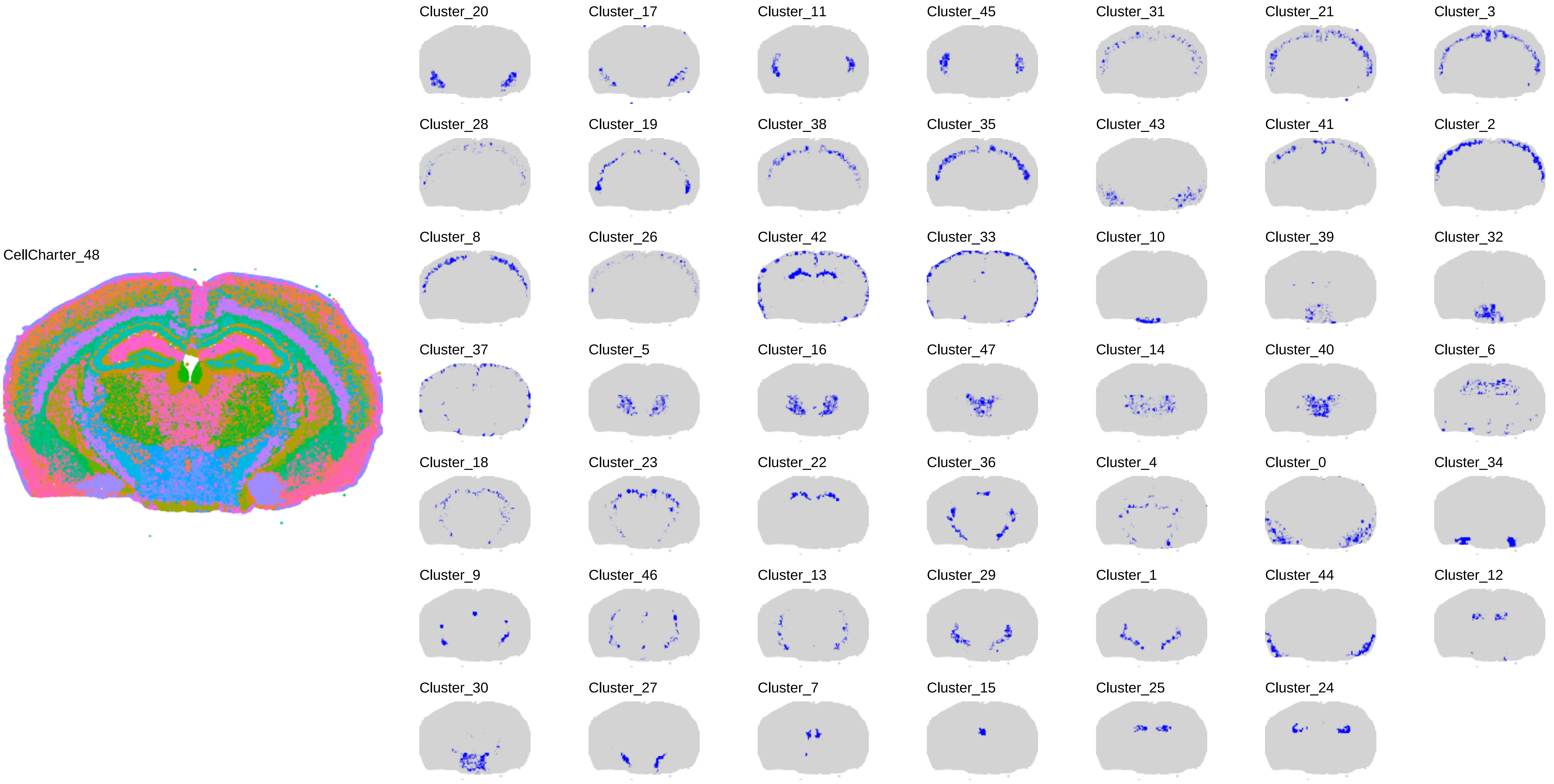
CellCharter_48
Cluster_20
Cluster_17
Cluster_11
Cluster_45
Cluster_31
Cluster_21
Cluster_3
Cluster_28
Cluster_19
Cluster_38
Cluster_35
Cluster_43
Cluster_41
Cluster_2
Cluster_8
Cluster_26
Cluster_42
Cluster_33
Cluster_10
Cluster_39
Cluster_32
Cluster_37
Cluster_5
Cluster_16
Cluster_47
Cluster_14
Cluster_40
Cluster_6
Cluster_18
Cluster_23
Cluster_22
Cluster_36
Cluster_4
Cluster_0
Cluster_34
Cluster_9
Cluster_46
Cluster_13
Cluster_29
Cluster_1
Cluster_44
Cluster_12
Cluster_30
Cluster_27
Cluster_7
Cluster_15
Cluster_25
Cluster_24

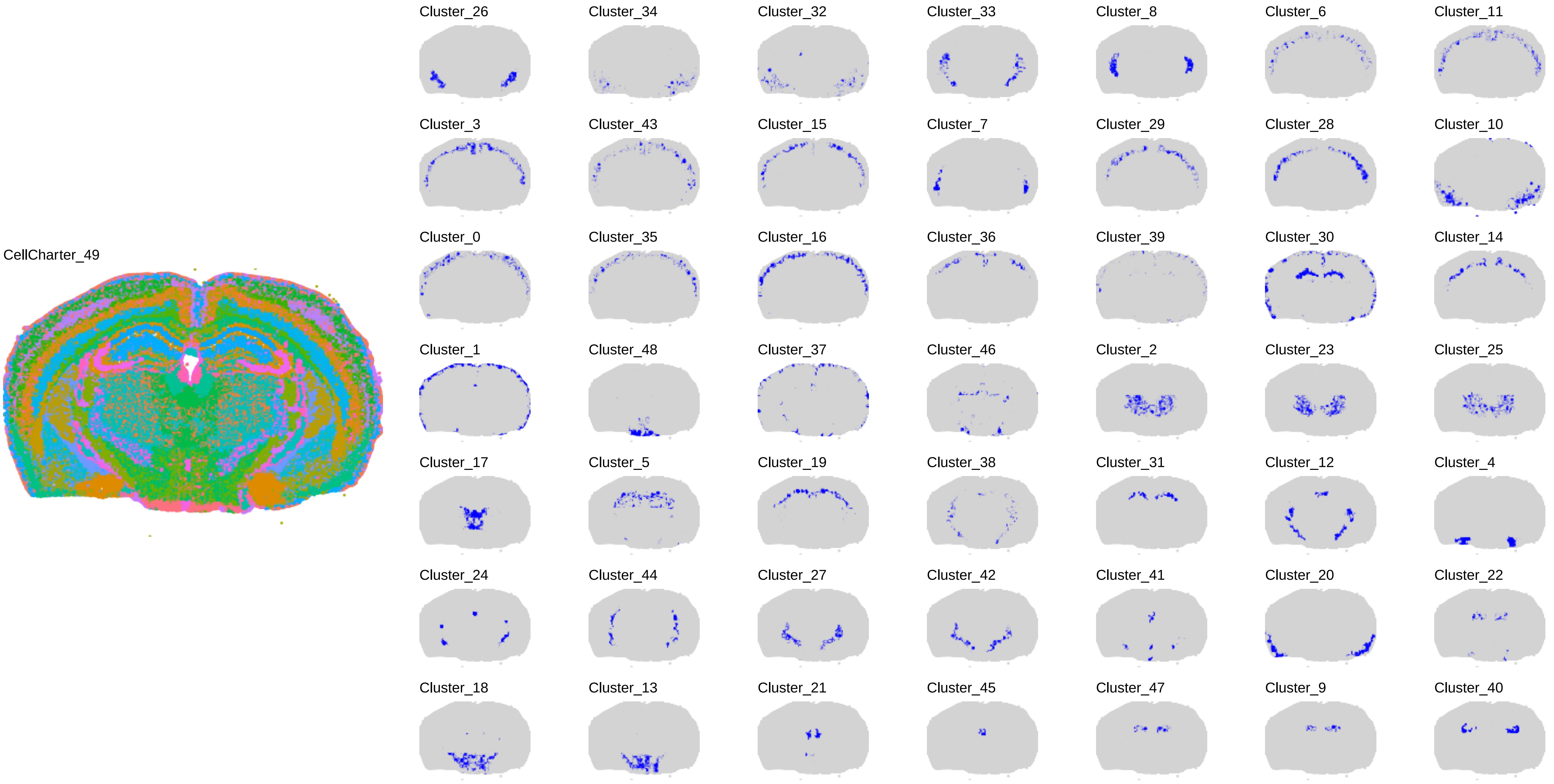
CellCharter_49
Cluster_26
Cluster_34
Cluster_32
Cluster_33
Cluster_8
Cluster_6
Cluster_11
Cluster_3
Cluster_43
Cluster_15
Cluster_7
Cluster_29
Cluster_28
Cluster_10
Cluster_0
Cluster_35
Cluster_16
Cluster_36
Cluster_39
Cluster_30
Cluster_14
Cluster_1
Cluster_48
Cluster_37
Cluster_46
Cluster_2
Cluster_23
Cluster_25
Cluster_17
Cluster_5
Cluster_19
Cluster_38
Cluster_31
Cluster_12
Cluster_4
Cluster_24
Cluster_44
Cluster_27
Cluster_42
Cluster_41
Cluster_20
Cluster_22
Cluster_18
Cluster_13
Cluster_21
Cluster_45
Cluster_47
Cluster_9
Cluster_40
